# Systematic Determination of the Impact of Structural Edits on Peptide Accumulation into Mycobacteria

**DOI:** 10.1101/2025.01.17.633618

**Authors:** Rachita Dash, Zichen Liu, Irene Lepori, Mahendra D. Chordia, Karl Ocius, Kadie Holsinger, Han Zhang, Ryan Kenyon, Wonpil Im, M. Sloan Siegrist, Marcos M. Pires

**Affiliations:** Department of Chemistry; Department of Microbiology, Immunology, and Cancer University of Virginia Charlottesville, VA, United States; Molecular and Cellular Biology Graduate Program; Department of Microbiology University of Massachusetts Amherst, MA, United States; Department of Biological Sciences Lehigh University Bethlehem, PA, United States

## Abstract

Understanding the factors that influence the accumulation of molecules beyond the mycomembrane of *Mycobacterium tuberculosis* (*Mtb*) – the main barrier to accumulation – is essential for developing effective antimycobacterial agents. In this study, we investigated two design principles commonly observed in natural products and mammalian cell-permeable peptides: backbone *N*-alkylation and macrocyclization. To assess how these structural edits impact molecule accumulation beyond the mycomembrane, we utilized our recently developed Peptidoglycan Accessibility Click-Mediated Assessment (PAC-MAN) assay for live-cell analysis. Our findings provide the first empirical evidence that peptide macrocyclization generally enhances accumulation in mycobacteria, while *N*-alkylation influences accumulation in a context-dependent manner. We examined these design principles in the context of two peptide antibiotics, tridecaptin A1 and griselimycin, which revealed the roles of *N*-alkylation and macrocyclization in improving both accumulation and antimicrobial activity against mycobacteria in specific contexts. Together, we present a working model for strategic structural modifications aimed at enhancing the accumulation of molecules past the mycomembrane. More broadly, our results also challenge the prevailing belief in the field that large and hydrophilic molecules, such as peptides, cannot readily traverse the mycomembrane.

## MAIN

Tuberculosis (TB) is a major global public health concern, with over 10 million cases reported worldwide in 2022.^1^ Often recognized as the deadliest infectious disease, TB has recently seen a surge in incidence, reversing a decade-long trend of decline.^2^ The causative agent of TB is *Mycobacterium tuberculosis* (*Mtb)*. *Mtb* and other mycobacteria feature a complex cellular envelope that includes an outer mycomembrane, an arabinogalactan layer, a peptidoglycan (PG) layer, and an inner membrane.^3^ The mycomembrane is uniquely thick and hydrophobic, primarily composed of long-chain lipids (mycolic acids) and trehalose-based glycolipids.^3^ This mycomembrane is widely regarded as the primary permeation barrier to the entry of antibiotics, providing mycobacteria with a high level of intrinsic resistance.^4–7^

Most antibiotics currently used in the treatment of TB, such as isoniazid, ethambutol, and pyrazinamide, are small, hydrophobic molecules developed several decades ago.^8^ These restrictive molecular features of TB-active antibiotics are thought to result directly from the challenge of crossing the mycomembrane barrier. The overuse of these therapeutics, compounded by suboptimal patient adherence due to prolonged treatment regimens, has led to the emergence of multi-drug resistant (MDR) and total drug-resistant (XDR) strains of *Mtb*.^9^ In recent years, only two new antibiotics, bedaquiline and pretomanid (both small and hydrophobic), have been introduced to treat drug-resistant tuberculosis, underscoring the urgent need to expand the antimycobacterial pipeline.^10^

There are notable exceptions to the size constraints typically associated with anti-TB antibiotics (such as rifampicin), suggesting that larger hydrophilic molecules, including peptides, could be viable candidates for antimycobacterial therapy. Peptidic molecules that exceed Lipinski’s Rule of Five (Ro5)^11^ have garnered considerable interest in drug design^12–14^ across various disease areas (e.g., oncology^15^ and metabolic^16,17^ disorders) due to their ability to bind targets with greater specificity and affinity. A significant example is the development of a new class of macrocyclic peptide antibiotics, Zosurabalpin (developed by Roche), which specifically targets the lipopolysaccharide transporter in carbapenem-resistant *Acinetobacter baumannii.*^18^ Additionally, peptide candidates are actively being developed for mycobacterial infections.^19,20^ This growing interest is exemplified by the recent discoveries of evybactin^21^ and cyclomarin A^22^ (**Fig. 1a**), which have served as the basis for a promising series of BacPROTACs.^23–26^

**Fig. 1.**
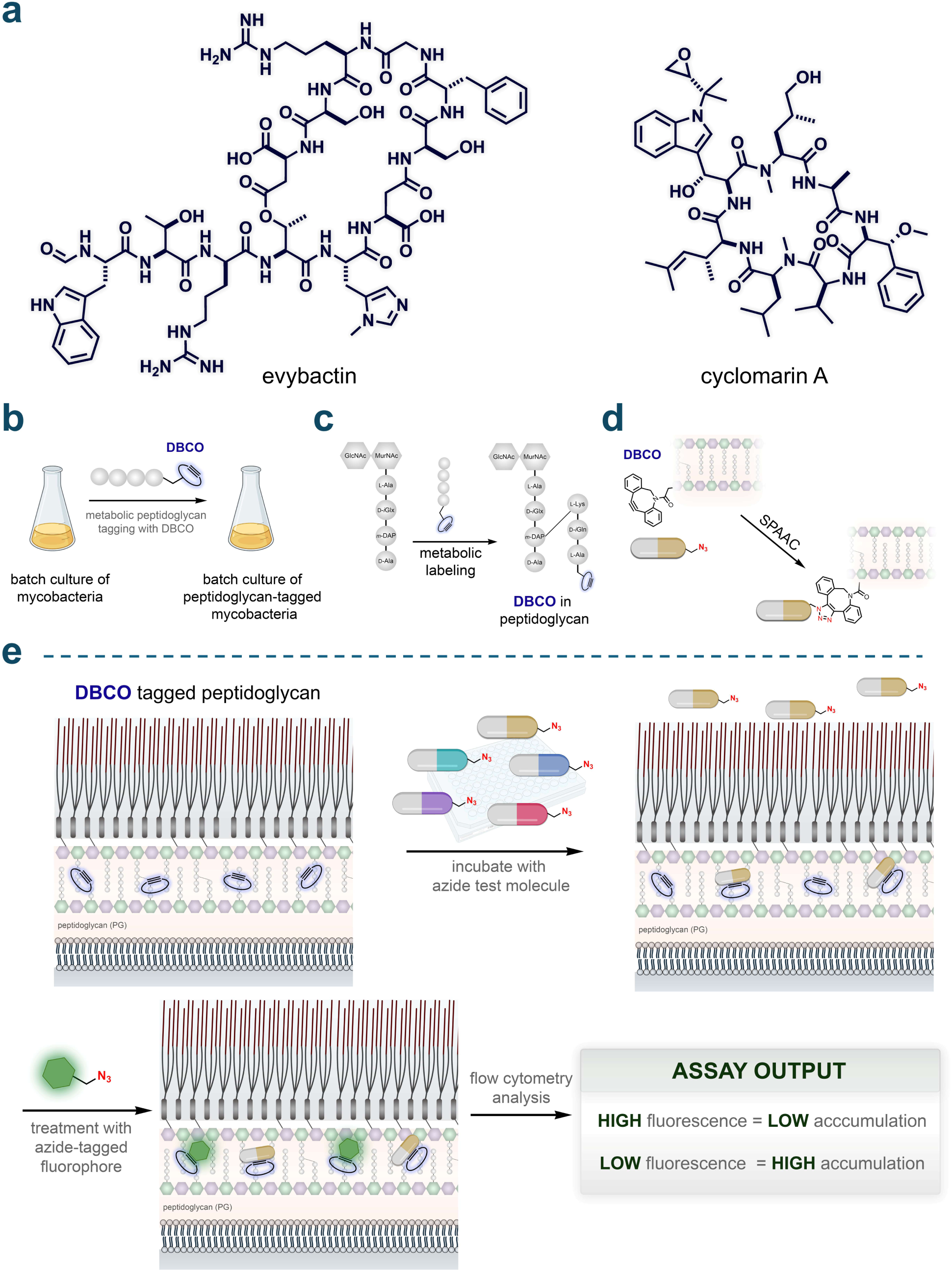
**(a)** Chemical structures of the peptides evybactin and cyclomarin A. **(b)** General schematic illustrating the labeling of mycobacteria with DBCO. Mycobacteria were grown in batches and metabolically labeled with DBCO using a modified exogenous stem peptide probe. **(c)** Schematic depicting site-specific tagging of the peptidoglycan layer with DBCO. An exogenous stem peptide modified with a DBCO moiety on its *N*-terminus is crosslinked onto the existing peptidoglycan framework. This occurs *via* the installation of a covalent link between the meso-α,ε-diaminopimelic acid (m-DAP) residue on a stem peptide of the existing peptidoglycan framework and the Lys residue on the exogenously added stem peptide, by transpeptidases. **(d)** Schematic illustration of strain-promoted azide-alkyne cycloaddition (SPAAC) occurring between the azide-tagged test molecules and the installed DBCO landmarks within the PG. **(e)** Schematic illustration of PAC-MAN. Incubation of azide-tagged molecules with the DBCO-labeled mycobacteria allows the registration of their arrival past the mycomembrane onto the PG through SPAAC. A subsequent incubation step with an azide-tagged fluorophore in a secondary SPAAC reveals the unoccupied DBCO landmarks. Cellular fluorescence is then measured by flow cytometry where high fluorescence values correspond to low accumulation and vice versa.

Backbone *N*-alkylation and macrocyclization are common modifications observed in natural peptide products.^27–32^ These structural changes have been extensively leveraged to enhance the passive membrane permeability of peptides in mammalian systems.^33–37^ However, these strategies have not yet been empirically validated in mycobacteria. A significant challenge in this field is the general lack of tools to readily measure the accumulation of molecules past the mycomembrane. In our previous work, we developed the Peptidoglycan Accessibility Click-Mediated Assessment (PAC-MAN) assay for live-cell analysis, demonstrating its capability to assess the accumulation of small molecules^38^ and antibiotics^7^ across the mycomembrane. Here, we expand upon this work to systematically evaluate the potential of backbone *N*-alkylation and macrocyclization in peptides to enhance their molecular accumulation levels in live mycobacteria. Through a comprehensive series of peptides, we demonstrate that these structural features can significantly influence accumulation levels. This discovery lays the groundwork for a set of prescriptive modifications that can be employed in developing more effective antibiotics targeting mycobacteria.

## RESULTS

### Establishing PAC-MAN Assay Parameters

Measuring molecular accumulation in bacteria presents notable technical challenges and remains an area of ongoing research in our laboratory.^38–42^ Liquid chromatography-tandem mass spectrometry (LC-MS/MS) is considered the gold standard for quantifying whole-cell associated molecule concentrations in bacteria^43–46^ without the need for chemical tags. However, there are two principal limitations in its current application: (a) limited throughput capacity for conventional setups,^45^ and (b) a lack of precision in identifying the exact subcellular location of molecules, as it typically measures ‘whole-cell’ colocalization or association.^44,47,48^ Despite these challenges, recent efforts in the Hergenrother laboratory have led to the development of the eNTRy rules in *Escherichia coli*^49^ and the PASsagE rules in *Pseudomonas aeruginosa*^50^ using LC-MS/MS. In contrast, to the best of our knowledge, large-scale LC-MS/MS analyses in *Mtb* have not been previously described; the largest screen of molecule accumulation conducted prior to the development of PAC-MAN involved a series of ten sulfonyladenosines.^51^ Critically, PAC-MAN registers the arrival of compounds in the periplasmic space once they have passed the mycomembrane and reached the peptidoglycan (PG) layer.

We previously demonstrated that the PG layer of live mycobacteria can be metabolically labeled with fluorophore-tagged PG analogs.^52–54^ In a variation of this labeling strategy for PAC-MAN analysis, the fluorophore is replaced with a strained alkyne dibenzocyclooctyne (DBCO) unit (**Fig. 1b**). When mycobacterial cells are treated with DBCO-tagged PG analogs, transpeptidases incorporate these analogs into the existing PG matrix through crosslinking, creating a distinct, site-specific biorthogonal chemical landmark (**Fig. 1c**).^38,52^ The incubation of azide test molecules with the DBCO-tagged mycobacterial cells enables the detection of their arrival past the mycomembrane *via* a strain-promoted alkyne-azide cycloaddition (SPAAC) reaction (**Fig. 1d**). Subsequent treatment with an azide-tagged fluorophore reveals the unoccupied DBCO landmarks, making cellular fluorescence inversely proportional to the level of molecule accumulation (**Fig. 1e**). With this setup, accumulation profiles of hundreds of molecules can be obtained in a 96-well format in parallel within a single experiment. More importantly, the arrival of the molecules past the most critical barrier to entry is confirmed by covalent SPAAC reactions.

Initially, we aimed to build on our previous assay parameters using *Mycobacterium smegmatis* (*Msm*), a model organism that recapitulates key features of the pathogenic *Mtb*.^55^ Live cell PG labeling was initiated by incubating *Msm* cells with **TetD**, a PG analog featuring DBCO on the *N*-terminus of the stem peptide (**Fig. 2a**). Two azide-tagged peptides were assembled to establish parameters for the PAC-MAN assay in the context of peptides, which served as the baseline members of a backbone *N*-alkylated sub-library (**Nmet0**) (**Fig. 2b**) and a linear sub-library (**Lin0**) (**Fig. S1**). Phenylalanine residues were included to obtain definitive concentrations based on the UV absorbance of the chromophore, and lysine residues were incorporated to enhance water solubility.

**Fig. 2.**
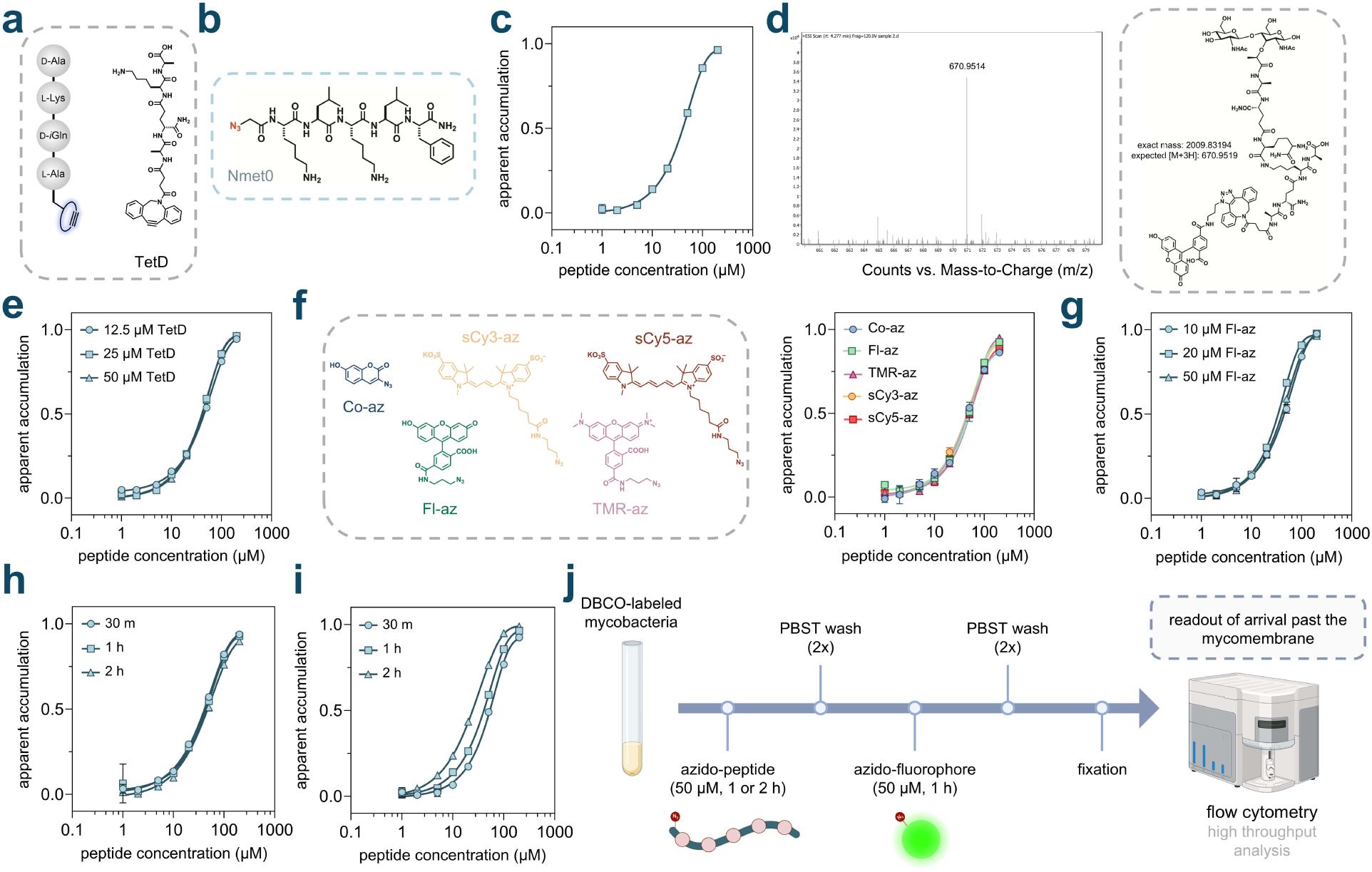
**(a)** Schematic illustration and chemical structure of **TetD**. **(b)** Chemical structure of the peptide **Nmet0**. **(c)** Dose-response analysis showing the apparent accumulation of **Nmet0** across the mycomembrane in *Msm*. **(d)** Mass spectrometry analysis of sacculi fragments isolated from **TetD** treated cells that were subsequently treated with the dye **Fl-az**. **(e)** Dose-response analysis showing the apparent accumulation of **Nmet0** across the mycomembrane in *Msm* labeled with varying concentrations of **TetD**. **(f)** Dose-response analysis showing the apparent accumulation of **Nmet0** across the mycomembrane in *Msm* treated with different azide-tagged fluorophores, and their chemical structures. **(g)** Dose-response analysis showing the apparent accumulation of **Nmet0** across the mycomembrane in *Msm* treated with varying concentrations of **Fl-az**. **(h)** Dose-response analysis showing the apparent accumulation of **Nmet0** across the mycomembrane in *Msm* with varying **Fl-az** treatment durations. **(i)** Dose-response analysis showing the apparent accumulation of **Nmet0** across the mycomembrane in *Msm* with varying **Nmet0** treatment durations. **(j)** Schematic illustration of optimized assay conditions. For the rest of experiments, **TetD** labeled mycobacterial cells were incubated with 50 μM azide-peptides for 1 h or 2 h and washed with PBST (phosphate buffered saline with 0.05% tween80) twice. Cells were then treated with 50 μM azide-tagged fluorophore for 1 h and washed with PBST twice and fixed in 4% formaldehyde (in PBS) before high throughput analysis on flow cytometry. All dose-response analyses involving **Nmet0** were performed with a 1 h incubation period with the peptide. Data are represented as mean ± SD (n= 3). For dose-response curves, Boltzmann sigmoidal curves were fitted to the data using GraphPad Prism.

In each assay, fluorescence intensities from cells without DBCO landmarks were considered as 0%, while fluorescence intensities from DBCO-tagged cells treated only with the azide-tagged fluorophore were regarded as 100% for the normalization of raw data. To facilitate visual interpretation, the data were processed as “1 − (normalized fluorescence intensity)”, a term referred to as ‘apparent accumulation’, which positively correlates with the accumulation of molecules. We began by testing varying concentrations of **Nmet0** *via* the PAC-MAN assay. Our results showed a clear concentration-dependent increase in apparent accumulation, indicating that the peptide **Nmet0** was able to arrive within the periplasmic space of *Msm* cells (**Fig. 2c**). The shape of the fitted curve is indeed similar to the concentration-dependent curves generated by the Chloroalkane Penetration Assay (CAPA), developed in the Kritzer lab for measuring penetration into mammalian cells.^56^ Given the size and hydrophilicity of this peptide, and considering the physicochemical properties of most antimycobacterial agents, the extent of its apparent accumulation is notable. The prevailing understanding in the field suggests that this type of peptide would likely show minimal accumulation past the mycomembrane.

A series of additional experiments was conducted to establish key aspects of the PAC-MAN assay. To confirm the installation of DBCO within the PG scaffold, sacculi were isolated from **TetD**-treated cells after treatment with the dye **Fl-az**. This isolation was performed using our previously reported procedures, which involved enzymatic depolymerization of the sacculi with mutanolysin and lysozyme.^38,57^ Fragments were analyzed by high-resolution quadrupole time-of-flight (Q-TOF) mass spectrometry, which revealed muropeptide fragments containing the click chemistry product (**Fig. 2d**). Despite the higher absolute fluorescence intensities resulting from treatment with elevated concentrations of **TetD** (**Fig. S2**), the apparent accumulation profiles for **Nmet0** remained nearly identical across the tested concentration range (**Fig. 2e**). This indicates that upon normalization, the accumulation profiles for peptides are broadly unaffected by the absolute amount of DBCO landmarks within the PG scaffold, consistent with our previous findings.^7^ Based on these results, we designated 25 μM of **TetD** as the labeling concentration for PG labeling in all subsequent assays. Importantly, no significant changes in cell morphology or cell envelope integrity were observed upon treatment with **TetD**, as evidenced by microscopy (**Fig. S3**) and ethidium bromide (EtBr) accumulation/efflux experiments (**Fig. S4**). EtBr is used as an indicator of mycomembrane integrity, which when disrupted, results in increased intracellular accumulation of EtBr, leading to elevated cellular fluorescence.^58–62^

Next, we tested the tolerance of the PAC-MAN assay toward a range of azide-tagged fluorophores. In addition to **Fl-az**, we evaluated 3-azido-7-hydroxy coumarin (**Co-az)**, azido-tetramethyl rhodamine (**TMR-az**), azido-sulfo Cy3 (**sCy3-az**), and azido-sulfo Cy5 (**sCy5-az**) with *Msm* cells (**Fig. 2f**). As expected, the resulting apparent accumulation curves overlapped across the different azide-tagged fluorophores used. This indicates that upon normalization, the accumulation profiles are largely unaffected by the choice of azide-tagged fluorophores used to detect the unoccupied DBCO landmarks within the PG scaffold. Considering that cellular fluorescence signals for cells treated with **Fl-az**, **sCy3-az**, or **sCy5-az,** all exhibited large signal-to-noise ratios (**Fig. S5**), we chose **Fl-az**, which is more widely available, for further experiments.

Similarly, we observed nearly identical normalized cellular fluorescence responses in the PAC-MAN assay when using various concentrations of **Fl-az** (**Fig. 2g**) or during varying incubation periods with **Fl-az** (**Fig. 2h**) in the chase step. However, longer incubation periods with the test molecule during the pulse step resulted in higher apparent accumulation (**Fig. 2i**), which is expected given the nature of detection upon periplasmic arrival, which is analogous to the mammalian CAPA.^56^ We note that the second baseline peptide (**Lin0**) underwent similar assay development steps as **Nmet0,** demonstrating comparable results (**Fig. S6**). Similar assay parameters and features for the PAC-MAN assay were also identified when optimized with azide-tagged small molecules.^7^ Together, these conditions established the workflow for analyzing test peptides in PAC-MAN (**Fig. 2j**), in which the concentration and incubation time of the test peptides were the most critical parameters.

### Effect of Backbone *N-*alkylation on Peptide Accumulation

Methylation of the amide backbone in peptides and peptidic natural products has been previously evaluated for its ability to modulate membrane permeability in mammalian systems.^31,63–66^ This structural edit can alter the hydrogen bonding networks between the peptides and surrounding water molecules, potentially reducing the overall hydrophilicity of the peptide and promoting its desolvation prior to passive diffusion across a hydrophobic environment (**Fig. S7**). To investigate this feature using PAC-MAN, we constructed libraries of *N*-methylated peptides derived from the unmodified peptide, **Nmet0**. In the first subseries, peptides **Nmet1** to **Nmet5** contain a varying total number of methylation marks within their backbone (**Fig. 3a**).

**Fig. 3.**
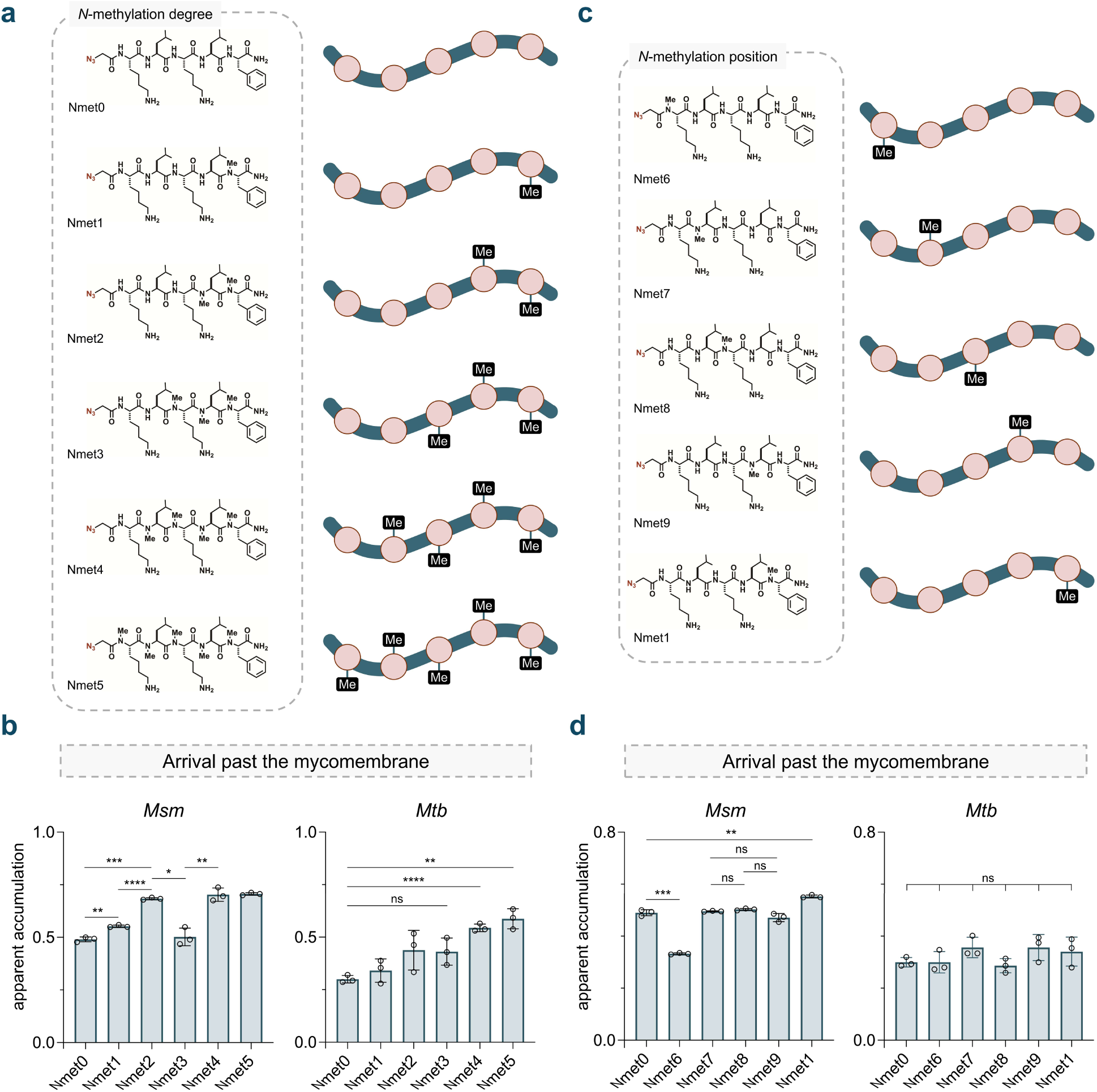
**(a)** Chemical structures of the *N*-methylation sub-library with varying degrees of backbone *N*-methylation (**Nmet0**-**5**). **(b)** Apparent accumulation of the *N*-methylation sub-library with varying degrees of backbone *N*-methylation across the mycomembrane in *Msm* and *Mtb* following 1 h of treatment with 50 µM compound. **(c)** Chemical structures of the *N*-methylation sub-library with varying positions of backbone *N*-methylation (**Nmet6-9** and **Nmet1**). **(d)** Apparent accumulation of the *N*-methylation sub-library with varying positions of backbone *N*-methylation across the mycomembrane in *Msm* and *Mtb* following 1 h of treatment with 50 µM compound. Data are represented as mean ± SD (n= 3). P-values were determined by a two-tailed t-test (ns = not significant, *p < 0.1, **p < 0.01, ***p < 0.001, ****p < 0.0001).

Our data revealed a significant increase in apparent accumulation with a single methylation added to the backbone of the *C*-terminal residue in **Nmet1**. This increase was even more pronounced with the addition of a second methylation for **Nmet2**. However, adding more than three methylation groups to the peptide did not further enhance its accumulation in *Msm*. Interestingly, the trimethylated molecule, **Nmet3**, deviated from this trend, exhibiting a reduction in apparent accumulation (**Fig. 3b**). Previous studies in mammalian systems found that a higher number of *N*-methyl groups may not necessarily correlate with improved permeability, potentially due to conformational preferences that are difficult to predict.^67,68^ In *Mtb*, no statistical differences were observed among the first three methylation marks, while an improvement was noted beyond three (**Fig. 3b**).

In the second subseries (**Nmet6** to **Nmet9**), a single methylation was systematically installed across the peptide backbone of **Nmet0** (**Fig. 3c**). Similar apparent accumulation was observed when methylation was applied to the central backbone amides (**Nmet7-9**); however, a slight decrease in accumulation was noted when the methylation mark was positioned at the *N-*terminus for **Nmet6** (**Fig. 3d**). Interestingly, no significant differences in accumulation were found within the mono-methylation series in *Mtb* (**Fig. 3d**). Extensive work by the Kessler and Lokey laboratories has highlighted the structural elements related to the degree and positional effects of backbone *N*-methylation on mammalian permeability in the context of oral bioavailability.^67–73^ These data suggest an overall trend that backbone *N*-methylation can modulate peptide accumulation across the mycomembrane, although the optimal degree and position may be species-specific.

To explore the contribution of the backbone amides in a different but biologically relevant context, we constructed a peptoid library that retained the same side chain sequence as the *N*-methylation library. Similar to *N*-methylated peptides, peptoids lack the hydrogen bond donor capacity of the backbone amide^74–79^, which could potentially enhance their permeability compared to their peptide counterparts.^74,80^ Like the *N*-methylation peptide library, peptoids **Nalk1** to **Nalk5** were assembled with an increasing number of *N*-substituted glycine units from the *N-* to the *C-*terminus (**Fig. S8a**) to systematically evaluate different degrees of backbone *N*-alkylation. Peptoids **Nalk5** to **Nalk9** were synthesized to vary the positions of a single *N*-substituted glycine unit across the peptidic backbone (**Fig. S8b**). In contrast to our results for backbone *N*-methylated peptides (**Nmet1**-**9**), we observed a much greater impact and variability in apparent accumulation with *N*-alkylated peptoids in *Msm*. The apparent accumulation of the peptoids significantly increased, reaching a maximum level with three *N*-substituted glycine units; however, an unexpected drop was noted for the fully substituted peptoid, **Nalk5**. No significant difference in apparent accumulation was observed for the *N*-terminal alkylated peptoid **Nalk6** compared to the base peptide **Nmet0**. Additionally, there was no preference for substitutions on the central residues for **Nalk7-9** (**Fig. S8c**). Overall, our results suggest that backbone alkylation in peptoids can lead to higher accumulation across the mycomembrane compared to their corresponding canonical versions.

Next, we aimed to test potential differences in SPAAC reactivity within our series of azide-tagged molecules as they engage with the installed DBCO landmarks. Previous work from our group investigated the reactivity of azide-tagged small molecules using a bead-based assay.^38^ Flow cytometry-compatible DBCO-coated beads, chemically modified from commercially available amino-functionalized beads, served as surrogates for mycobacterial cells without the critical mycomembrane barrier. Similar to our live cell PAC-MAN assay, beads were incubated with the same concentration of azide-tagged test molecules, followed by a chase step with an azide-tagged fluorophore. Background fluorescence levels from unmodified beads treated with **Fl-az** were normalized to 0%, while fluorescence levels from DBCO-modified beads treated with the vehicle were normalized to 100%. All molecules in both the *N*-methylation library and the peptoid library fully reacted with the DBCO landmarks on the beads (**Fig. S9, S10**). These results strongly support the notion that our apparent accumulation findings across the mycomembrane reflect the level of molecule arrival within the periplasmic space. We also investigated the integrity of the mycobacterial cell envelope after metabolic labeling by **TetD** and subsequent treatment with our library by using EtBr accumulation measurements. No differences were detected in EtBr accumulation upon treatment with different azide molecules (**Fig. S11, S12**). These findings suggest that the mycomembrane is not artificially disrupted during incubation with the azide peptides, supporting our assay as a reliable method for testing the accumulation of modified peptides past this critical barrier.

### Effect of Macrocyclization on Permeability

Cyclization has been extensively explored to improve the pharmacokinetic properties of large molecules such as peptides.^81,82^ Cyclic molecules present a more compact and favorable shape compared to their extended acyclic counterparts, which can facilitate their penetration through membrane barriers.^29,34,83^ The cyclic conformation may promote intramolecular hydrogen bonding and reduce the exposure of polar functional groups to the surrounding environment, thereby decreasing their solubility in water and promoting membrane integration (**Fig. 4a**). For the assembly of our macrocyclic peptide library, we chose a thioether-based cyclization strategy because of its metabolic stability and ease of formation under mild conditions.^84^ Additionally, thioether macrocyclic peptides are being extensively explored as potential drug scaffolds.^85–88^ We hypothesized that a combination of two modifications on peptides would enable in-solution cyclization: (1) an *N*-chloroacetyl group coupled onto the *N*-terminus of the peptide, and (2) a cysteine residue incorporated into the peptide sequence (**Fig. S13**). Importantly, we avoided placing cysteine residues at the *C*-terminal position due to concerns regarding potential epimerization^89^ and the formation of β-piperidinylalanine^90^ during peptide chain extension. For each macrocyclic peptide, we designed a linear counterpart with a largely preserved sequence, while replacing cysteine residues with serine and acetylating the *N*-terminus to closely mimic the composition of the cyclic peptide. We did not retain the cysteine residues in the linear peptides due to their susceptibility to rapid oxidation in the live cell assay, which could lead to various unique oxidized structures that are difficult to measure. Based on this macrocyclic library design, we synthesized three distinct sub-libraries combining standard solid-phase peptide synthesis and in-solution cyclization.

**Fig. 4.**
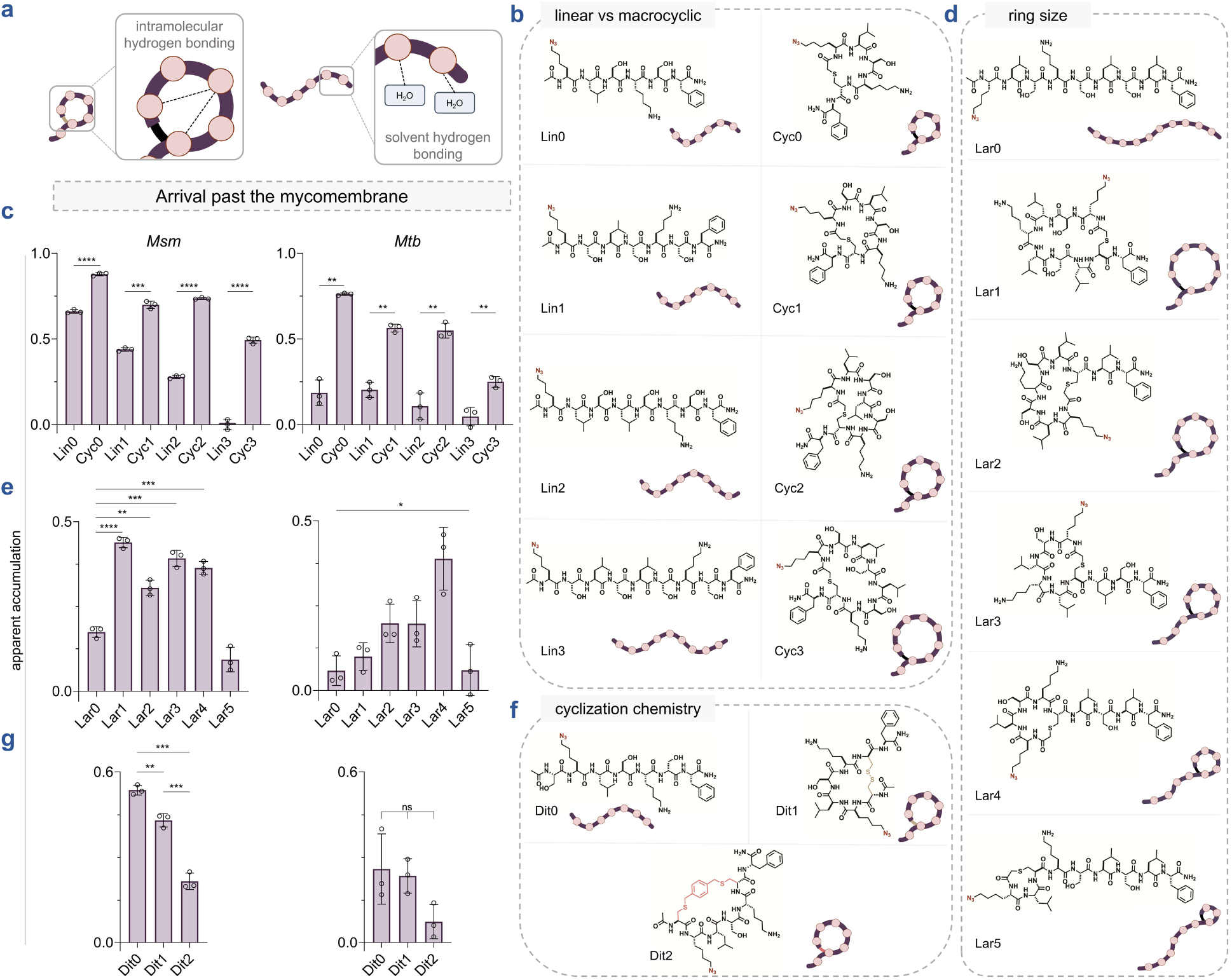
**(a)** Schematic illustration of a macrocyclic and a linear peptide interacting with the surrounding water molecules in an aqueous solution. Hydrogen bonds are indicated by dashed lines. Macrocyclization can promote intramolecular hydrogen bonding resulting in the reduction of hydrogen bonds between the backbone amides and the solvent, thereby potentially reducing membrane permeation associated desolvation penalty. **(b)** Chemical structures of the linear vs macrocyclic sub-library members containing macrocyclic peptides of increasing sizes (**Cyc0**-**3)** and their linear counterparts (**Lin0-3**). **(c)** Apparent accumulation of the linear vs macrocyclic sub-library across the mycomembrane in *Msm* and *Mtb* following 2 h of treatment with 50 µM compound. **(d)** Chemical structures of the ring size (lariat) sub-library members containing macrocyclic peptides of increasing ring sizes (**Lar1**-**5**) and their linear counterpart (**Lar0**). **(e)** Apparent accumulation of the ring size sub-library across the mycomembrane in *Msm* and *Mtb* following 2 h of treatment with 50 µM compound. **(f)** Chemical structures of the cyclization chemistry sub-library members containing a disulfide bonded macrocyclic peptide (**Dit1**), a bis-electrophilic linker based macrocyclic peptide (**Dit2**) and their linear counterpart (**Dit0**). **(g)** Apparent accumulation of the cyclization chemistry sub-library across the mycomembrane in *Msm* and *Mtb* following 2 h of treatment with 50 µM compound. Data are represented as mean ± SD (n= 3). P-values were determined by a two-tailed t-test (ns = not significant, *p < 0.1, **p < 0.01, ***p < 0.001, ****p < 0.0001). (*N*-*H*) with deuterium, with the exchange rate reflecting the solvent accessibility of the *N*-*H* bonds. The HDX process is tracked using ^1^H-NMR spectroscopy. **(b)** ^1^H-NMR spectra illustrating the time-dependent HDX of the backbone amide protons in **Lin0** and **Cyc0** over 1 h. **(c)** Fraction of K_az_1 *N-H* protons of **Lin0** and **Cyc0** exchanged to *N-D* over time during HDX. **(d)** Fraction of L2 *N-H* protons of **Lin0** and **Cyc0** exchanged to *N-D* over time during HDX. **(e)** 3-D representation and chemical structure of **Cyc0** showing hydrogen bonding observed through molecular dynamics simulations. **(f)** Comparison of serum stability of **Lin0** and **Cyc0** incubated in mouse serum at 37 °C over 1 h. For serum stability analyses, one-phase decay curves were fitted to the data using GraphPad Prism.

Recognizing that the molecular size of macrocyclic peptides can influence membrane permeability,^91,92^ we designed a set of macrocyclic peptides with varying numbers of leucine and serine residues to systematically increase the cycle size (**Cyc0**-**3**) (**Fig. 4b**). This sub-library was then tested using our PAC-MAN assay in live mycobacteria. Generally, across all sizes tested, the macrocyclic peptides demonstrated better apparent accumulation across the mycomembrane compared to their linear counterparts, as observed in both *Msm* and *Mtb* (**Fig. 4c**). Additionally, in both organisms, the apparent accumulation of both linear and macrocyclic peptides diminished with increasing molecular size (**Fig. 4c**). **Cyc0**, the smallest macrocyclic peptide in our panel, exhibited better accumulation than any other member of this macrocyclic sub-library, which was comparable to the most permeable molecule identified in our previous work.^38^

Next, inspired by natural products, we focused our attention on lariat-shaped molecules. These molecules are characterized by a looped or “lasso” structure and have recently gained recognition for their utility as drug scaffolds^93^ and membrane permeators.^68^ A subset of these molecules has shown promising antimycobacterial activity, such as lassomycin.^94^ Their flexible linear tails provide additional functional versatility that can be modulated to specifically target proteins of interest. We prepared a series of lariat peptides by cyclizing between an *N*-terminal chloroacetyl group, and a cysteine side chain positioned at various distances from the *N*-terminus, thereby decreasing the size of their cyclic portion (**Lar1**-**5**) (**Fig. 4d**). The linear control, **Lar0**, was designed to be *N*-acetylated without any cysteine residues. The macrocyclic lariat-shaped peptides **Lar1**-**4** demonstrated better accumulation profiles than the linear **Lar0** in *Msm* (**Fig. 4e**). In contrast, **Lar5** exhibited poor accumulation in *Msm*, which we attributed to the small size of its ring, leaving a substantial portion of the peptide flexible and exposed to the solvent (**Fig. 4e**). Within the lariat-shaped macrocyclic series (**Lar1**-**5**), no discernible trend was observed in either *Mtb* or *Msm*. However, **Lar4** appeared to have a privileged scaffold over the other lariats and the linear control **Lar0** in terms of accumulation into *Mtb* (**Fig. 4e**).

We recognized that the method of cyclization could influence the conformation and structural rigidity of peptides, which might affect their interaction with the mycomembrane. It has been shown in mammalian systems that the structural features of the linker can alter peptide accumulation.^95^ In mycobacteria, we expanded our cyclic peptide library to include peptides cyclized using two cysteine side chains. First, we synthesized an *N*-acetylated dithiol peptide scaffold that possesses the same sequence as **Cyc0**, but with an additional cysteine residue at the *N*-terminus. This dithiol peptide was then directly oxidized to generate a disulfide bond between the two cysteine side chains, resulting in **Dit1** (**Fig. 4f**). This analysis is significant due to the growing interest in disulfide-cyclized peptides as drug candidates.^96–100^ Additionally, we cyclized the linear dithiol peptide scaffold with a bis-electrophilic linker, yielding **Dit2**, which contains two thioether bonds (**Fig. 4f**). These types of linkers are frequently used in constructing macrocyclic peptide libraries due to their broad sequence compatibility and suitability for late-stage conformational diversification,^101–104^ prompting us to investigate a test case. Lastly, we synthesized a linear control, **Dit0**, by replacing the cysteine residues with serine (**Fig. 4f**).

Among this sub-library, we observed that the linear control, **Dit0**, exhibited the highest level of accumulation in *Msm* compared to its macrocyclic counterparts **Dit1** and **Dit2** (**Fig. 4g**). For **Dit1**, this may be partly due to interception by thiol/disulfide-displaying proteins in the cell envelope before it reaches the PG.^105^ Thiol-disulfide exchange reactions between disulfide-containing molecules and exofacial protein thiols/disulfides have previously been shown to modulate permeability.^106,107^ In the case of **Dit2**, we pose that the hydrophobicity of the linker itself may significantly contribute to peptide accumulation, similar to observations made in mammalian systems.^95^ In *Mtb*, we observed no statistical differences in apparent accumulation within this series (**Fig. 4g**). The difference in the apparent accumulation profiles of **Cyc0** and **Dit1** can be attributed to the difference of just one amino acid residue and the mode of cyclization. As previously established, we confirmed that the macrocyclic peptide library readily reacted with our DBCO-modified beads (**Fig. S14**) and that cell integrity remained unaffected upon treatment with the peptides, as indicated by EtBr analysis (**Fig. S15**). Taken together, our data suggest a general trend that macrocyclization can improve peptide accumulation across the mycomembrane, with additional considerations regarding ring size and the mode of cyclization.

We then sought to decipher the mechanistic basis for the enhanced accumulation of macrocyclic peptides in comparison to their linear counterparts. By evaluating the physicochemical properties of our libraries, we found that most library members, including the peptide with the highest accumulation levels, significantly exceeded the rule-of-five (Ro5) parameters established by Lipinski^11^ and Veber^102^ (**Fig. S16**). This finding highlights a gap in our understanding of the molecular characteristics that enable traversal of the mycomembrane. It is well established that intramolecular hydrogen bonding and the solvent-accessible surface area of a molecule are critical factors influencing permeability across membrane bilayers.^109–112^ This feature often informs the rationale for both the macrocyclization^82,113^ and *N*-methylation^114^ of peptides to improve membrane permeability by reducing solvent accessibility. Therefore, we aimed to determine whether variable solvent accessibility of the amide *N−H* between the linear and macrocyclic peptides is responsible for the differences observed in their accumulation across the mycomembrane.

To investigate this, we conducted hydrogen/deuterium exchange (HDX) experiments with the molecules **Cyc0** and **Lin0** using ^1^H-NMR (**Fig. 5a**). We characterized these molecules using 1D and 2D-NMR, assigning all *N-H* resonances. Generally, the backbone amide hydrogens of **Lin0** exchanged more rapidly with the deuterated solvent molecules than those of **Cyc0**, showing complete exchange within an hour (**Fig. 5b**). As expected, not all amide hydrogens exchanged equally; for example, the amide hydrogen on the azide-lysine (K_az_) residue of **Lin0** exhibited complete exchange within the dead time of the assay (∼5 mins) (**Fig. 5c**). In contrast, the exchange of the amide hydrogen on K_az_ of **Cyc0** was not complete even at the 25-minute mark (**Fig. 5c**).

**Fig. 5.**
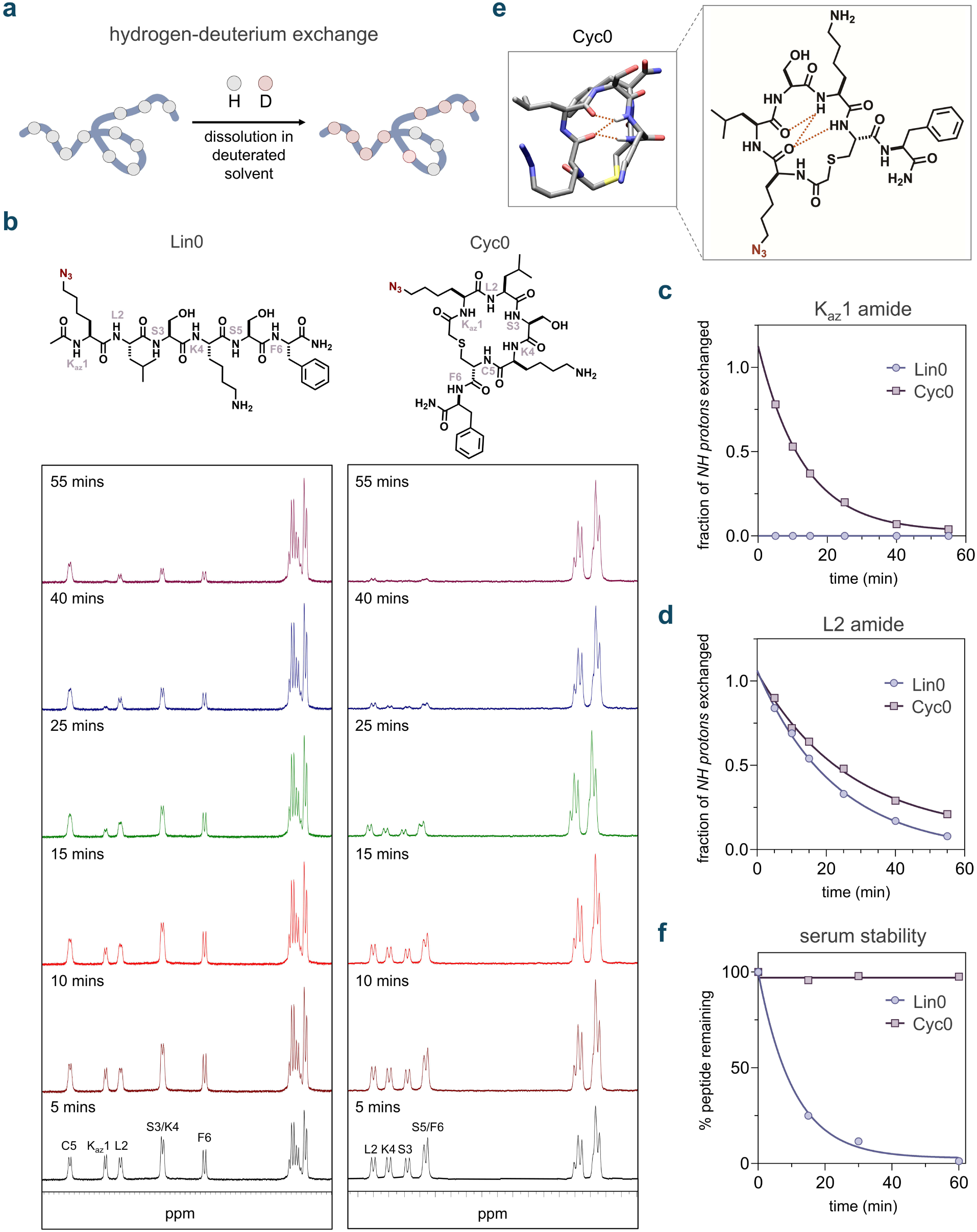
**(a)** Schematic illustration of the hydrogen-deuterium exchange (HDX) analysis of peptides. Peptides are incubated in a D2O buffer, allowing the exchange of amide protons (N-H) with deuterium, with the exchange rate reflecting the solvent accessibility of the NH bonds. The HDX process is tracked using 1H-NMR spectroscopy. (**b**) 1H-NMR spectra illustrating the time-dependent HDX of the backbone amide protons in **Lin0** and **Cyc0** over 1 h. (**c**) Fraction of Kaz1 N-H protons of **Lin0** and **Cyc0** exchanged to N-D over time during HDX. (**d**) Fraction of L2 N-H protons of **Lin0** and **Cyc0** exchanged to N-D over time during HDX. (**e**) 3-D representation and chemical structure of **Cyc0** showing hydrogen bonding observed through molecular dynamics simulations. (**f**) Comparison of serum stability of **Lin0** and **Cyc0** incubated in mouse serum at 37 °C over 1 h. For serum stability analyses, one-phase decay curves were fitted to the data using GraphPad Prism.

To further compare these two peptides, we focused on the backbone amide L2, which was present in both molecules and was well-resolved in the NMR spectra. The L2 amide on **Cyc1** exhibited a half-life of 18 minutes, compared to 16 minutes for **Lin0** (**Fig. 5d**). This observation highlights how macrocyclization can potentially shield the L2 backbone amide from solvent hydrogen bonding. The L2 backbone amide of **Cyc0** may also participate in intramolecular hydrogen bonding, which could hinder exchange with solvent molecules. To support this analysis, molecular dynamics (MD) simulations were performed with **Cyc0** and **Lin0**. Our results indicated that the **Cyc0** backbone amides extensively participate in intramolecular hydrogen bonding (**Fig. 5e**), while no observable intramolecular hydrogen bonding was detected for the linear counterpart, **Lin0**.

Another well-recognized biochemical consequence of peptide cyclization is greater resistance to proteases.^111,115–117^ This protection may be partly attributed to the inability of the macrocyclic peptide to be recognized by the active site of serine proteases in the ideal conformation, given its limited flexibility.^116,118^ Upon testing the metabolic stability of both **Cyc0** and **Lin0**, we observed that **Lin0** fully degraded within 1 hour of serum incubation, while **Cyc0** remained stable with no significant degradation (**Fig. 5f**). A similar observation was made regarding backbone *N*-methylation, where peptide **Nmet5** exhibited considerably greater serum stability than peptide **Nmet0,** likely due to the steric hindrance introduced by *N*-methylation toward serine proteases (**Fig. S17**).^119,120^

### Effect of Structural Edits on Peptide Antibiotics

Building on our findings, we set out to modulate the activity and accumulation profile of a peptidic antibiotic with potential activity against mycobacteria by applying these structural edits. Tridecaptin A1 is a non-ribosomal lipopeptide known for its antimicrobial activity against Gram-negative bacteria. Its mode of action is related to its binding to the bacterial cell wall precursor lipid II and subsequently disrupting the proton motive force.^121,122^ Binding to lipid II is a common mode of action for many potent antibiotics, including the glycopeptide vancomycin.^123^ To the best of our knowledge, tridecaptin A1 has not been shown to be active against mycobacteria. Given the theoretical necessity of crossing the mycomembrane to bind lipid II, the inherent lack of appreciable antimycobacterial activity may be attributed to insufficient accumulation of the peptide in the periplasmic space. Therefore, we propose that structural edits aimed at enhancing accumulation while maintaining target lipid II engagement could improve its antibacterial properties.

Considering the structural similarities between mycobacterial lipid II and Gram-negative lipid II, we hypothesized that backbone *N*-methylation or macrocyclization could potentially modulate the activity of tridecaptin A1 against mycobacteria by enhancing its accumulation past the mycomembrane. In redesigning tridecaptin, we started with the octanamide analog, Octyl-tridecaptin, as it retained biological activities and is more synthetically tractable.^124^ Based on previous studies, an alanine scan found that amino acid residues in positions 1, 4, 6, 10, and 11 can be substituted with alanine without significantly compromising their biological activity against Gram-negative pathogens.^125^ Furthermore, it was reported that the side chain carboxylate of Glu10 can be conjugated with antibiotic warheads to improve its activity *in vitro.*^126^ Relying on these prior efforts, we designed a series of analogs from the parental peptide, substituting the Glu10 residue with an azide-bearing lysine analog (**tri**) for compatibility with our assay (**Fig. 6a**).

**Fig. 6.**
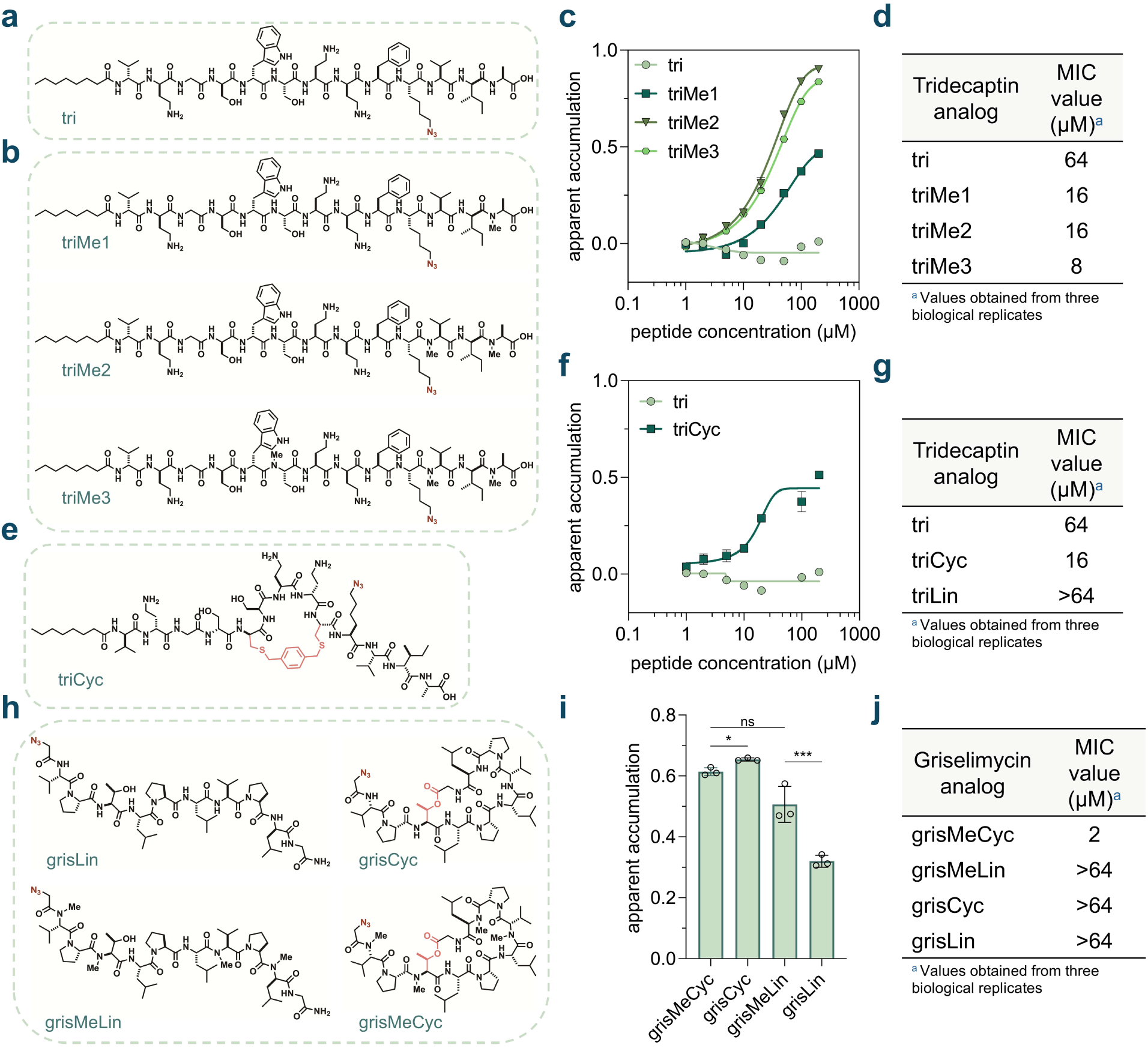
**(a)** Chemical structure of the peptide **tri,** the tridecaptin parent analog in our series. **(b)** Chemical structures of the peptides **triMe1**, **triMe2**, **triMe3**, the *N*-methylated tridecaptin analogs in our series. **(c)** Dose-response analysis showing the apparent accumulation of the *N*-methylated tridecaptin analogs across the mycomembrane in *Msm*. **(d)** MIC values of the *N*-methylated tridecaptin analogs against *Msm*. **(e)** Chemical structure of **triCyc**, the macrocyclic tridecaptin analog in our series. **(f)** Dose-response analysis showing the apparent accumulation of the macrocyclic tridecaptin analog **triCyc** in comparison to the parent tridecaptin analog **tri**, across the mycomembrane in *Msm.* **(g)** MIC values of the macrocyclic and linear tridecaptin analogs **triCyc** and **triLin**, in comparison to the parent tridecaptin analog **tri**, against *Msm*. **(h)** Chemical structures of the peptides **grisLin**, **grisCyc**, **grisMeLin**, and **grisMeCyc**, the griselimycin analogs in our study. **(i)** Apparent accumulation of the griselimycin analogs across the mycomembrane in *Msm* following 6 h of treatment with 50 µM compound. **(j)** MIC values of the of the griselimycin analogs against *Msm.* All MIC values were obtained from three biological replicates which showed identical results. Other data are represented as mean ± SD (n= 3). For dose-response curves, Boltzmann sigmoidal curves were fitted to the data using GraphPad Prism. All dose-response analyses involving the tridecaptin series were performed with a 1 h incubation period with the peptides. P-values were determined by a two-tailed t-test (ns = not significant, * p < 0.1, ** p < 0.01, *** p < 0.001, **** p < 0.0001).

*N*-methylation of the parent peptide should ideally not disrupt target binding. *N*-methylation sites were selected by examining the reported solution NMR structure of the tridecaptin A1-lipid II complex.^121^ The amide backbones of residues Ser6, Val11, or Ala13 were selected for methylation. Specifically, we generated monomethylated (**triMe1** with meAla13), dimethylated (**triMe2** with meVal11 and meAla13), and trimethylated (**triMe3** with meSer6, meVal11, and meAla13) analogs of **tri** (**Fig. 6b**). The dose-response accumulation profiles of the *N*-methylated peptides were then obtained *via* live cell PAC-MAN in *Msm* (**Fig. 6c, S18**). **triMe1** and **triMe2** exhibited better accumulation compared to the unmethylated **tri.** However, **triMe3** showed slightly worse accumulation than **triMe2**, suggesting that additional methylation on Ser6 negatively affects its apparent accumulation in mycobacteria. We noted that the unmethylated parent peptide **tri** demonstrated fluorescence intensities higher than those of the vehicle control at lower concentrations, resulting in negative apparent accumulation values (**Fig. S18**). We attributed this observation to potential cell envelope disruption by **tri**, which could theoretically modulate the accumulation level of the fluorophore and lead to higher fluorescence intensities. Indeed, incubation with **tri** significantly affected cell envelope integrity, allowing for higher EtBr accumulation (**Fig. S19**). No significant effects were observed in any of the *N*-methylated analogs. Importantly, we also ensured that the peptides readily reacted with our DBCO-modified beads (**Fig. S20**). Next, we studied the effect of *N*-methylation on the antimicrobial activity of **tri** against *Msm via* a Minimal Inhibitory Concentration (MIC) assay. A marked decrease in the MIC value of the peptide antibiotic was observed with an increasing degree of methylation, with trimethylated **triMe3** exhibiting an 8-fold reduction in MIC value compared to unmethylated **tri** (**Fig. 6d**). Interestingly, the unmethylated analog **tri** exhibited fast amyloid-like aggregation compared to the trimethylated analog **triMe3**, indicated by a thioflavin T (ThT) fluorescence assay (**Fig. S21**). The ThT fluorescence assay is a semi-quantitative method used for the analysis of amyloid-like aggregates in the system.^127,128^ This feature has been reported as an critical mechanism of action for the antimicrobial activity of teixobactin.^127–129^ On the contrary, in our hands, diminished aggregation in the case of the trimethylated analog **triMe3** was correlated with better antimycobacterial activity in comparison to **tri**. Overall, these data indicate that backbone *N*-methylation of **tri** enhances its activity against *Msm* by improving its apparent accumulation across the mycomembrane.

We then sought to analyze the effect of macrocyclization on the activity of **tri** against *Msm.* Octyl-tridecaptin A1 had previously been cyclized by replacing D-Trp5 and L-Phe9 with D- and L-cysteine residues, respectively, and the peptides were subsequently cyclized using bis-electrophilic linkers.^130^ Adopting a similar approach, we synthesized **triCyc**, wherein the peptide was cyclized with a bis-bromomethyl-benzene linker between the cysteine residues, and Glu10 was substituted with an azide-bearing lysine analog (**Fig. 6e**). The dose-response accumulation profile of **triCyc** was then analyzed *via* live cell PAC-MAN in *Msm* (**Fig. 6f, S22**). The cyclic form (**triCyc**) exhibited better accumulation across the mycomembrane compared to the parent peptide **tri.** To further elucidate the impact of macrocyclization, we also synthesized **triLin**, in which the D-Trp5 and L-Phe9 residues were replaced with D- and L-serine residues, respectively, to serve as a linear control (**Fig. S23a**). Interestingly, we observed no significant difference in accumulation between **triCyc** and **triLin** (**Fig. S23b, c**). Furthermore, EtBr accumulation experiments revealed that both **triCyc** and **triLin** had no significant effect on cell envelope integrity (**Fig. S24**). The peptides also readily reacted with our DBCO-labeled beads (**Fig. S25**). Next, we sought to examine the antimicrobial activity of **triCyc** and **triLin** against *Msm*. For **triCyc**, we also observed a 4-fold reduction in the MIC value when compared to the parent peptide **tri** (**Fig. 6g**). **triCyc** also exhibited over a 4-fold improvement in antimicrobial activity compared to **triLin**. Taken together, these data suggest that macrocyclization enhances the antimicrobial activity of **tri** against *Msm*.

We then aimed to investigate the influence of *N*-alkylation and macrocyclization on the antimicrobial activity and accumulation of a peptidic antibiotic that inherently possesses these structural features. Griselimycin, a peptide originally derived from Streptomyces,^131^ has been reported to exhibit antimicrobial activity against both *Msm* and *Mtb*.^132–134^ The antimicrobial activity of this peptide is mediated through its binding to the mycobacterial DNA polymerase III sliding clamp (DnaN), resulting in the disruption of DNA replication.^132^ Notably, the sliding clamps of other bacterial species have also been investigated as potential targets for griselimycin^135,136^ and other antimicrobial agents^137–142^. Structurally, griselimycin is a lariat-shaped peptide, cyclized through an ester bond between the *C*-terminus and the side chain hydroxyl group of its internal Thr3 residue. The peptide also contains four backbone *N*-methylation marks at residues - Val1, Thr3, Val7, and D-Leu9. To facilitate compatibility with the PAC-MAN assay, we designed analogs in which the *N*-terminal acetyl group present in the parental peptide was replaced with a 2-azido-acetyl group. Additionally, the 4-methyl-proline residues in the parental peptide were substituted with proline residues due to greater commercial availability, while acknowledging that this substitution may affect target engagement. The analogs were synthesized in both linear and cyclic forms, with or without backbone *N*-methylation, resulting in four distinct variants: **grisMeCyc**, **grisCyc**, **grisMeLin**, and **grisLin** (**Fig. 6h**).

The accumulation profiles of the griselimycin series were then obtained *via* live cell PAC-MAN in *Msm.* We observed that macrocyclization was important for the accumulation of **grisLin,** which had considerably lower accumulation than **grisCyc**. This effect was however absent in the case of **grisMeCyc** which did not significantly differ in its accumulation with respect to **grisMeLin** (**Fig. 6i**). The *N-*methylation data were interesting in that **grisCyc** in fact exhibited slightly improved accumulation over **grisMeCyc,** whereas **grisLin** accumulated to a much lower level than **grisMeLin** (**Fig. 6i**). As before, we ensure that the peptides readily reacted with our DBCO-labeled beads (**Fig. S26**)

Upon evaluating the antimicrobial activity of the peptides, the loss of macrocyclization in **grisMeCyc** resulted in over a 32-fold increase in the MIC value of **grisMeLin** (**Fig. 6j**). Similarly, the removal of methylation also led to a reduction in activity, with **grisCyc** performing weaker than **grisMeCyc** (**Fig. 6j**). The simultaneous loss of both macrocyclization and methylation was also detrimental to the peptide’s activity. Collectively, these results highlight the critical role of macrocyclization and *N*-methylation in maintaining the antimicrobial activity of griselimycin against *Msm*.

## DISCUSSION

Nature possesses a variety of biosynthetic machineries capable of constructing both small and large molecules. Over time, the evolution of small molecules with high molecular complexity and membrane permeability might lead us to anticipate that nature favors antibiotics with low molecular weights. After all, since most antibiotics target intracellular sites, they must overcome one or more membrane bilayers to reach these targets. However, this is not exclusively the case in nature. A substantial number of biologically active natural products, including many antibiotics, are large, polar, and often exceed the Rule of Five (Ro5) criteria. These molecules typically have a peptidic nature, a trait that can result from biosynthetic pathways, such as in ribosomally synthesized and post-translationally modified peptides (RiPPs). Larger peptide-like molecules offer significant advantages, primarily by interacting with their molecular targets more extensively, resulting in higher affinity and specificity. This size trade-off must be balanced with considerations of accumulation inside target cells. Consequently, it is commonly observed that peptidic antibiotics are extensively modified to maintain sufficient membrane permeability.

The assembly of most antibiotic peptides offers an avenue for unusual modifications, including the incorporation of structural elements that enhance permeability. While the range of structural alterations is vast, the alkylation of the amide backbone and macrocyclization are particularly notable. *N*-alkylation reduces the number of hydrogen bonds formed with the solvent, thereby decreasing the desolvation penalty associated with passive membrane diffusion. Some peptides, or molecules in general, can also exhibit chameleonic behavior during passive membrane diffusion, a trait enhanced by backbone alkylation.^30,31,143^ Indeed, *N*-methylation is used as an effective approach to improve peptide permeability through mammalian cell membranes.^31,64–66,144^ A similar approach to reducing the number of backbone hydrogen bond donors is to shift peptide side chains from the α-carbon to the backbone nitrogen atom to create peptidomimetics, i.e. peptoids. Peptoids are easy to synthesize from primary amines and allow the possibility of versatile side chain libraries – albeit at the cost of chiral centers on the backbone.^145^ Various peptoids have demonstrated effectiveness against drug-resistant bacterial pathogens.^75–79^ Proline residues, like peptoids, naturally lack backbone hydrogen bond donors due to alkylation. Notably, some peptide antibiotics against mycobacteria, such as griselimycin^132^ and callyaerin^146^ have three or more prolines in their structure. Similarly, macrocyclization enhances the suitability of peptides as drug candidates by improving passive permeability through intramolecular hydrogen bonding,^35^ thereby reducing the desolvation penalty associated with membrane permeation.^109–111^ Furthermore, the structural preorganization provided by macrocyclization mitigates entropic loss during target binding.^147^ Additionally, it can enhance other pharmacological properties, such as metabolic stability,^17,148^ which may be especially relevant in therapeutic contexts.

The field of *de novo* and natural product-based peptide antibiotics continues to expand, making the prospect of orally available peptide drugs increasingly realistic. However, it remains largely undefined how *N*-alkylation and macrocyclization affect accumulation in mycobacteria. Given the unique composition of the mycomembrane and its central role in determining which molecules can effectively impact mycobacterial cells, understanding the impact of these peptidic alterations is crucial for the design and redesign of natural product drug leads. In turn, a comprehensive understanding of how structural features drive accumulation will facilitate the identification of high-potential drug candidates and enable the strategic redesign of existing chemical scaffolds to enhance their accumulation within mycobacterial cells.

By assessing the accumulation of *N*-alkyl and peptoid-likes compounds in mycobacteria, our work demonstrated a general trend: higher degrees of *N*-alkylation usually led to higher accumulation levels past the mycomembrane.^149^ Moreover, in our panels of macrocyclic peptides, we observed that generally cyclic peptides had higher apparent accumulation across the mycomembrane than their linear counterparts, underscoring the significant impact of ring size and cyclization chemistry in some cases. It is important to note that the field differs on whether cyclization helps improve accumulation. A report from the Kodadek laboratory challenges the notion that macrocyclization necessarily enhances the cell permeability of linear peptides in every case,^150^ presenting findings that contradict other studies supporting such an improvement.^35,151^ Our findings reveal that the improvement in accumulation past the mycomembrane may not be absolute, but rather context-dependent. We propose that critical factors such as lipophilicity, ring size, and cyclization chemistry should be carefully considered when attempting to enhance permeability through macrocyclization. Such empirical evaluations are enabled by methodologies like our PAC-MAN assay.

Based on our findings regarding the impact of *N*-alkylation and macrocyclization, we applied these principles to a known antibiotic. Unexpectedly, we observed that the parent molecule, **tri**, inherently disrupted the mycobacterial cell envelop, whereas the *N*-methylation of the tridecaptin peptide mitigated the membrane disruption effect and improved its antimycobacterial activity. In contrast, previous studies reported that backbone *N*-methylation of teixobactin^152^ and other reported antimicrobial peptides^153^ negatively affected their antimicrobial activity. These results emphasize the value of empirical analyses of accumulation, separate from MIC assessments, in understanding how structure influences accumulation. Overall, both *N*-alkylation and macrocyclization enhanced the efficacy of tridecaptin A1 against mycobacteria, underscoring the importance of our new insights into the role of structural factors in improving accumulation beyond the mycomembrane. This case illustrates how redesigning a known antimicrobial molecule, previously not shown to be active against mycobacteria, can enhance its accumulation and activity. Notably, since nature frequently combines these structural features in molecular design, further analysis of tridecaptin A1 peptides and other molecules modified with both *N*-alkylation and macrocyclization presents an intriguing research opportunity. We are currently exploring additional modifications to tridecaptin to optimize its activity and potentially guide an analog toward clinical evaluation.

To investigate the impact of systematically removing these inherent structural features, we examined griselimycin, a macrocyclic, *N*-methylated peptide that is active against *Msm*. Our results revealed a significant loss of activity upon the deletion of one or both of these structural features, highlighting the critical role they play in preserving efficacy against *Msm*. Notably, the indispensability of these features in this case underscores how even subtle structural modifications can profoundly impact antimicrobial activity, highlighting the importance of optimizing these key elements in the design of effective antibiotics.

PAC-MAN offers several key advantages over other technologies, including ease of deployment, which makes it potentially adoptable by most laboratories; suitability for high-throughput analysis; and the capability to achieve subcellular resolution, as opposed to whole-cell analysis. However, a limitation of PAC-MAN is its inability to provide information regarding the arrival of the test molecule in the cytosol. Further work is needed to develop methodologies that identify structural modifications capable of enhancing accumulation into the cytoplasm. Since the mycomembrane is often regarded as the primary barrier to accumulation, we propose that establishing guidelines to facilitate passage through this membrane represents a critical first step in creating an anti-mycobacterial pipeline to combat deadly diseases, such as TB.

In this work, we present the first systematic governing rules for the rational redesign of potential antimycobacterial agents. Our comprehensive exploration of structural modifications – specifically *N*-alkylation and macrocyclization – provides key insights into designing more effective antimycobacterial agents. Our study elucidates how these modifications enhance peptide accumulation across the intricate mycomembrane, challenging the conventional belief that large hydrophilic molecules cannot efficiently penetrate this barrier. The novel application of these strategies to the peptide antibiotic tridecaptin A1 highlights their potential to overcome intrinsic resistance mechanisms and optimize antibacterial efficacy. Similarly, the marked loss of activity upon deletion of these structural features in griselimycin underscores their fundamental role in preserving its antimicrobial activity and thus the importance of structural optimization in antibiotic design. Moreover, our PAC-MAN assay not only serves as a high-throughput platform for evaluating accumulation dynamics but also lays the groundwork for future empirical investigations into improving cytoplasmic delivery. These findings significantly contribute to the growing arsenal against TB by guiding the rational design of next-generation therapeutics capable of breaching the formidable defenses of mycobacteria. Further research integrating these structural features with conventional drug frameworks may yield potent treatments tailored to combat the escalating threat of drug-resistant mycobacterial strains.

## ACKNOWLEDGEMENT

This study was supported by the NIH grants GM124893-01 (M.M.P.), R01AI178975-01 (M.M.P. and W.I) and R01AI179080-01 (M.M.P., W.I., and M.S.S.). We thank Jeff Ellena, Department of Chemistry, University of Virginia, for NMR spectroscopy assistance.

## SUPPORTING INFORMATION

Additional figures, tables, and materials/methods are included in the supporting information file.

## Notes

### Competing Interest Statement

The authors have declared no competing interest.

### Summary of Updates

There were additional experiments that are now added to Figure 6.

