## Supplementary Information for "Systematic Determination of the Impact of Structural Edits on Peptide Accumulation into Mycobacteria"

[5]Department of Chemistry
[6]Departments of Biological Sciences and Bioengineering
Lehigh University
Bethlehem, PA, USA

### TABLE OF CONTENTS


| SUPPLEMENTARY FIGURES | S5-S30 |
|---|---|
| **Figure S1.** Chemical structure of peptide **Lin0**. | S5 |
| **Figure S2.** Concentration dependent labeling of *Msm* cells with **TetD**. | S6 |
| **Figure S3.** Phase-contrast microscopy of *Mtb* cells labeled with **TetD**. | S7 |
| **Figure S4.** EtBr based mycomembrane integrity analysis of *Msm* cells labeled with **TetD**. | S8 |
| **Figure S5.** Signal-to-noise ratios observed upon treatment of **TetD** labeled *Msm* cells with various azide-tagged fluorophores. | S9 |
| **Figure S6.** PAC-MAN assay optimization with **Lin0** in *Msm.* | S10 |
| **Figure S7.** Schematic illustration of an *N*-methylated peptide interacting with the surrounding environment. | S11 |
| **Figure S8.** Structures and apparent accumulation of the *N*-alkylation (peptoid) sub-library across the mycomembrane in *Msm*. | S12 |
| **Figure S9.** DBCO-modified polystyrene bead reactivity assay with the *N*-methylation sub-library. | S13 |
| **Figure S10.** DBCO-modified polystyrene bead reactivity assay with the *N*-alkylation sub-library. | S14 |
| **Figure S11.** EtBr based mycomembrane integrity analysis of *Msm* cells treated with the *N*-methylation sub-library. | S15 |
| **Figure S12.** EtBr based mycomembrane integrity analysis of *Msm* cells treated with the *N*-alkylation sub-library. | S16 |
| **Figure S13.** Thioether-based cyclization of macrocyclic peptides. | S17 |
| **Figure S14.** DBCO-modified polystyrene bead reactivity assay with the macrocyclization library. | S18 |
| **Figure S15.** EtBr based mycomembrane integrity analysis of *Msm* cells treated with the macrocyclization library. | S19 |
| **Figure S16.** Physicochemical properties of the *N*-methylation and macrocyclization peptide libraries. | S20 |
| **Figure S17.** Serum stability of **Nmet0** and **Nmet5**. | S21 |
| **Figure S18.** Apparent accumulation of the *N*-methylated tridecaptin series across the mycomembrane in *Msm*. | S22 |
| **Figure S19.** EtBr based mycomembrane integrity analysis of *Msm* cells treated with the *N*-methylated tridecaptin series. | S23 |
| **Figure S20.** DBCO-modified polystyrene bead reactivity assay with the *N*-methylated tridecaptin series. | S24 |
| **Figure S21.** ThT fluorescence assay comparing tridecaptin analogs **triMe3** and **tri**. | S25 |

| | |
|---|---|
| **Figure S22.** Apparent accumulation of the macrocyclic and parent tridecaptin analogs **triCyc** and **tri** across the mycomembrane in *Msm*. | S26 |
| **Figure S23.** Structure of the linear tridecaptin analog **triLin** and its apparent accumulation in comparison to the macrocyclic tridecaptin analog **triCyc**, across the mycomembrane in *Msm.* | S27 |
| **Figure S24.** EtBr based mycomembrane integrity analysis of *Msm* cells treated with the linear and macrocyclic tridecaptin analogs **triLin** and **triCyc**. | S28 |
| **Figure S25.** DBCO-modified polystyrene bead reactivity assay with the linear and cyclic tridecaptin analogs **triLin** and **triCyc**. | S29 |
| **Figure S26.** DBCO-modified polystyrene bead reactivity assay with the griselimycin series. | S30 |
| **EXPERIMENTAL PROCEDURES** | S31-S42 |
| **Materials** | S31 |
| **Biological Methods** | S33 |
| Mycobacteria cell culture | S33 |
| DBCO-tetrapeptide (**TetD**) based labeling of *Msm* | S33 |
| DBCO-tetrapeptide (**TetD**) based labeling of *Mtb* | S33 |
| Accumulation assay (PAC-MAN) across the *Msm* mycomembrane | S33 |
| Accumulation assay (PAC-MAN) across the *Mtb* mycomembrane | S34 |
| Isolation of peptidoglycan and muropeptides | S34 |
| Ethidium bromide (EtBr) whole-cell accumulation assay | S35 |
| DBCO-modified polystyrene bead reactivity assay | S35 |
| Calculation of physiochemical properties of the test molecules | S36 |
| Hydrogen-deuterium exchange (HDX) studies | S36 |
| Molecular dynamics (MD) simulation studies | S36 |
| Serum stability assay | S37 |
| Minimum inhibitory concentration (MIC) assay | S37 |
| Thioflavin T (ThT) fluorescence assay | S37 |
| **Synthesis and Characterization of Test Molecules** | S39 |
| General procedure for the solid-phase synthesis of peptides | S39 |
| Synthesis of **TetD** | S39 |

| | |
|---|---|
| Synthesis of the macrocyclization peptide library | S39 |
| Synthesis of the *N*-methylation and *N*-alkylation peptide libraries | S40 |
| Synthesis of the tridecaptin series | S41 |
| Synthesis of the griselimycin series | S41 |
| Characterization of test molecules | S42 |
| **REFERENCES** | S89-S90 |

#### SUPPLEMENTARY FIGURES

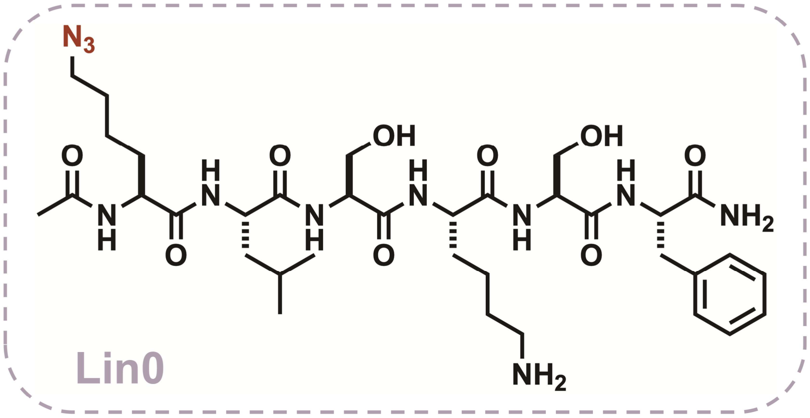


**Figure S1.** Chemical structure of the peptide **Lin0**.

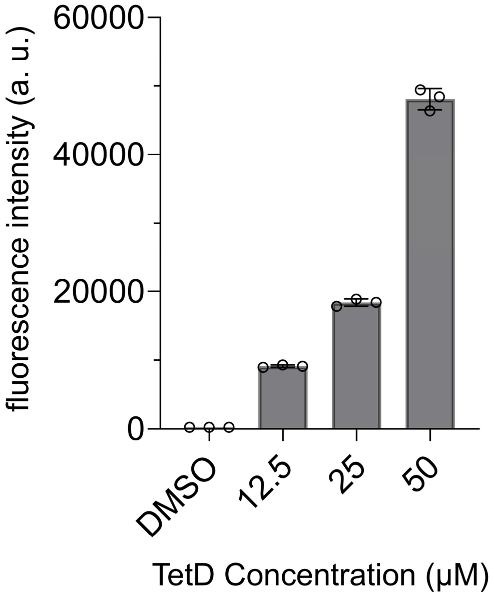


**Figure S2. TetD** concentration dependent labeling of *Msm*. Mid-log phase *Msm* cells were incubated with varying concentrations of **TetD** for 18 h and then treated with **Fl-az**. Data are represented as mean ± SD (n= 3).

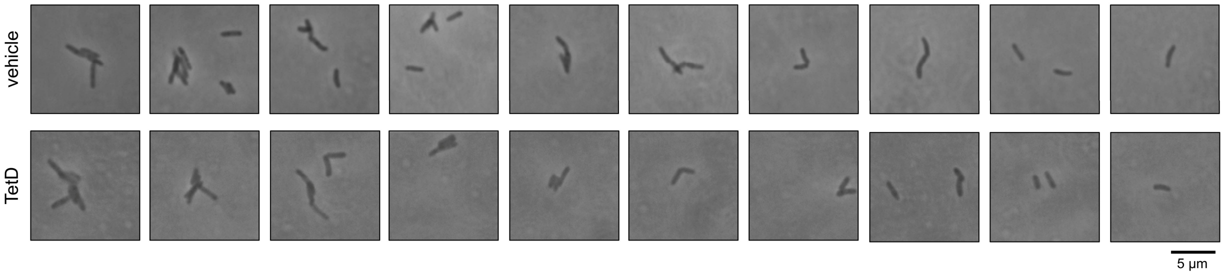


**Figure S3.** Phase-contrast microscopy of *Mtb* cells grown in 7H9/OADC +/- 25 µM **TetD** for 96 h. Fixed bacteria were imaged using conventional microscopy (Zeiss Axioscope A1 with 100x objectives).

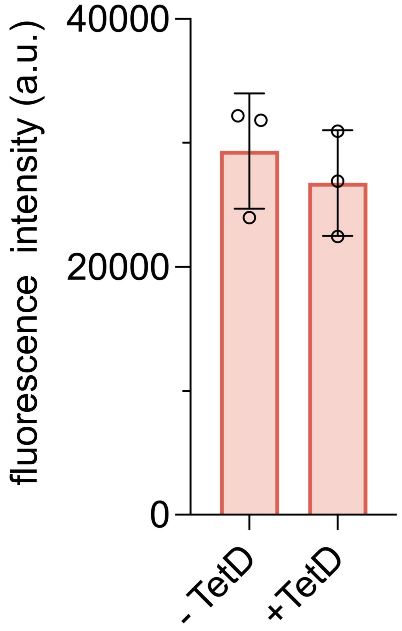


**Figure S4.** Ethidium Bromide (EtBr) based mycomembrane integrity analysis of *Msm* cells labeled with 25 µM **TetD**. Following labeling, cells were treated with EtBr and cellular fluorescence intensities reported here were measured after 60 minutes of incubation. Treatment with DMSO (**-TetD**) served as the control. Data are represented as mean ± SD (n= 3).

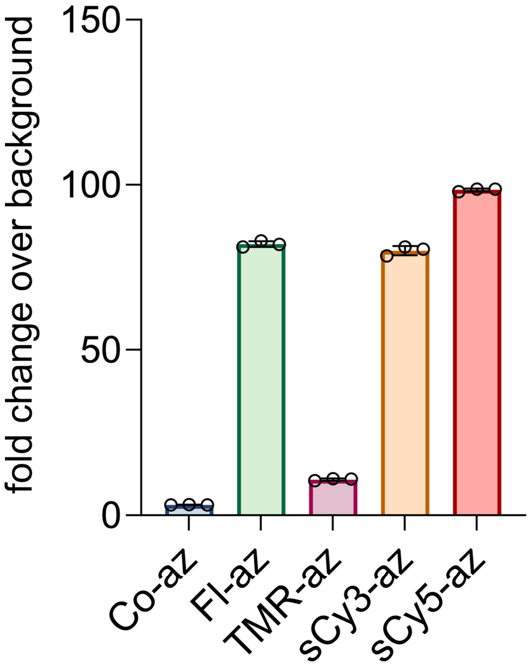


**Figure S5.** Signal-to-noise ratios observed upon treatment of **TetD** labeled *Msm* cells with various azide-tagged fluorophores for a duration of 1 h. Data are represented as mean ± SD (n= 3).

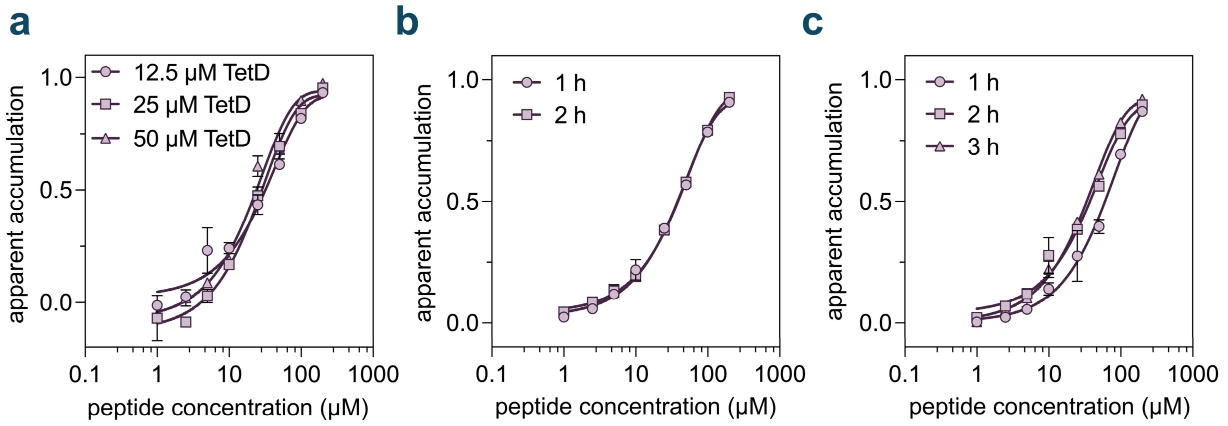


**Figure S6. (a)** Dose-response analysis showing the apparent accumulation of **Lin0** across the mycomembrane in *Msm* labeled with different concentrations of **TetD**. **(b)** Dose-response analysis showing the apparent accumulation of **Lin0** across the mycomembrane in *Msm* with varying **Fl-az** treatment durations. **(c)** Dose-response analysis showing the apparent accumulation of **Lin0** across the mycomembrane in *Msm* with varying **Lin0** treatment durations. All dose-response analyses involving **Lin0** were performed with a 2 h incubation period with the peptide. Data are represented as mean ± SD (n= 3). For dose-response curves, Boltzmann sigmoidal curves were fitted to the data using GraphPad Prism.

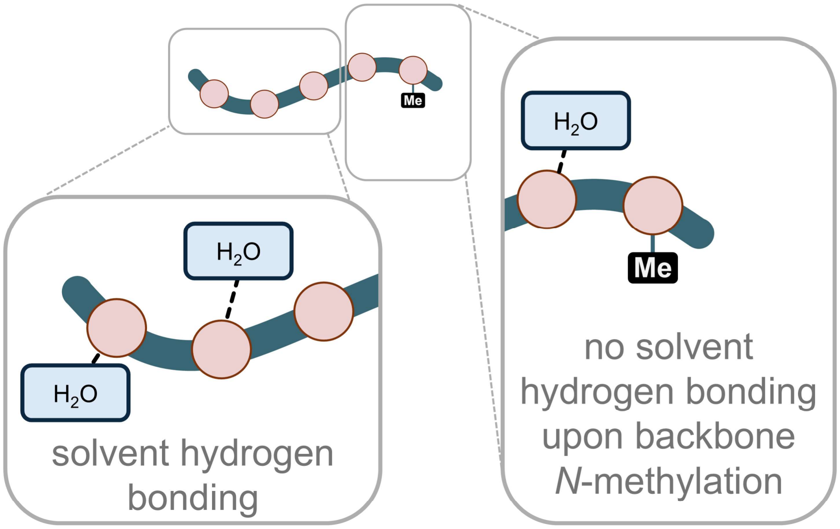


**Figure S7.** Schematic illustration of an *N*-methylated peptide interacting with the surrounding water molecules in an aqueous solution. Hydrogen bonds are indicated by dashed lines. Installation of a methyl group on a backbone amide eliminates the possibility of hydrogen bonding with the solvent for that specific amide, thereby potentially reducing membrane permeation associated desolvation.

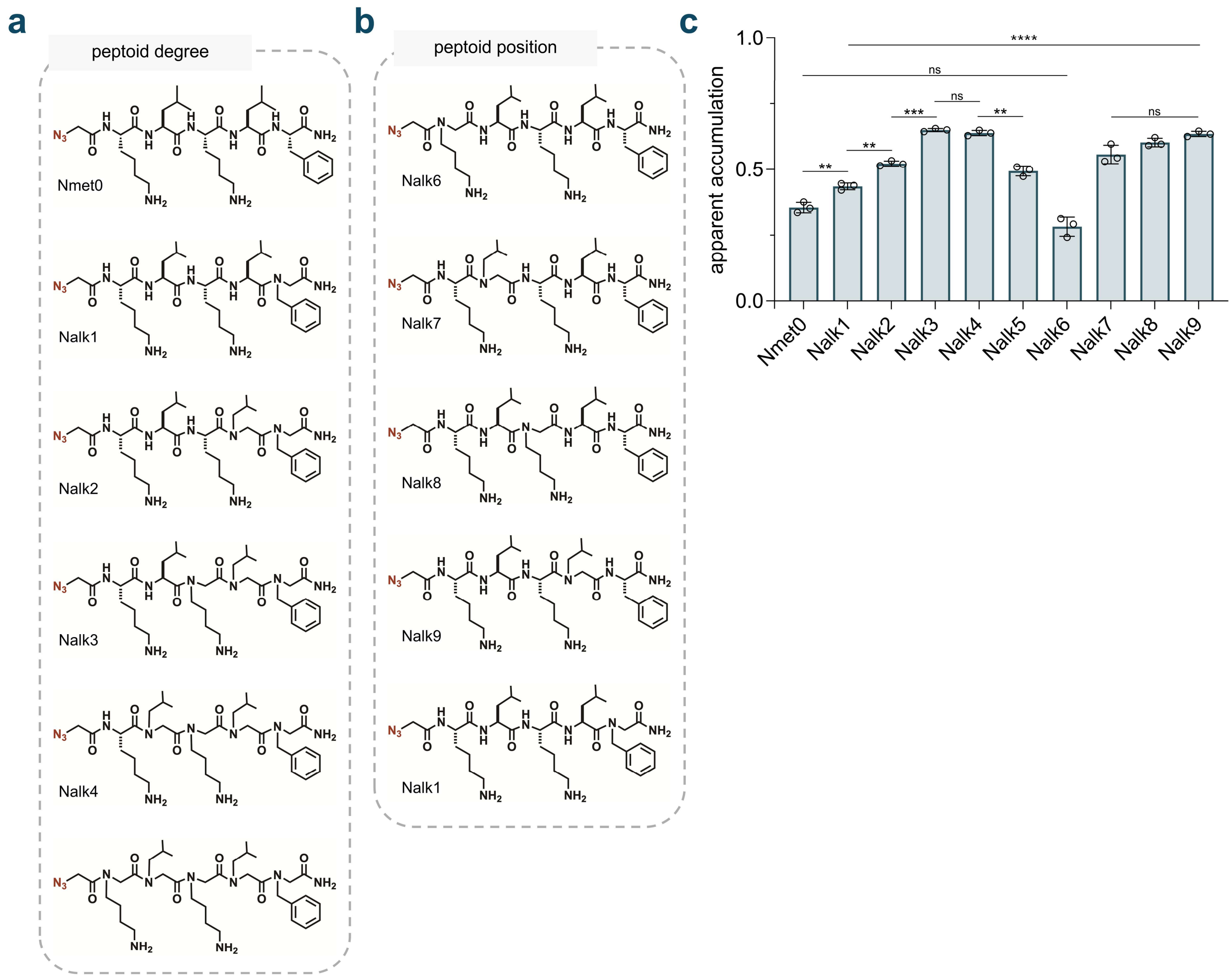


**Figure S8. (a)** Chemical structures of the *N*-alkylation (peptoid) sub-library with varying degrees of backbone *N*-alkylation (**Nalk1**-**5**) and the non-alkylated control (**Nmet0**). **(b)** Chemical structures of the *N*-alkylation sub-library with varying positions of backbone *N*-alkylation (**Nalk6**-**9** and **Nalk1**). **(c)** Apparent accumulation of the *N*-alkylation library across the mycomembrane in *Msm* following 1 h of treatment with 50 μM compound. Data are represented as mean ± SD (n= 3). P-values were determined by a two-tailed t-test (ns = non-significant, * $p < 0.1$, ** $p < 0.01$, *** $p < 0.001$, **** $p < 0.0001$).

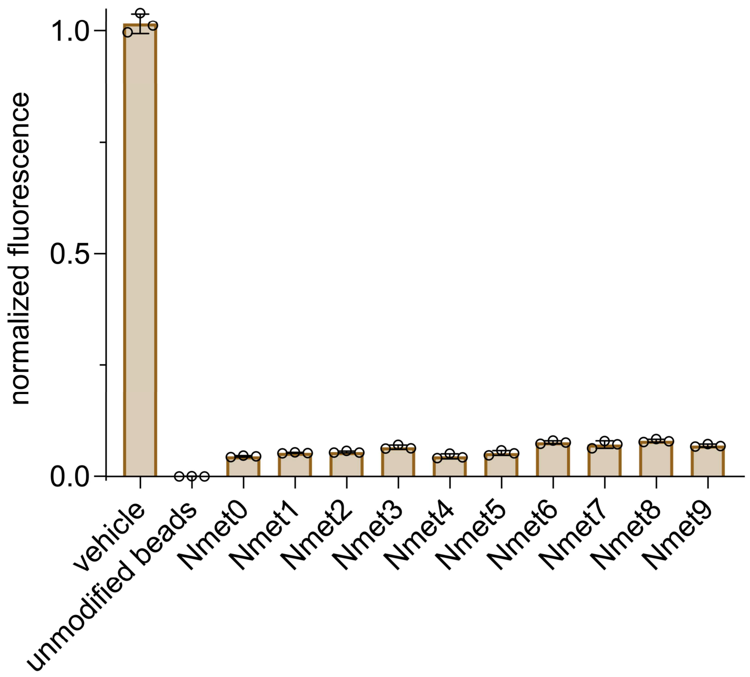


**Figure S9.** DBCO-modified polystyrene bead reactivity assay performed with the *N*-methylation sub-library, at a concentration of 50 μM over 1 h. Data are represented as mean ± SD (n= 3).

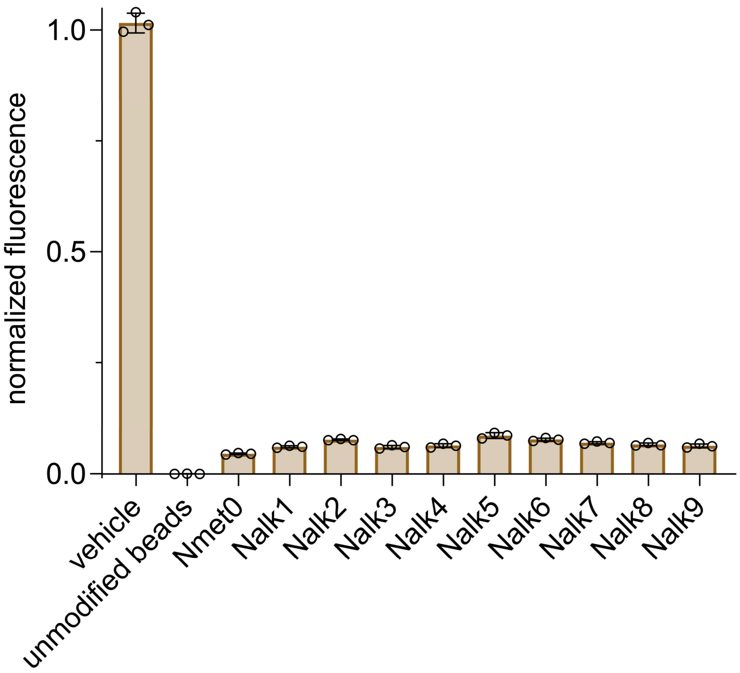


**Figure S10.** DBCO-modified polystyrene bead reactivity assay performed with the *N*-alkylation sub-library, at a concentration of 50 μM over 1 h. Data are represented as mean ± SD (n= 3).

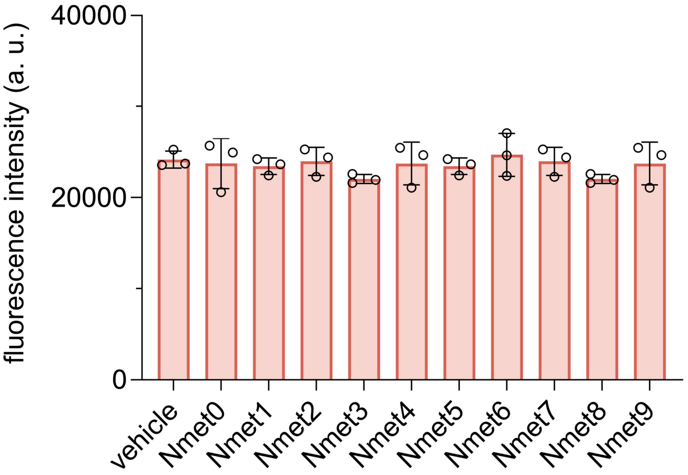


**Figure S11.** Ethidium Bromide (EtBr) based mycomembrane integrity analysis of *Msm* cells labeled with **TetD** and treated with the *N*-methylation sub-library at 50 μM for 1 h. Following treatment, cells were incubated with EtBr and cellular fluorescence intensities reported here were measured after 1 h of incubation. Data are represented as mean ± SD (n= 3).

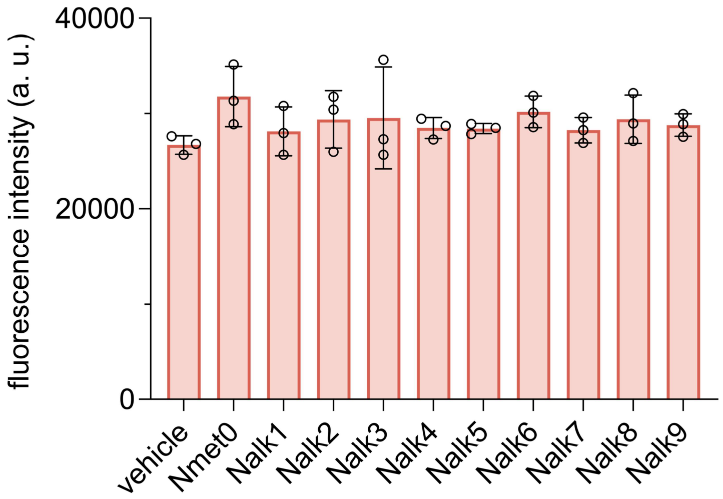


**Figure S12.** Ethidium Bromide (EtBr) based mycomembrane integrity analysis of *Msm* cells labeled with **TetD** and treated with the *N*-alkylation sub-library at 50 μM for 1 h. Following treatment, cells were incubated with EtBr and cellular fluorescence intensities reported here were measured after 1 h of incubation. Data are represented as mean ± SD (n= 3).

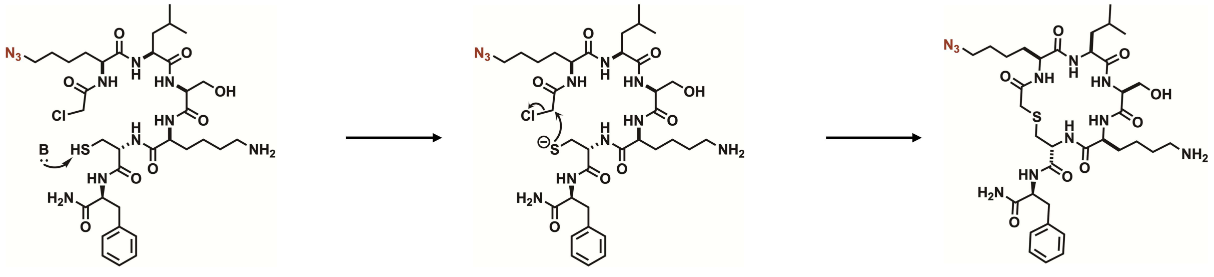

**Figure S13.** Scheme illustrating the cyclization chemistry adopted for the synthesis of the thioether based macrocyclic peptides. Macrocyclic peptide scaffolds with an *N*-terminal chloroacetyl group and a cysteine chain were then dissolved in a mixture of $H_2O/CH_3CN$ (1:1, v/v) to an approximate concentration of ~1.5 mM. This was followed by the addition of 0.5 M $NH_4HCO_3$ (pH 8.5) to a final concentration of 0.05 M and stirring overnight.

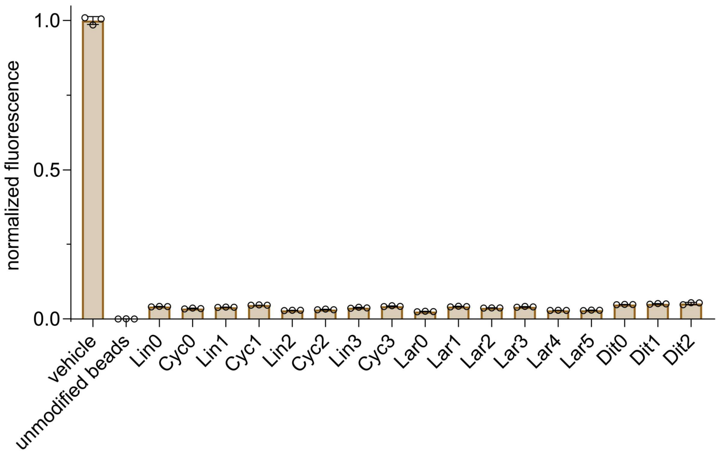


**Figure S14.** DBCO-modified polystyrene bead reactivity assay performed with the macrocyclization library at a concentration of 50 μM over 2 h. Data are represented as mean ± SD (n= 3).

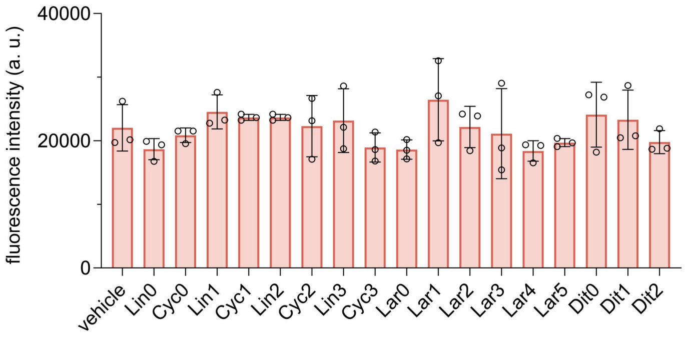


**Figure S15.** Ethidium Bromide (EtBr) based mycomembrane integrity analysis of *Msm* cells labeled with **TetD** and treated with the macrocyclization library at 50 μM for 2 h. Following treatment, cells were incubated with EtBr and cellular fluorescence intensities reported here were measured after 1 h of incubation. Data are represented as mean ± SD (n= 3).

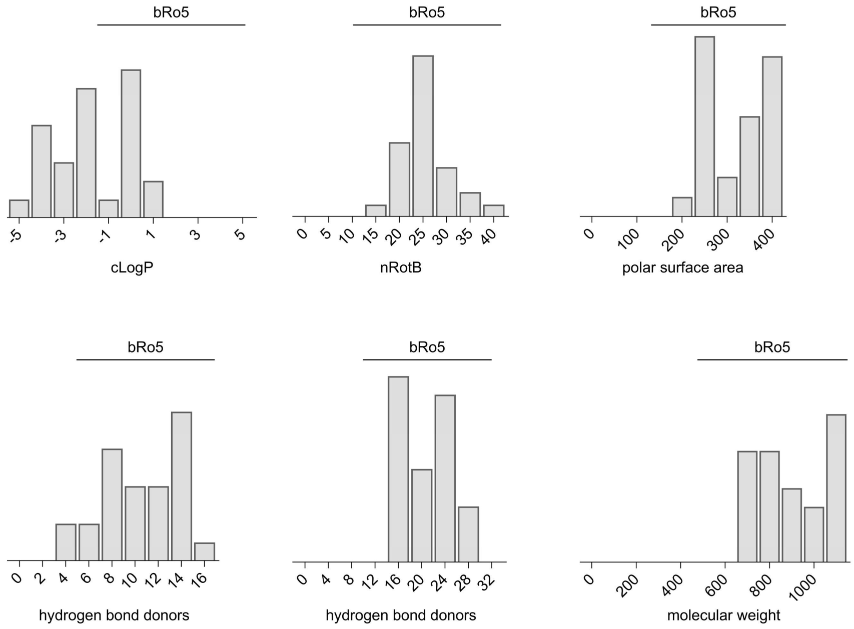


**Figure S16.** Frequency distributions of key physicochemical properties of the *N*-methylation and macrocyclization peptide libraries, including cLogP, number of rotatable bonds (nRotBs), polar surface area, number of hydrogen bond donors, number of hydrogen bond acceptors, and molecular weight. The chemical space beyond the rule-of-five (bRo5) is highlighted.

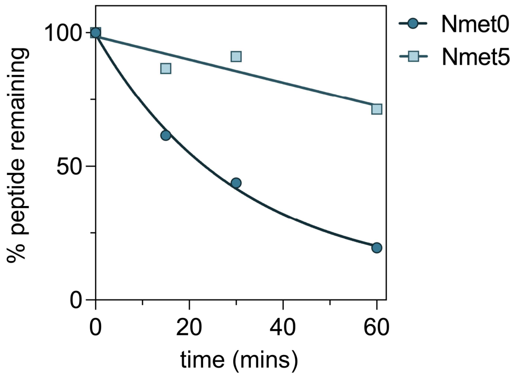


**Figure S17.** Comparison of serum stability of **Nmet0** and **Nmet5** incubated in mouse serum at 37 °C over 1 h. For serum stability analyses, one-phase decay curves were fitted to the data using GraphPad Prism.

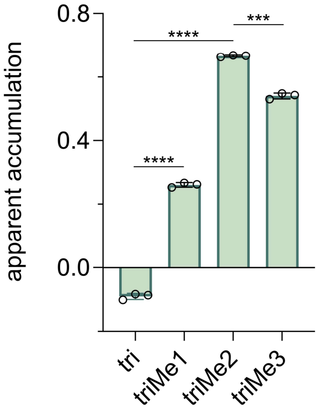


**Figure S18.** Apparent accumulation of the *N*-methylated tridecaptin series across the mycomembrane in *Msm* following 1 h of treatment with 50 µM compound. Data are represented as mean ± SD (n= 3). P-values were determined by a two-tailed t-test (ns = not significant, *p < 0.1, **p < 0.01, ***p < 0.001, ****p < 0.0001).

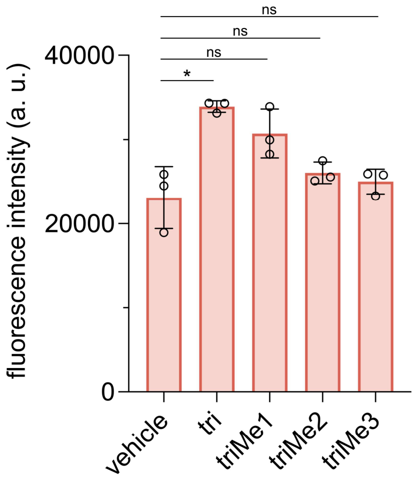


**Figure S19.** Ethidium Bromide (EtBr) based mycomembrane integrity analysis of *Msm* cells labeled with **TetD** and treated with the *N*-methylated tridecaptin series at 50 μM for 1 h. Following treatment, cells were incubated with EtBr and cellular fluorescence intensities reported here were measured after 1 h of incubation. Data are represented as mean ± SD (n= 3). P-values were determined by a two-tailed t-test (ns = not significant, *p < 0.1, **p < 0.01, ***p < 0.001, ****p < 0.0001).

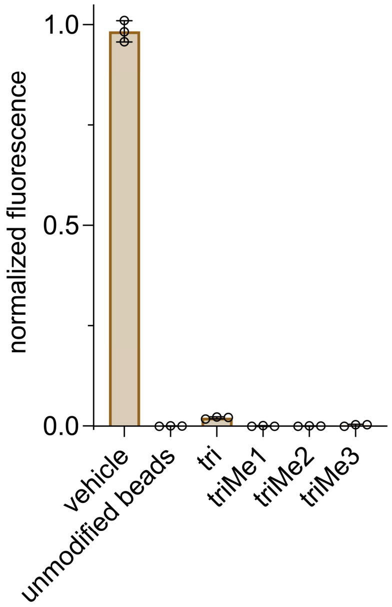


**Figure S20.** DBCO-modified polystyrene bead reactivity assay performed with the *N*-methylated tridecaptin series, at a concentration of 50 μM over 1 h. Data are represented as mean ± SD (n= 3).

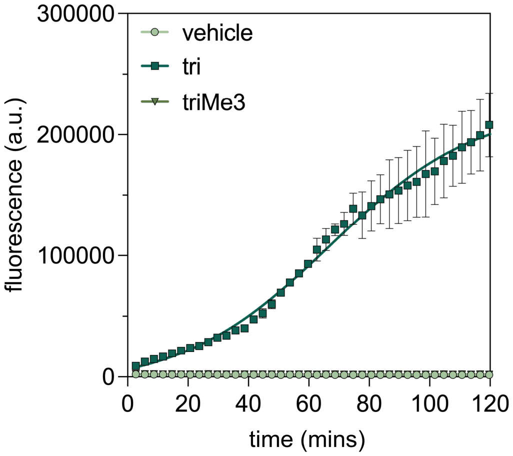


**Figure S21.** ThT fluorescence assay performed with the tridecaptin peptide analogs **triMe3** and **tri**. 50 μM of each peptide was incubated with 10 μM ThT for 2 h in PBST (pH 7.4, 37 °C). Fluorescence intensities were measured every 3 min for 2 h. Data are represented as mean ± SD (n= 3).

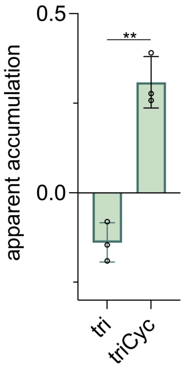


**Figure S22.** Apparent accumulation of the macrocyclic tridecaptin analog **triCyc** in comparison to the parent tridecaptin analog **tri**, across the mycomembrane in *Msm* following 1 h of treatment with 50 μM compound. Data are represented as mean ± SD (n= 3). P-values were determined by a two-tailed t-test (ns = not significant, *p < 0.1, **p < 0.01, ***p < 0.001, ****p < 0.0001).

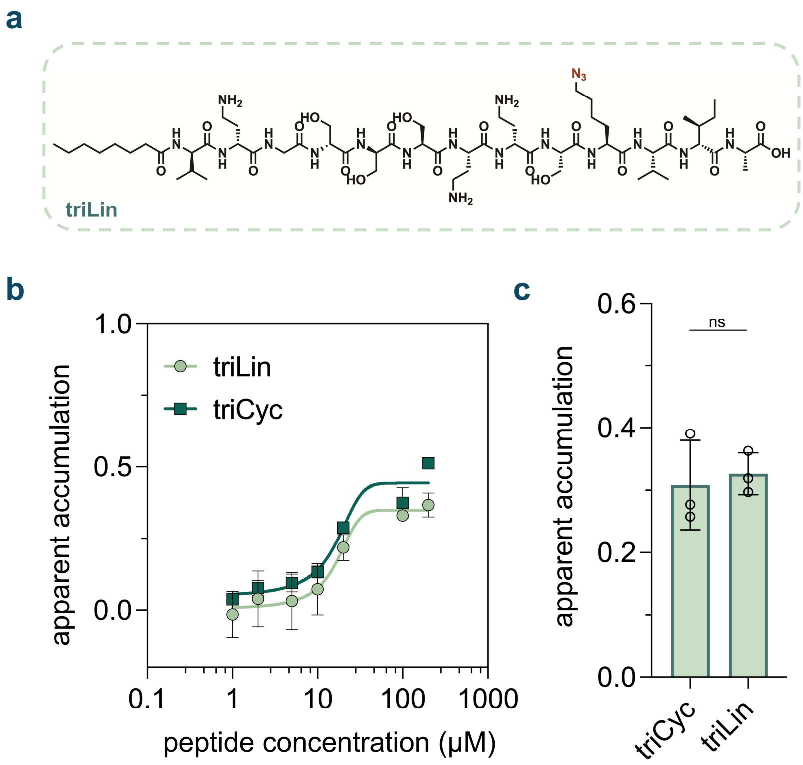


**Figure S23. (a)** Chemical structure of the linear tridecaptin analog **triLin** **(b)** Dose-response analysis showing the apparent accumulation of the macrocyclic and linear tridecaptin analogs **triCyc** and **triLin**, across the mycomembrane in *Msm*. Analysis was performed with a 1 h incubation period with the peptides. **(c)** Apparent accumulation of the macrocyclic and linear tridecaptin analogs **triCyc** and **triLin**, across the mycomembrane in *Msm* following 1 h of treatment with 50 µM compound. Data are represented as mean ± SD (n= 3). P-values were determined by a two-tailed t-test (ns = not significant, *p < 0.1, **p < 0.01, ***p < 0.001, ****p < 0.0001). For dose-response curves, Boltzmann sigmoidal curves were fitted to the data using GraphPad Prism.

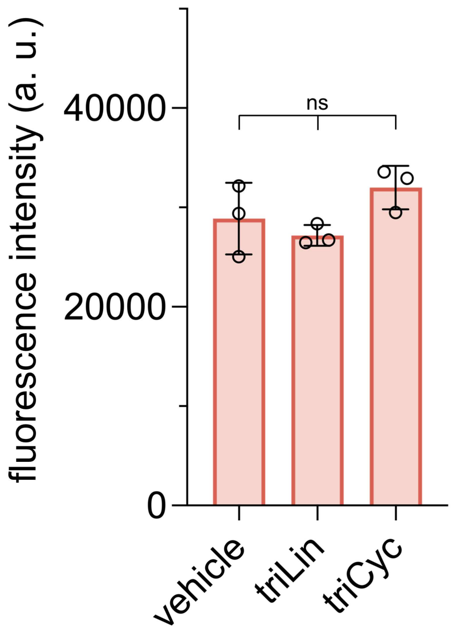


**Figure S24.** Ethidium Bromide (EtBr) based mycomembrane integrity analysis of *Msm* cells labeled with **TetD** and treated with the macrocyclic and linear tridecaptin analogs **triCyc** and **triLin** at 50 μM for 1 h. Following treatment, cells were incubated with EtBr and cellular fluorescence intensities reported here were measured after 1 h of incubation. Data are represented as mean ± SD (n= 3). P-values were determined by a two-tailed t-test (ns = not significant, *p < 0.1, **p < 0.01, ***p < 0.001, ****p < 0.0001).

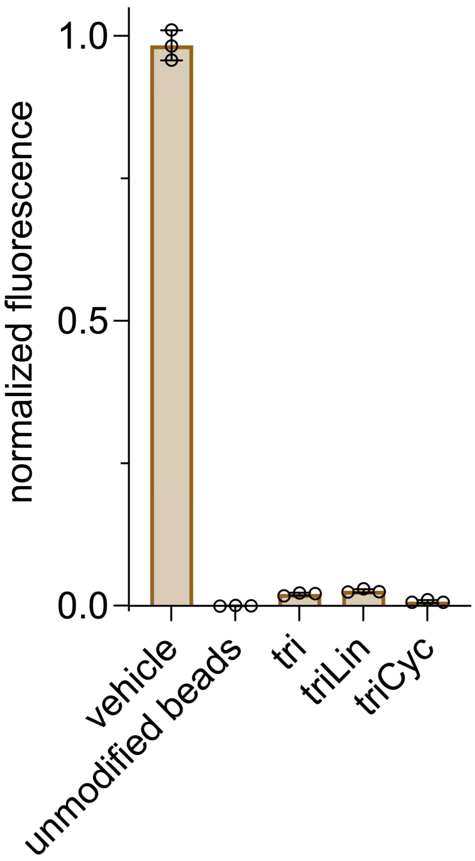


**Figure S25.** DBCO-modified polystyrene bead reactivity assay performed with the macrocyclic and linear tridecaptin analogs **triCyc** and **triLin**, at a concentration of 50 µM over 1 h. Data are represented as mean ± SD (n= 3).

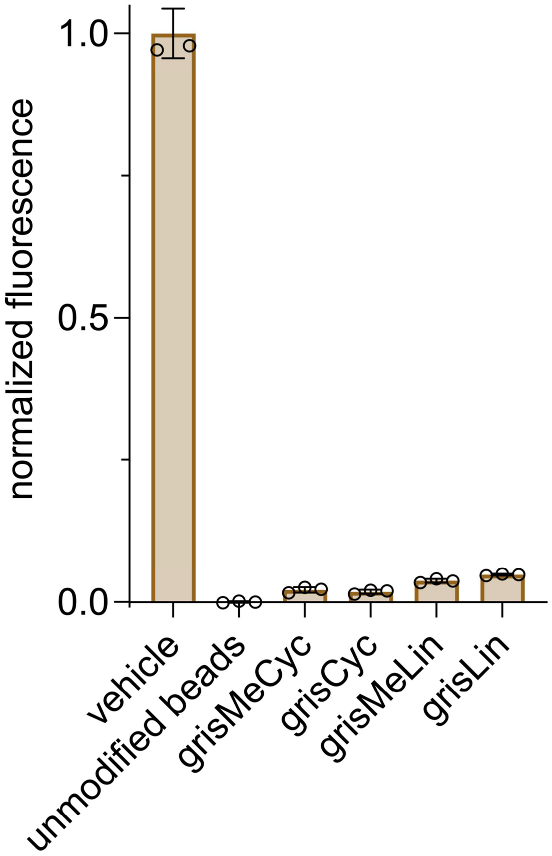


**Figure S26.** DBCO-modified polystyrene bead reactivity assay performed with the griselimycin series, at a concentration of 50 μM over 6 h. Data are represented as mean ± SD (n= 3).

#### EXPERIMENTAL PROCEDURES

##### Materials

| REAGENTS | VENDOR SOURCE | CATALOG # |
|---|---|---|
| **Reagents for biological methods** | | |
| Middlebrook 7H9 media | VWR | 90003-876 |
| Oleic Albumin Dextrose Catalase (OADC) | BD | BD 212351 |
| Catalase from bovine liver | Millipore Sigma | C1345-1G |
| D(+)-Glucose, anhydrous (Dextrose) | Chem Impex | 00805 |
| Bovine serum albumin fraction V | Millipore Sigma | 10735078001 |
| Glycerol | Millipore Sigma | G7893 |
| Tween 80 | VWR | 97061-674 |
| Formaldehyde solution | Millipore Sigma | 252549 |
| 16% Paraformaldehyde | Ted Pella | NC0528893 |
| 6-azido-fluorescein **(Fl-az)** | Lumiprobe | D5130 |
| TAMRA azide, 6-isomer **(TMR-az)** | Lumiprobe | D8130 |
| sulfo-Cyanine3 azide **(sCy3-az)** | Lumiprobe | B1330 |
| sulfo-Cyanine5 azide (**sCy5-az)** | Lumiprobe | C3330 |
| 3-azido-7-hydroxy coumarin (**Co-az**) | Millipore Sigma | 909513-5MG |
| CalFluor 647 azide | Vector laboratories | CCT-1372-5 |
| Amino terminated polystyrene beads | Spherotech | AP-30-10 |
| Trichloroacetic acid solution | Millipore Sigma | T0699 |
| Ethidium bromide solution (10 mg/mL), Biotechnology Grade | VWR | 97064-970 |
| Protease inhibitor | Millipore Sigma | 11836153001 |
| SDS | Millipore Sigma | 436143 |
| Proteinase K | Millipore Sigma | SAE0009-1G |
| Mutanolysin | Millipore Sigma | M9901-10KU |
| Lysozyme | MP Biomedicals | 100834 |
| L- leucine | Millipore Sigma | L8000 |
| D-Pantothenic acid hemicalcium salt | Millipore Sigma | 21210 |
| Mouse serum | Atlanta Biologicals | S18193 |
| **Reagents for the synthesis and characterization of test molecules** | | |
| α-*N*-Fmoc-amino acids | Chem Impex | Various |

| | | |
|---|---|---|
| *N*-methylated amino acids | Chem Impex | Various |
| 2-azido-acetic-acid | Chem Impex | 35113 |
| 2-Chlorotrityl chloride resin (1.0-2.0 meq/g, 100-200 mesh | Chem Impex | 03498 |
| Rink amide resin, (0.3-0.6 meq/g, 100-200 mesh) | Chem Impex | 12662 |
| *N*, *N'*-Diisopropylethylamine (DIEA) | Chem Impex | 00141 |
| Hexafluorophosphate Benzotriazole Tetramethyl Uronium (HBTU) | Chem Impex | 02011 |
| *N*, *N'*-Diisopropylcarbodiimide (DIC) | Chem Impex | 00110 |
| Ethyl Cyano(hydroxyimino)acetate (Oxyma) | TCI Chemicals | E0847 |
| *N, N*-Dimethylformamide (DMF) ACS-grade | Millipore Sigma | 319937 |
| Dichloromethane ACS-grade | Millipore Sigma | D65100 |
| Methanol ACS-grade and HPLC-grade | Millipore Sigma | 179337, 34860 |
| Acetonitrile ACS-grade and HPLC-grade | Millipore Sigma | 360457, 34851 |
| Piperidine | Millipore Sigma | 104094 |
| Trifluoroacetic acid (TFA) HPLC grade | Millipore Sigma | 302031 |
| Trifluoroacetic acid (TFA) reagent-grade | Chem Impex | 00289 |
| Dibenzocyclooctyne- DBCO-NHS ester | Broadpharm | BP-22231 |
| Ammonium Bicarbonate | Fisher Scientific | A643-500 |
| Ellman's Reagent (5,5-dithio-bis-(2-nitrobenzoic acid)) | Thermo Scientific | 22582 |
| 1,4-Bis(bromomethyl)benzene | Ambeed | A575067 |
| TIPS | | |
| **General materials and equipment** | | |
| 96-well clear conical bottom deep well plates | VWR | 76210-524 |
| 96-well clear untreated round bottom plates | VWR | 82050-622 |
| 96-well black half area black flat bottom plates | Millipore Sigma | CLS3694 |
| Amicon® Ultra Centrifugal Filter, 3 kDa MWCO | Millipore Sigma | UFC5003 |

#### Biological Methods

**Mycobacteria cell culture.** *Mycobacterium smegmatis* strain mc$^2$155 *(Msm)* was grown in 7H9 media with 0.5% glycerol, 0.05% tween 80, and 1x ADC enrichment. 10x ADC was made with 5 g bovine serum albumin, 2 g dextrose, 3 mg catalase in 100 mL autoclaved milliQ $H_2O$. Glycerol stocks were made using stationary phase cells in 30% glycerol and aliquots were stored at - 80 ºC. The double auxotroph *M. tuberculosis* (*Mtb*) mc$^2$6206 strain (H37Rv Δ*panCD* Δ*leuCD*[1] was provided by Dr. William Jacobs and was cultured in Middlebrook 7H9 supplemented with 0.5% glycerol, 10% Middlebrook Oleic Albumin Dextrose Catalase (OADC; BD), and either 0.05% Tyloxapol, for strain maintenance, or 0.05% Tween-80, for experimental manipulation. Growth media was additionally supplemented with 50 µg/mL L-leucine and 50 µg/mL pantothenic acid. Bacterial stocks were prepared by directly freezing *Mtb* culture (OD600 ~0.5) at -80 °C.

**DBCO-tetrapeptide (TetD) based labeling of *Msm*.** *Msm* cells were inoculated from the glycerol stock into the 7H9 media in standard culture tubes and grown for 24 hours at 37 ºC until they reached an OD of 0.5-0.6. **TetD** probe stock solution was then added to the media to yield a final concentration of 25 µM and the cells were grown overnight until stationary phase. Notably, blank cells (-DBCO) remained untreated in comparison to the labeled cells at this point. The cells were harvested the next day and centrifuged for 2 min at 3000×g and then washed with phosphate-buffered saline containing 0.05% tween 80 (PBST) twice and resuspended in PBST to yield **TetD**-modified *Msm*. To a 96-well plate, 90 µL of labeled cells were added in triplicate. For the blank wells, 90 µL of unlabeled cells were added in triplicate. 10 µL of 500 µM **Fl-az** (50 µM final) was then added to each well followed by incubation at 37 ºC for 1 h. For the experiment testing different azide-tagged fluorophores, 10 µL of 500 µM **Fl-az**, 3-azido-7-hydroxy coumarin (**Co-az)**, azido-tetramethyl rhodamine (**TMR-az**), azido-sulfo Cy3 (**sCy3-az**) or azido-sulfo Cy5 (**sCy5-az**) were added to **TetD**-treated *Msm* cells. The cells were then centrifuged down, and the pellets were washed with 100 µL PBST twice and fixed with 100 µL 4% formaldehyde for 15 min. The samples were then analyzed by AttuneTM NxT Acoustic Focusing Cytometer.

**DBCO-tetrapeptide (TetD) based labeling of *Mtb***. *Mtb* cells were diluted to $OD_{600}$ ~0.015 and incubated in 7H9 (supplemented with Oleic Albumin Dextrose Catalase, 0.05% Tween-80, 0.5% glycerol) +/- 25 µM **TetD** for 4 days (96 h) until they reached $OD_{600}$ ~0.25. Cells were then harvested and washed twice with the same medium and incubated with 17 µM of the fluorogenic label CalFluor647-azide in 7H9 for 1 h at 37 °C shaking. The fluorophore was removed by spinning down and removal of the supernatant. The bacteria were then fixed with fresh prepared 4% paraformaldehyde (PFA) in PBS for at least 2 h at room temperature. The bacteria were then resuspended in physiological solution (0.9% NaCl in water) and analyzed by BD DUAL LSRFortessa.

**Accumulation assay (PAC-MAN) across the *Msm* mycomembrane.** To a 96-well plate, 90 µL of **TetD**-modified *Msm* cells were added in triplicate. Next, 5 µL of 1 mM azide-tagged peptide stock solutions were added to yield a final concentration of 50 μM. The volume in each well was made up to 100 μL using PBS. To the blank and PBS control wells, 10 μL of PBS was added at this point. For the blank wells, 90 μL of unlabeled cells were added in triplicate. The plates were then incubated at 37 ºC with shaking. For the *N*-methylated library, *N*-alkylated library, and tridecaptin series, the treatment duration was 1 h. For the macrocyclization library, the treatment duration was 2 h. For the griselimycin series, the treatment duration was 6 h. Importantly, for dose-response analyses, varying concentrations of azide-tagged peptide were used, and an appropriate amount of stock solution was added accordingly, for each concentration. The cells were centrifuged for 2 min at 3000×g, and the supernatant was discarded. The pellets were then washed with 100 μL PBST twice and finally resuspended in 90 μL of PBST. 10 μL of 500 μM **Fl-az** (50 μM final) was then added to each well followed by incubation at 37 ºC for 1h. The cells were then centrifuged again, and the pellets were washed with 100 μL PBST twice and fixed with 100 μL 4% formaldehyde in PBS for 15 min. The samples were then analyzed by AttuneTM NxT Acoustic Focusing Cytometer.

**Accumulation assay (PAC-MAN) across the *Mtb* mycomembrane**. *Mtb* cells were diluted to $OD_{600}$ ~0.015 and incubated in 7H9 (supplemented with Oleic Albumin Dextrose Catalase, 0.05% Tween-80, 0.5% glycerol) +/- 25 μM **TetD** for 4 days until $OD_{600}$ ~0.25, washed twice with the same medium then incubated with 50 μM of azide-peptides in 7H9 in 96-well plates at 37 °C with shaking. For the *N*-methylated library, the treatment duration was 1 h. For the macrocyclization library, the treatment duration was 2 h. Azide-peptides were removed by spinning down and removal of the supernatant. The bacteria were then incubated with 17 μM of the fluorogenic label CalFluor647-azide in 7H9 for 1 hour at 37 °C while shaking. The fluorophore was removed by spinning down and removal of the supernatant. The bacteria were then fixed with fresh prepared 4% paraformaldehyde (PFA) in PBS for at least 2 h at room temperature. Then bacteria were resuspended in physiological solution (0.9% NaCl in water) and analyzed by BD DUAL LSRFortessa.

**Isolation of peptidoglycan and muropeptides**. *Msm* cells were inoculated from the glycerol stock into the 7H9 media in standard culture tubes and grown for 24 hours at 37 ºC until they reached an OD of 0.5-0.6. **TetD** probe stock solution was then added to the media to yield a final concentration of 25 μM and the cells were grown overnight until stationary phase. The cells were harvested the next day and centrifuged for 2 min at 3000×g and then washed with PBST twice and resuspended in PBST to yield **TetD**-modified *Msm*. To a 96-well plate, 90 μL of labeled cells were added in triplicate. 10 μL of 500μM **Fl-az** (50 μM final) was then added to each well followed by incubation at 37 °C for 1 h. The cells were then centrifuged down, and the pellets were washed with 100 μL PBST twice. The cell pellets were then resuspended in 200 μL 10 mM $NH_4HCO_3$ containing protease inhibitor and sonicated in an ultrasonic bath sonicator for 30 min. 10 μg/mL each of DNase and RNase were then added to each well directly after sonication and the plate was placed at 4 °C for 1 h. The cell wall-enriched fraction was then collected by centrifugation at 2700 g for 10 min. The pellets were treated with 200 μL PBS containing 2% sodium dodecyl sulfate (SDS) at 50 °C for 1 h while shaking at 250 rpm. The suspension was spun down at 2700 g for 10 min. This treatment was repeated twice. Then the resulting pellet was resuspended in 200 µL PBS containing 1% SDS and 0.1 mg/mL proteinase K each well, incubated at 37 °C for 1 h with shaking at 250 rpm. The suspension was then heated in boiling water for 1 h and spun down at 2700 g for 10 min. The supernatant was discarded and the 1% SDS extraction step was repeated twice. The pellet was then washed twice with PBS and four times with deionized water to give the mycolyl-arabinogalactan-peptidoglycan Complex (MAPc). The MAPc samples were then resuspended in 200 µL methanol containing 0.5% KOH and incubated at 37 °C for 4 days with shaking at 250 rpm. The mixture was then washed twice with methanol and twice with diethyl ether and air-dried to give arabinogalactan-peptidoglycan (AGPG). The resulting AGPG was resuspended in 200 µL deionized water and digested with 0.05 N $H_2SO_4$ at 37 °C for 5 days with shaking at 250 rpm. The samples were then washed four times with deionized water to give insoluble peptidoglycan (PG). The isolated sacculi from each well in each group was then combined and lyophilized, respectively. The lyophilized sacculi was resuspended in 0.4 mL 10 mM sodium acetate (pH 5) containing 25 µg/mL mutanolysin and lysozyme and digested at 37 °C over 20 h. The mixture was then spun down, and the supernatant was filtered with a 10 kDa MWCO filter and lyophilized. The lyophilized muropeptide samples were then dissolved in 30 µL milliQ water and an aliquot was analyzed *via* a C18(2) column (Luna 5 µm 100Å 250 x 4.6 mm) on an Agilent LC-QTOF system (Agilent 1260 Infinity II Prime LC with Agilent 6545B QTOF). Muropeptide samples were eluted with a 2 to 30% linear gradient of $H_2O/CH_3CN$ (0.5% formic acid) at 0.4 mL/min. The muropeptides were analyzed by MS using MassHunter software.

**Ethidium bromide (EtBr) whole-cell accumulation assay.** To a 96-well plate, 90 µL of **TetD** -modified *Msm* cells were added in triplicate. Next, 5 µL of 1 mM peptide stock solutions were added to yield a final concentration of 50 µM. The volume in each well was made up to 100 µL using PBS. To the blank and PBS control wells, 10 µL of PBS was added at this point. For the blank wells, 90 µL of unlabeled cells were added in triplicate. The plates were then incubated at 37 ºC for the same duration as the corresponding PAC-MAN assay incubation duration for that library. The cells were centrifuged for 2 min at 3000× g, and the supernatant was discarded. The pellets were then washed with 100 µL PBST twice and finally resuspended in 100 µL of PBST. 95 µL of these cells were then transferred to corresponding wells in a 96-well, black, flat-bottomed plate. To each well, 5 µL of 100 µM EtBr (5 µM final) was added. The fluorescent intensities were measured by a Biotek Synergy H1 microplate reader 60 m post-treatment with EtBr, with continuous orbital shaking. Wavelengths: ethidium bromide, excitation 530 nm/emission 590 nm.

**DBCO-modified polystyrene bead reactivity assay.** For functionalization with DBCO**,** 100 µL of amino functionalized polystyrene beads (5% w/v, 5 mg) were centrifuged at 21000 g for 10 min in a 1.7 mL eppendorf tube and washed with 1 mL deionized water. The beads were then resuspended in 1 mL 20 mM sodium borate buffer (pH 9) containing 1 µg/mL DBCO-NHS and incubated at 37 °C for 2 h with shaking. The resulting beads were centrifuged at 21000 g for 10 min and then washed and resuspended in sodium borate buffer. For capping the remaining unreacted amine sites on the beads, 20 µL of acetic anhydride was added to the bead suspension and the beads were then incubated at 37 °C for 2 h with shaking. The resulting beads were then centrifuged and washed twice with 1 mL PBS and resuspended in 1 mL PBS for further use. For the reactivity assay, the DBCO modified polystyrene beads were diluted 1:1 in PBS prior to the assay. A 96-well plate was loaded with 95 µL of test molecules (for final concentration of 50 µM) to which 5 µL of the beads was added to each well. The plate was incubated in 37 °C for 2 h and centrifuged. The supernatant was decanted and 100 µL 50 µM **Fl-az** was added to each well. The plate was then incubated at 37 °C for 1h and centrifuged at 2700 g for 10 min. The supernatant was decanted, and the beads were washed with 200 µL PBS and then resuspended in 200 µL PBS. The samples were then analyzed by AttuneTM NxT Acoustic Focusing Cytometer. Fluorescence intensities from unmodified beads were regarded as 0% and fluorescence intensities from DBCO modified beads treated with DMSO (vehicle) were regarded as 100% for the normalization of raw data. Normalization of fluorescence intensities from treatments with azido-peptides were then performed accordingly.

**Calculation of physiochemical properties of the test molecules.** DataWarrior software was used to calculate the molecular weight (MW), number of hydrogen bond donors (HBDs), number of hydrogen bond acceptors (HBAs), water/octanol partitioning coefficient (cLogP), polar surface area (PSA), and number of rotatable bonds (nRotBs) for each peptide.

**Hydrogen-deuterium exchange (HDX) studies.** NMR Data was acquired on either Bruker Ascend 400 MHz or Varian 600 MHz spectrophotometers. For both peptides, fresh solutions were made in deuterated solvents, namely, 70:30 [$CD_3CN$:$D_2O$] at ~10 mM concentration, prior to NMR studies. First water suppression 1H-NMR was acquired *via* a pre-saturation experiment using the Bruker Ascend 400 MHz spectrophotometer. Once the parameters for water pre-saturation were set, using the same parameters, all 2D-NMR (COSY, TOCSY, ROSEY) and 1D-NOESY spectra were acquired for the detailed assignment of each proton (especially amide NH) and its correlation with neighboring carbons and protons.[2] NMR data was processed and analyzed using the software MestreNova. Residual solvent signals from deuterated solvents were used as an internal standard with reference to tetramethylsilane (TMS) for defining chemical shifts. Chemical shifts are reported in ppm ($\delta$) and coupling constants (J) are reported in Hertz [Hz]. Time dependent spectra were continuously acquired after inserting sample into the instrument followed by locking and shimming, ~5 minutes past making fresh solutions. Area under the curve (integration) of each individual *NH* was used for determining the exchange rate, the initial integrations of these protons were normalized to non-exchangeable aromatic protons of the phenylalanine residue and were followed with time for reduced intensity by *N-H* to *N-D* exchange.

**Molecular dynamics** (**MD**) **simulation studies.** Three replicates of MD simulations were conducted for each of the **Lin0** and **Cyc0** peptide systems to compare their self-interactions. Each peptide was solvated using the TIP3P water model[3] and 0.15 M KCl was added to the system. The systems were prepared using the CHARMM scripts provided by CHARMM-GUI.[4] The CHARMM36m force field[5] was employed for the MD simulations. After energy minimization and 500 ps equilibration, a 100 ns production run was performed. All production MD simulations were performed for the NPT ensemble at 303.15 K and 1 atm using NAMD3. Long-range electrostatic interactions were treated using the particle mesh Ewald (PME) method[6] while Lennard-Jones (LJ) interactions were managed with a force-based switching function applied between 10 Å and 12 Å. Pressure and temperature were maintained using the Nosé–Hoover Langevin-piston method[7] with a piston period of 50 fs and a piston decay of 25 fs, coupled with Langevin temperature control with a friction coefficient of 1 $ps^{-1}$. A time step of 1 fs was chosen to capture detailed peptide dynamics. Hydrogen bonds between different residues within the same peptide were analyzed using the last 50 ns of the simulation.

**Serum stability assay.** Time course degradation in mouse serum was evaluated for peptides **Lin0**, **Cyc0**, **Nmet0** and **Nmet9**. Compounds were incubated in mouse serum at a concentration of 1.0 mM to a total of 200 μL at 37 °C. At each time point (0, 15, 30, 60 min), 40 μL of the solution was aliquoted and diluted in a 1:1 solution of $H_2O/CH_3CN$ (101 μL) containing trichloroacetic acid (9 μL) and 100 μM of internal standard Fmoc-L-phenylalanine. This mixture was incubated in ice for 5 min to precipitate serum proteins. The samples were then centrifuged at 21,000×g for 5 min. 140 μL of the supernatant was then transferred to a filter tube and centrifuged again at 21,000×g for 5 min to filter. 110 μL of the filtrate was then analyzed using Phenomenex Luna 5 μm C8(2) on RP-HPLC; gradient elution in $H_2O/CH_3CN$ with 0.1% TFA at 1 mL/min. Peptide identity was confirmed via peak collection and subsequent matrix-assisted laser desorption ionization time-of-flight (MALDI-TOF) mass spectroscopy (Shimadzu 8020). To eliminate variations in the overall signal intensity, area under the curve (AUC) for the peptides of interest at each time point was normalized to the AUC of the internal standard Fmoc-L-phenylalanine, the concentration of which was predetermined. These normalized AUC values were then further normalized to the "0 min" time point which was set to 100%. The percentage of peptide remaining was then plotted as a function of time.

**Minimum inhibitory concentration (MIC) assay.** *Msm* cells were inoculated from the glycerol stock to the according media and grown for 24 hours until 0.5-0.6 OD. Testing peptides were dissolved in the 7H9 growth media mentioned above and made into serial dilutions from 128 μM. 200 μL of each concentration was added to a deep well 96-well plate – with each concentration added in triplicates. The cells were diluted 100 times to OD of around 0.05 with fresh 7H9 growth media. 200 mL of diluted cells were added to the deep well plate each well and mixed well with the peptide solution. The plate was covered by an adhesive porous film and incubated at 37 °C with shaking at 250 rpm for 48 h. The contents from each well were then transferred to a flat-bottom 96-well plate for MIC value determination. MIC values were determined both visually and readings of OD (wells OD < 0.1) from a Biotek Synergy H1 microplate reader at 600 nm.

**Thioflavin T (ThT) fluorescence assay.** A solution of thioflavin T (ThT) was purchased as a DMSO stock. The concentration of ThT in the solution was determined using a UV-vis spectrophotometer ($\varepsilon$ = 36000 $M^{-1}$ $cm^{-1}$ at 412 nm). A working solution of 400 μM was freshly prepared in milliQ water from the DMSO stock before use. ThT fluorescence assays were conducted on a 96-well, black, flat-bottomed plate. 5-μL aliquots of the ThT working solution were transferred to each well on the 96-well plate in triplicate. Then, 10-μL aliquots of 1 mM solutions of each peptide were added to each of the wells, ensuring that the droplet did not touch the ThT solution. Final volume in each well was made up to 200 μL with PBST. The plate was then immediately inserted into a Biotek Synergy H1 microplate reader and incubated at 37 °C with shaking (262 rpm, fast speed, double orbital) and the fluorescence was thus monitored (444 nm excitation, 480 nm emission) every 3 min over 2 h.

#### SYNTHESIS AND CHARACTERIZATION OF TEST MOLECULES

**General procedure for the solid-phase synthesis of all peptides**. All peptides were prepared by standard Fmoc-based solid-phase chemistry using the appropriate resins. Briefly, to a 25 mL peptide synthesis vessel, an appropriate amount of resin was added, followed by 20% piperidine in *N, N*-Dimethylformamide (DMF, 15 mL). This was followed by shaking at room temperature for 30 min. The resin was then washed with methanol ($CH_3OH$) and dichloromethane (DCM) three times. After the last wash, 4 eqiv. of amino acid was added along with 4 eqiv. of ethyl cyanohydroxyiminoacetate (Oxyma) and 4 eqiv. of *N, N'*-Diisopropylcarbodiimide (DIC). The resin was shaken at room temperature for 2 h, then washed with $CH_3OH$/DCM. The remainder of the amino acids were coupled in the same manner. Peptides were cleaved from the resin using a TFA/TIPS/$H_2O$ mixture (95:2.5:2.5, v/v/v) (unless otherwise mentioned) shaking at room temperature for 2 h. The solution was filtered and concentrated prior to precipitation by the addition of cold diethyl ether to yield crude peptide. Crude peptides were purified by reverse-phased preparative high-performance liquid chromatography (RP-HPLC) equipped with Waters 1525 with 2489 UV/Visible Detector on a Phenomenex Luna 10 µm C8(2) 100 Å (250 x 21.2 mm) column using gradient elution with $H_2O$/MeOH with 0.1% TFA at 10 mL/min. The HPLC fractions of the desired purified compounds were first concentrated under reduced pressure using a rotary evaporator. The final concentrated aqueous solutions were lyophilized to dryness using Labconco Freezone 4.5 L (-84 $^oC$) lyophilizer. The peptides were analyzed for purity using Phenomenex Luna 5 µm C8(2) on the same RP-HPLC; gradient elution in $H_2O$/MeOH with 0.1% TFA at 1 mL/min. Peptide identities were confirmed via high resolution electrospray ionization mass spectrometry (HRMS, ESI/MS). Analyses were obtained on an Agilent 6545B Q-TOF LC/MS equipped with 1260 infinity II LC system with auto sampler. Samples were dissolved in $CH_3CN$ and eluted with a $CH_3CN/H_2O$ solution containing 0.1% formic acid

**Synthesis of TetD.** To prepare **TetD**, 2-Chlorotrityl chloride resin (0.142 mmol) was used such that the *C*-terminus would be a carboxylic acid upon cleavage. Synthesis was performed in accordance with established procedures for that resin. For the coupling of DBCO to the *N*-terminus of the peptide, 0.9 equiv. of DBCO-NHS was dissolved in 1 mL of dry DMF and added to the *N*-terminally deprotected peptide on resin resuspended in 2 mL of DMF. The coupling was carried out overnight at room temperature. Cleavage from the resin was then carried by the addition of 20% trifluoroacetic acid (TFA) in dichloromethane with subsequent shaking in room temperature for 1 h. The liquid phase was then filtered, concentrated and precipitated to yield crude peptide as described. The crude peptide was purified via HPLC as described in the general procedure. The **TetD** stock was characterized using UV-Vis absorbance at 309 nm ($\varepsilon$ = 12,000 $cm^{-1}M^{-1}$).

**Synthesis of the macrocyclization peptide library.** To prepare peptides in this library, Rink-amide resin (0.48 mmol/g of loading capacity) was used such that the *C*-terminus would be amidated upon cleavage. Synthesis was performed in accordance with established procedures for the resin. For all linear analogs within the library, the *N*-

terminus was acetylated by combining the peptide with a mixture of acetic anhydride, DIEA, and DMF (5:8.5:86.5, v/v) for 1 hour at room temperature. For the cyclization chemistry of the **Cyc0-4** and **Lar1-5** series, 10 equiv. of chloroacetic chloride and 15 equiv. of DIEA were dissolved in anhydrous DCM and coupled to the *N*-terminally deprotected peptide by shaking it for 2 hours at room temperature. Cleavage from resin was performed as described above. For the linear analogs, purification was then subsequently carried out as described above. For the macrocyclic peptides **Cyc0-4** and **Lar1-5**, the cleaved, crude peptide scaffolds were then dissolved in a mixture of $H_2O/CH_3CN$ (1:1, v/v) to an approximate concentration of ~1.5 mM. This was followed by the addition of 0.5 M $NH_4HCO_3$ (pH 8.5) to a final concentration of 50 mM to form the thioether bonds for macrocyclization. The cyclization reaction mixture was then stirred overnight at room temperature (~16 h). Cyclization was monitored using matrix-assisted laser desorption ionization time-of-flight (MALDI-TOF) mass spectroscopy (Shimadzu 8020). For the macrocyclization of peptide Dit1, the crude peptide scaffold was dissolved in a mixture of $H_2O/CH_3CN$ (1:1, v/v) to an approximate concentration of ~1.5 mM. This was followed by the addition of 0.5 M $NH_4HCO_3$ (pH 8.5) to a final concentration of 50 mM to form the disulfide bond required for macrocyclization. The cyclization reaction mixture was then stirred overnight at room temperature (~16 h). Cyclization was assessed by monitoring the presence of free thiols using Ellman's reagent as per the standard protocol provided by the manufacturer. For the macrocyclization **Dit2**, the crude peptide scaffold was dissolved in a mixture of $H_2O/CH_3CN$ (1:1, v/v) to an approximate concentration of ~1.5 mM. Further, the bis-electrophilic cyclization linker 1,4-Bis(bromomethyl)benzene was also dissolved into this mixture to an approximate concentration of ~1.5 mM (1 equiv.). Thioether bond formation between the **Dit2** peptide scaffold and the linker was brought about by the addition of 0.5 M $NH_4HCO_3$ (pH 8.5) to a final concentration of 50 mM. The cyclization reaction mixture was then stirred overnight at room temperature (~16 h). Cyclization was monitored using matrix-assisted laser desorption ionization time-of-flight (MALDI-TOF) mass spectroscopy (Shimadzu 8020). For all the above-mentioned macrocyclic peptides, upon confirmation of macrocyclization, the samples were then concentrated under reduced pressure using a rotary evaporator. The concentrated aqueous solutions were then lyophilized to dryness using a Labconco Freezone 4.5 L (-84 $^{o}$C) lyophilizer. Purification was subsequently carried out as described above. All peptides in this library were characterized using UV-Vis absorbance of the phenylalanine residue in their sequence at 257 nm ($\varepsilon$ = 195 $cm^{-1}M^{-1}$). For **Dit2**, the extinction coefficient at 257 nm of 1,4-Bis(bromomethyl)benzene was also determined separately using a calibration curve and accounted for during concentration determination of **Dit2**.

**Synthesis of the *N*-methylation and *N*-alkylation peptide libraries.** To prepare peptides in this library, Rink-amide resin (0.48 mmol/g of loading capacity) was used such that the *C*-terminus would be amidated upon cleavage. Synthesis was performed in accordance with established procedures for the resin. *N*-methyl amino acids were used for the *N*-methyl analogs in the peptide library. 2-azido-acetic acid was coupled on the *N*-terminus of each peptide on resin. For the peptoid library, the *N*-alkyl substituted glycine residues in the peptoids were added as per reported protocols. Briefly, a 25 mL peptide synthesis vessel containing 100 mg of Rink amide resin (0.048 mmol), conjugated with or without peptide residues, was added 20% piperidine in DMF (15 mL), with shaking at room temperature for 30 min. The resin was then washed with $CH_3OH$/DCM. After the last wash, to the resin was added 1 mL 0.6 M 2-bromo acetic acid with 200 μL of 50% DIC in DMF. The resin was shaken at room temperature for 20 min, then washed with DMF 5 times. 1 mL 1.5 M primary amine building blocks were then added to the resin and the resin was shaken at room temperature for 1-2 h. The rest of the amino acids are coupled in the same manner according to the sequence of the peptide. As before, 2-azido-acetic acid was coupled on the *N*-terminus of each peptide/peptoids on resin. Cleavage and purification were carried out as described before. All peptides in this library were characterized using UV-Vis absorbance of the phenylalanine residue in their sequence at 257 nm ($\varepsilon = 195\ cm^{-1}M^{-1}$).

**Synthesis of the tridecaptin series.** To prepare peptides in this library, 2-Chlorotrityl chloride resin (0.142 mmol) was used such that the *C*-terminus would remain as a carboxylic acid upon cleavage. Synthesis was performed in accordance with established procedures for the resin. Notably, n-Octanoic acid was coupled on the *N* terminus of the tridecaptin peptide derivatives on resin. n-Octanoic acid, (67.5 μL, 3 eq, 0.43 mmol) was dissolved in 5 mL DMF with Oxyma (3.0 eq, 60 mg, 0.43 mmol) and DIC (3.0 eq, 66 μL, 0.43 mmol), and added to the resin. For the macrocyclization **triCyc**, the crude peptide scaffold was dissolved in a mixture of $H_2O/CH_3CN$ (1:1, v/v) to an approximate concentration of ~1.5 mM. Further, the bis-electrophilic cyclization linker 1,4-Bis(bromomethyl)benzene was also dissolved into this mixture to an approximate concentration of ~1.5 mM (1 equiv.). Thioether bond formation between the **triCyc** peptide scaffold and the linker was brought about by the addition of 0.5 M $NH_4HCO_3$ (pH 8.5) to a final concentration of 50 mM. The cyclization reaction mixture was then stirred overnight at room temperature (~16 h). Cyclization was monitored using matrix-assisted laser desorption ionization time-of-flight (MALDI-TOF) mass spectroscopy (Shimadzu 8020). Upon confirmation of macrocyclization, the sample was then concentrated under reduced pressure using a rotary evaporator. The concentrated aqueous solution was then lyophilized to dryness using a Labconco Freezone 4.5 L (-84 $^{o}$C) lyophilizer. Purification was subsequently carried out as described above.

**Synthesis of the griselimycin series.** To prepare the linear peptide derivatives in this library, Sieber-amide resin (0.122 mmol) was used such that the *C*-terminus would be amidated upon cleavage. Synthesis was performed in accordance with established procedures for the resin. *N*-methyl amino acids were used accordingly for the *N*-methyl residues on the peptide. 2-azido-acetic acid was coupled on the *N*-terminus of each peptide on resin. Peptides were cleaved from the resin using 20% trifluoroacetic acid (TFA) in dichloromethane with 1% TIPS, with shaking at room temperature for 1 h. Cyclic griselimycin derivatives were synthesized on 2-Chlorotrityl chloride resin (0.142 mmol) such that the *C*-terminus would remain as a carboxylic acid upon cleavage. Synthesis was begun with Fmoc-*N*-methyl-D-Leu/Fmoc-D-Leu and further procedures were followed based on established protocols,[8] yielding linear peptides bound to resin with an unprotected Thr side chain. *N*-methyl amino acids were used accordingly for the *N*-methyl residues on the peptide. 2-azido-acetic acid was coupled to the *N*-terminus of each peptide on resin. Esterification was then carried out on resin by adding Fmoc-glycine (10 eq) dissolved in anhydrous DMF/DCM (1:9, roughly 0.2 g/mL) with DMAP (0.25 eq) and DIC (10 eq). The solution was first shaken rapidly until the formation of solid precipitate was observed, before adding to the resin. The mixture was then shaken at room temperature for 3 h and drained. Another portion of the solution with the same amount of the reactants (10 eq Fmoc-glycine, 0.25 eq DMAP, 10 eq DIC) was immediately added without washing the resin, followed by shaking overnight at room temperature. The resin was drained and washed with DMF until the thick precipitate formed during the reaction was removed, then washed with $CH_3OH$/DCM (3x). The Fmoc group on glycine residues was then deprotected with 20% piperidine in DMF, followed by cleavage of the branched peptide from the resin using 25% hexafluoro isopropanol (HFIP) in DCM (1 h shaking at room temperature). Solvents were then filtered and evaporated under reduced pressure using a rotary evaporator. DCM was added and evaporated 2-3 times until uniform crystals were formed. The sample was stored in a vacuum desiccator overnight prior to cyclization. The cyclization reaction was carried out in HPLC grade $CH_3CN$ with a final concentration of approximately 1 mM of peptide. Briefly, COMU (3 eq) was placed in a round-bottom flask with a stir bar, followed by $CH_3CN$ and DIEA (3eq). In a separate vessel, the branched linear peptide was dissolved in $CH_3CN$ (10-15 mL) and DIEA (3 eq). The peptide was added dropwise using a pressure equalizing addition funnel in portions to the round-bottom flask with rapid stirring during a period of 30 minutes. Rapid stirring was continued at room temperature for 16 h. Cyclization was monitored using matrix-assisted laser desorption ionization time-of-flight (MALDI-TOF) mass spectroscopy (Shimadzu 8020) until no uncyclized precursor can be detected. The solution was then concentrated under reduced pressure using a rotary evaporator for downstream purification. Purification was subsequently carried out as described above.

**Characterization of test molecules.** The peptides were analyzed for purity using Phenomenex Luna 5 µm C8(2) on the same RP-HPLC; gradient elution in $H_2O$/MeOH with 0.1% TFA at 1 mL/min. Peptide identities were confirmed via high resolution electrospray ionization mass spectrometry (HRMS, ESI/MS). Analyses were obtained on an Agilent 6545B Q-TOF LC/MS equipped with 1260 infinity II LC system with auto sampler. Samples were dissolved in $CH_3CN$ and eluted with a $CH_3CN/H_2O$ solution containing 0.1% formic acid. Characterization data are provided below.

**Lin0**

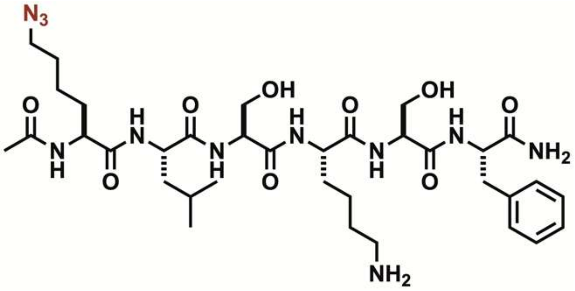

Analytical HPLC Chromatogram of **Lin0**.

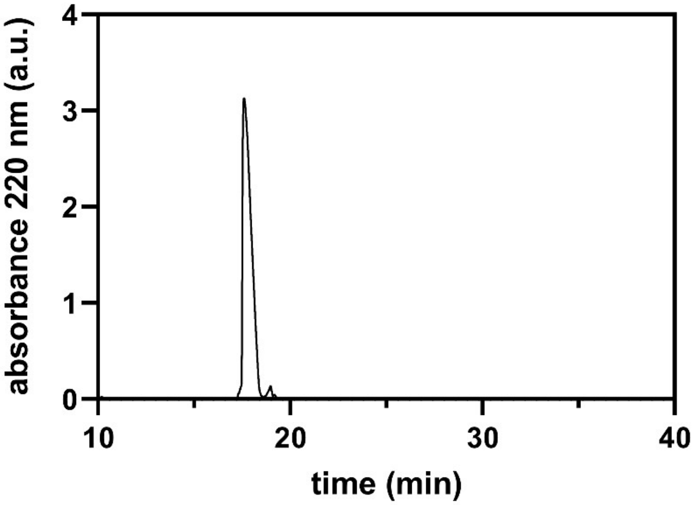


QTOF High Resolution Mass Spectrum for **Lin0** (m/z 776.4341 for $[M+H]^+$ found 776.4506)

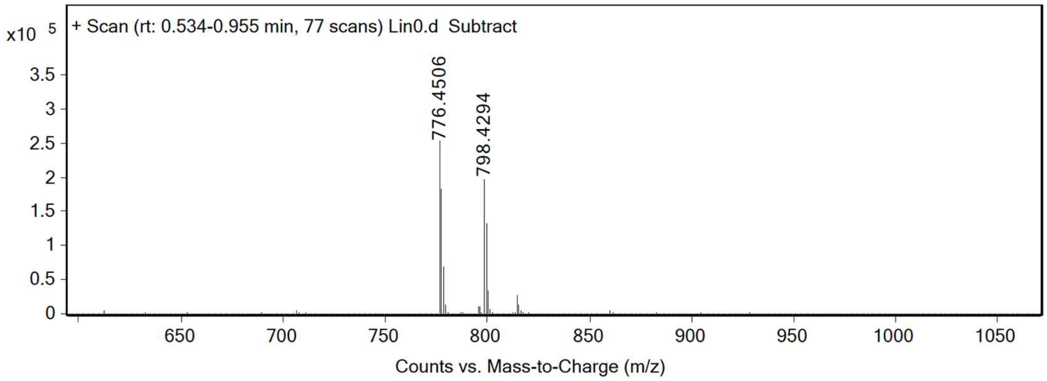

**Lin1**

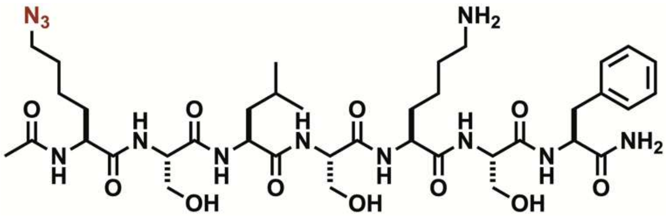


Analytical HPLC Chromatogram of **Lin1**.

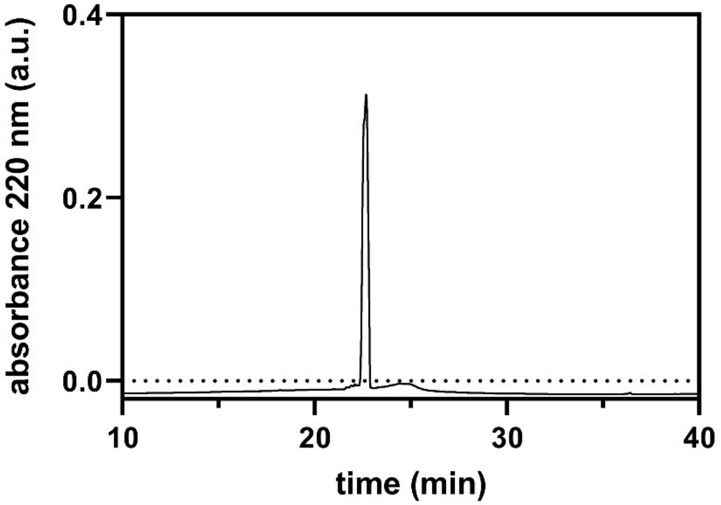


QTOF High Resolution Mass Spectrum for **Lin1** (m/z 863.4661 for $[M+H]^+$ found 863.4808)

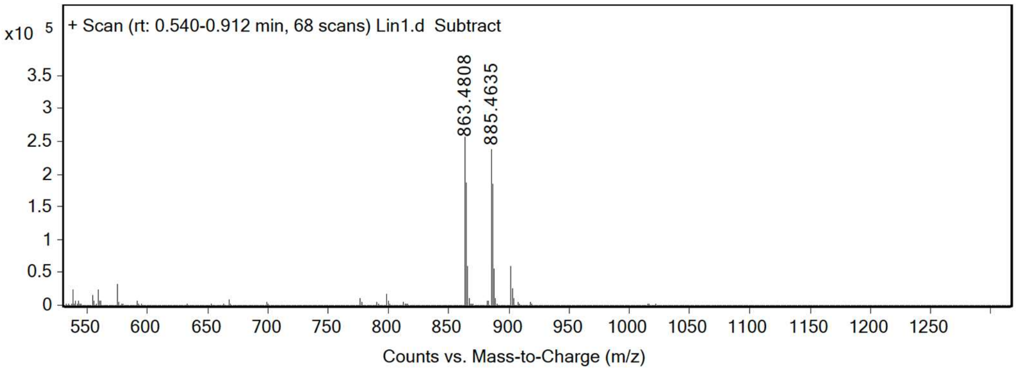

**Lin2**

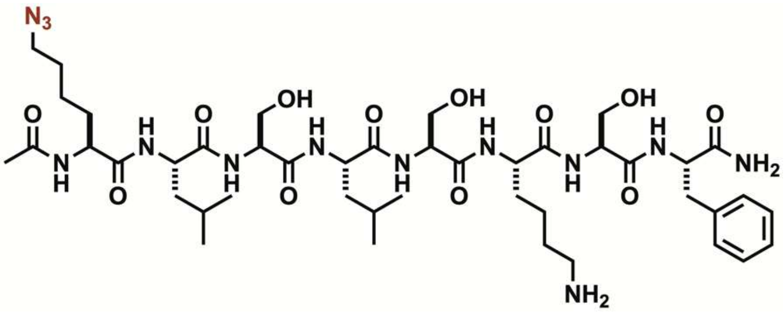


Analytical HPLC Chromatogram of **Lin2**.

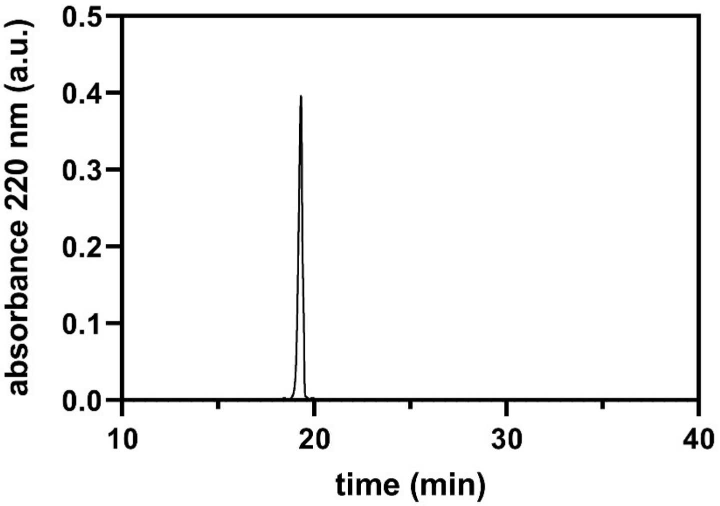


QTOF High Resolution Mass Spectrum for **Lin2** (m/z 976.5502 for $[M+H]^+$ found 976.5724)

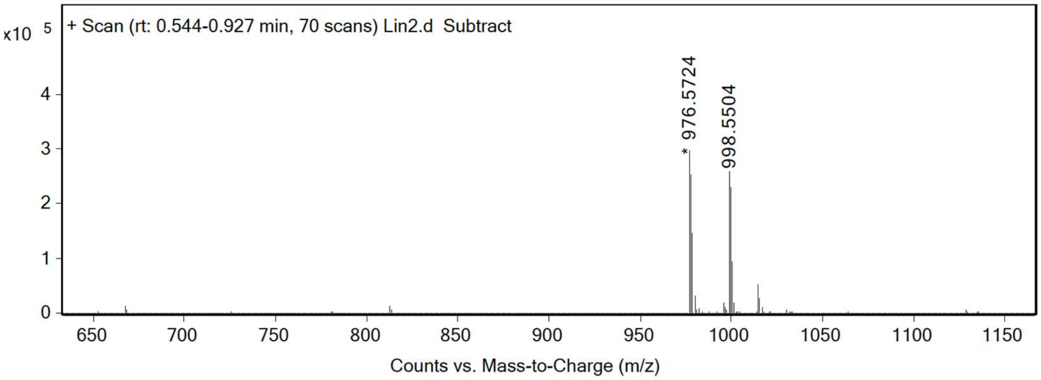

**Lin3**

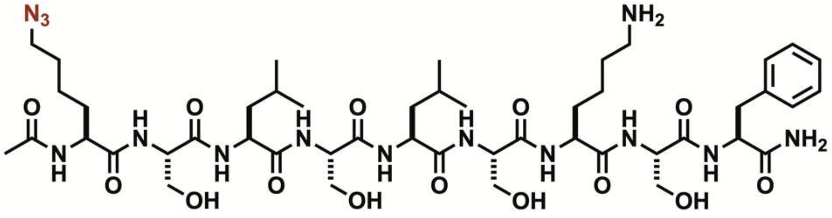


Analytical HPLC Chromatogram of **Lin3**.

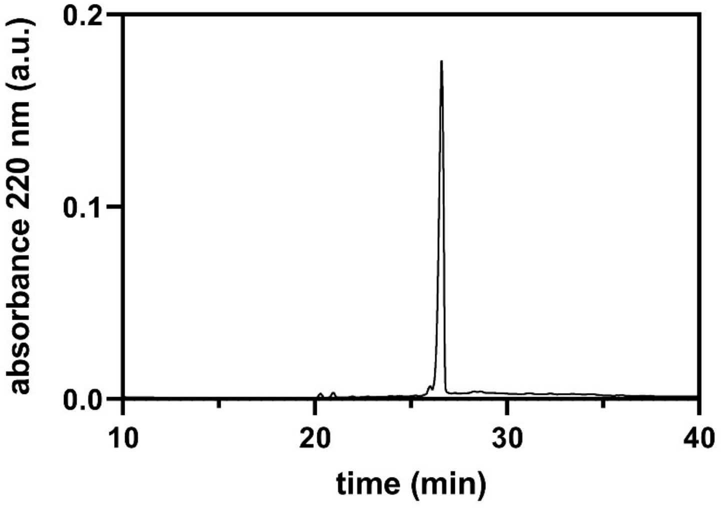


QTOF High Resolution Mass Spectrum for **Lin3** (m/z 1063.5822 for $[M+H]^+$ found 1063.5973)

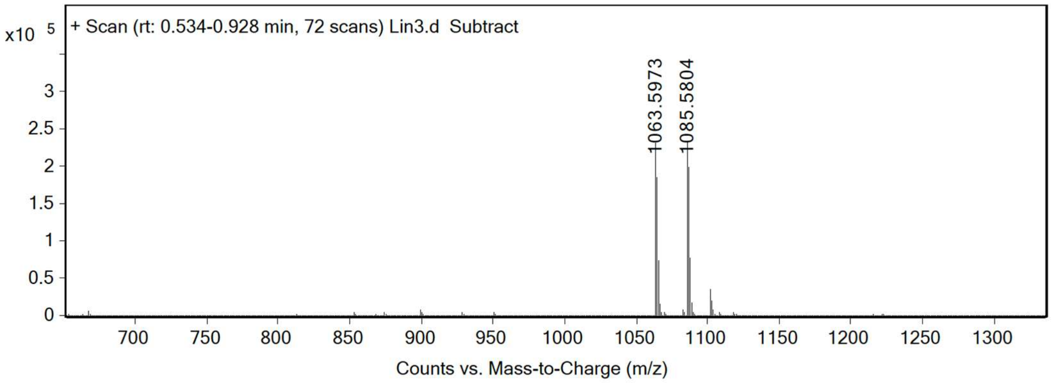

**Cyc0**

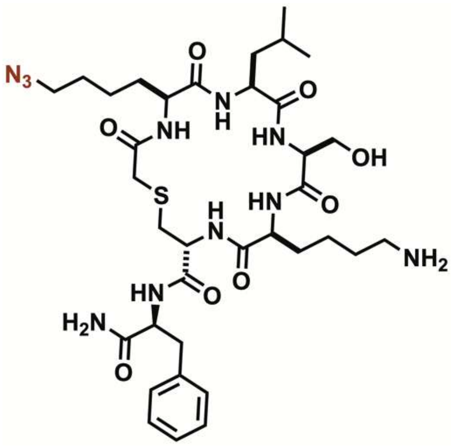


Analytical HPLC Chromatogram of **Cyc0**.

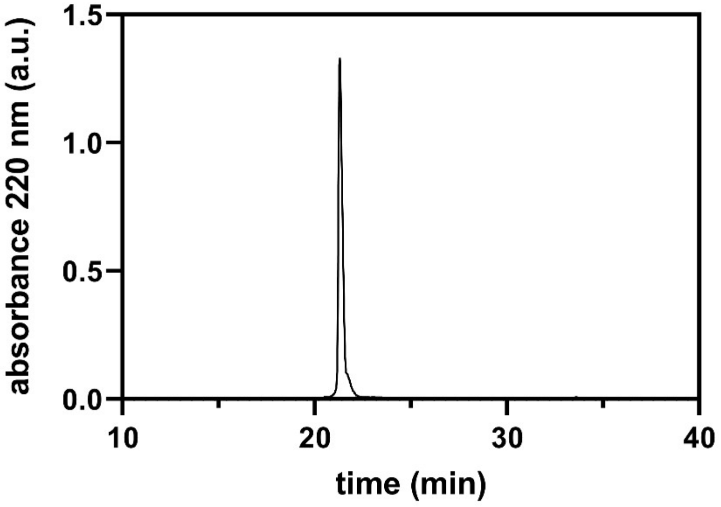


QTOF High Resolution Mass Spectrum for **Cyc0** (m/z 790.3956 for $[M+H]^+$ found 790.4043)

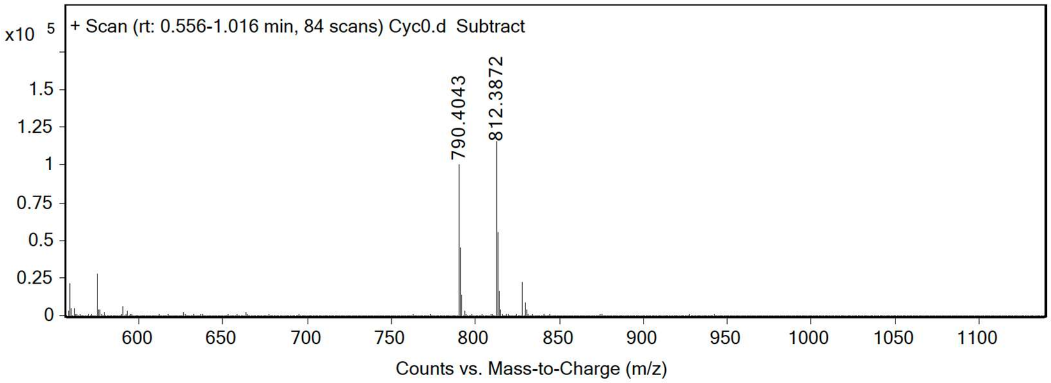

**Cyc1**

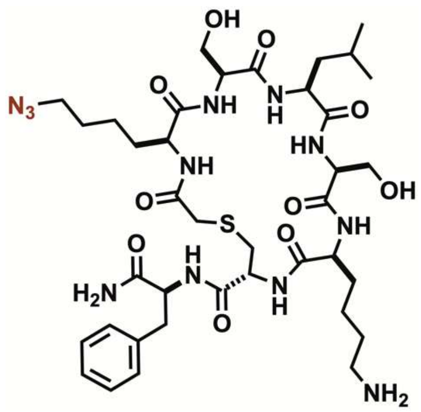

Analytical HPLC Chromatogram of **Cyc1**.

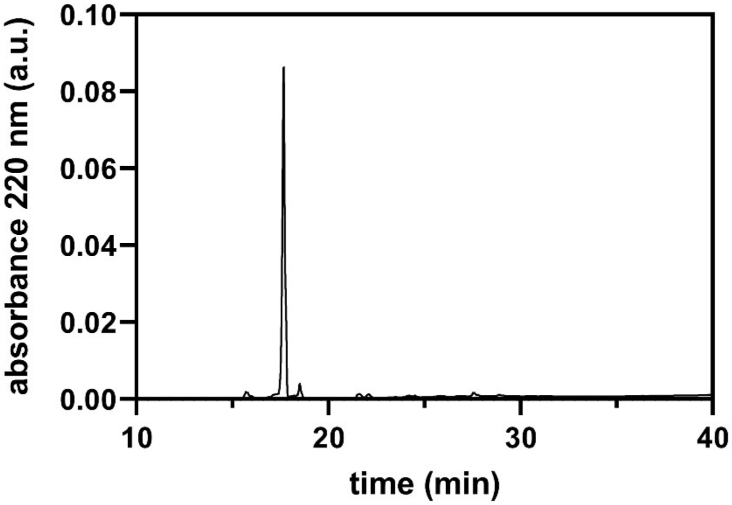


QTOF High Resolution Mass Spectrum for **Cyc1** (m/z 877.4276 for $[M+H]^+$ found 877.4363)

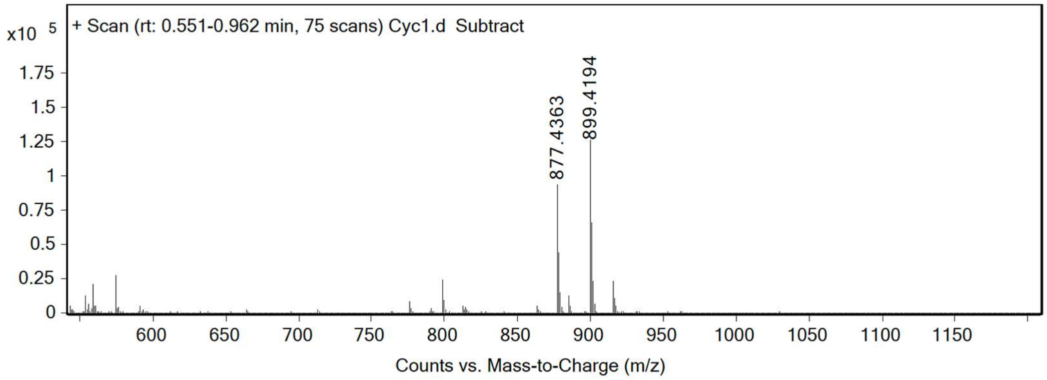

**Cyc2**

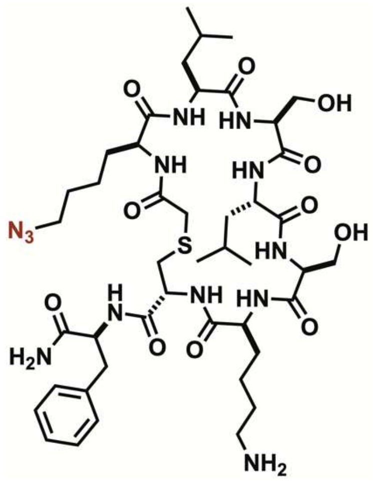


Analytical HPLC Chromatogram of **Cyc2**.

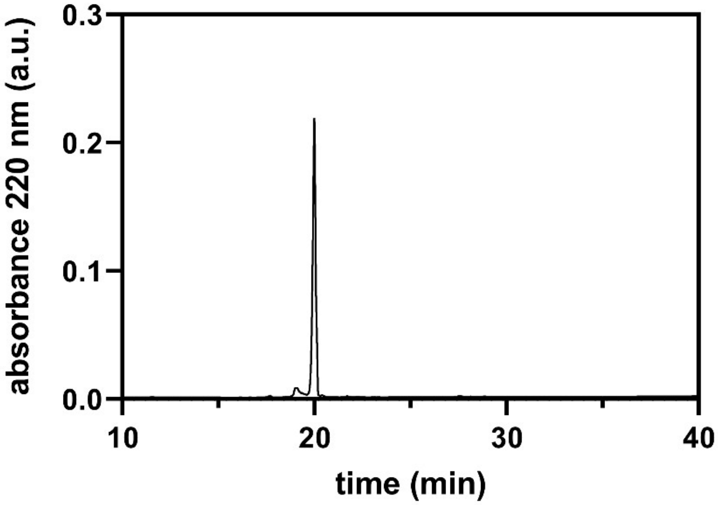


QTOF High Resolution Mass Spectrum for **Cyc2** (m/z 990.5117 for $[M+H]^+$ found 990.5213)

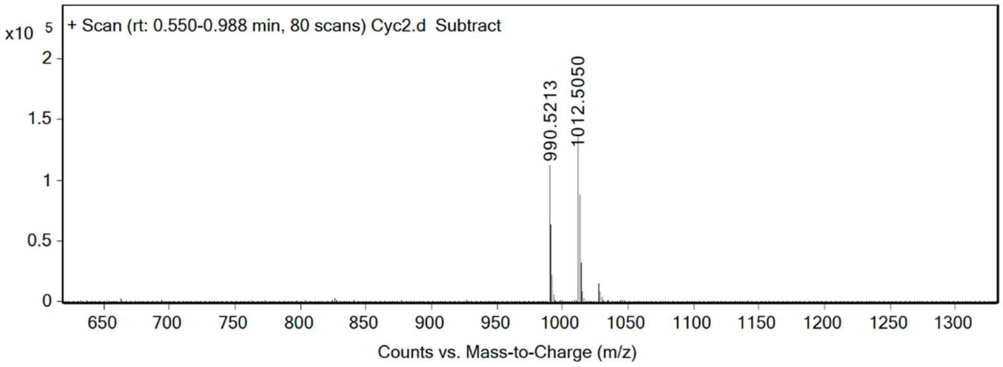

**Cyc3**

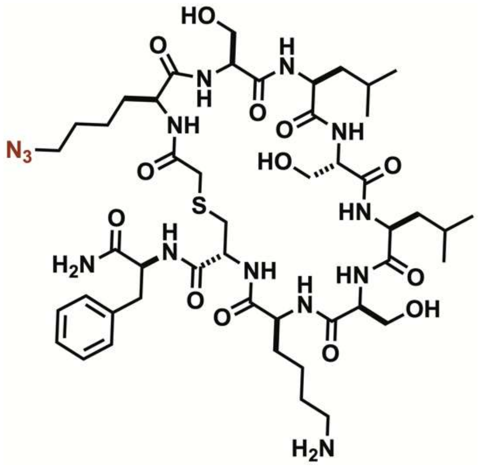


Analytical HPLC Chromatogram of **Cyc3**.

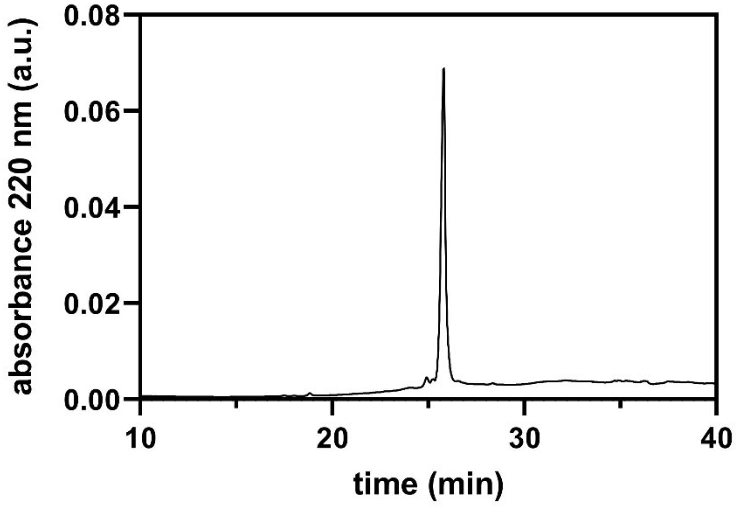


QTOF High Resolution Mass Spectrum for **Cyc3** (m/z 1077.5437 for $[M+H]^+$ found 1077.5532)

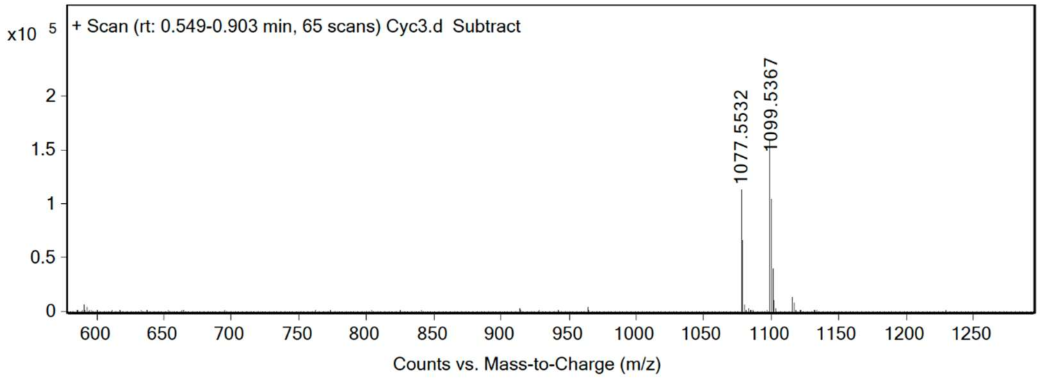

**Lar0**

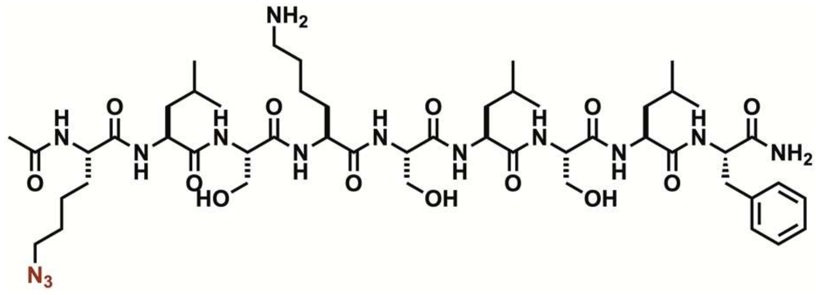


Analytical HPLC Chromatogram of **Lar0**.

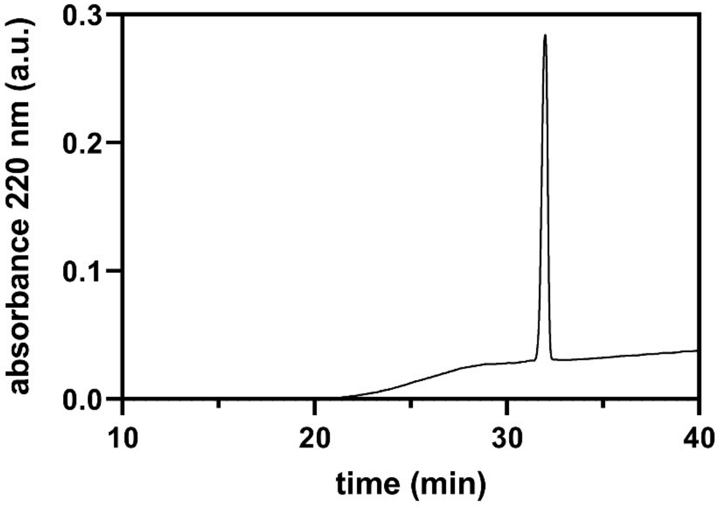


QTOF High Resolution Mass Spectrum for **Lar0** (m/z 1089.6342 for $[M+H]^+$ found 1089.6507)

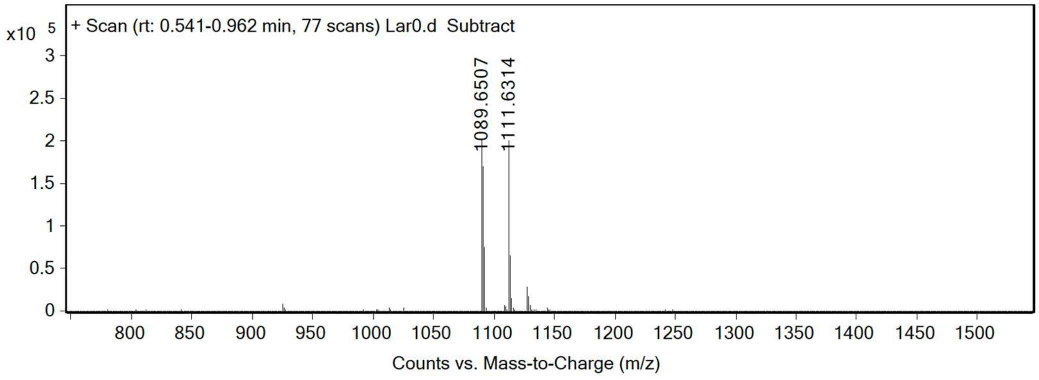

**Lar1**

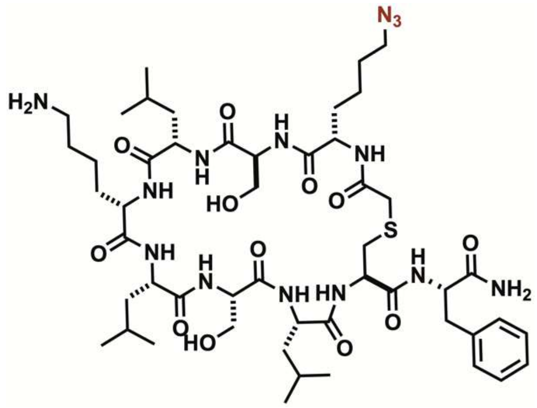


Analytical HPLC Chromatogram of **Lar1**.

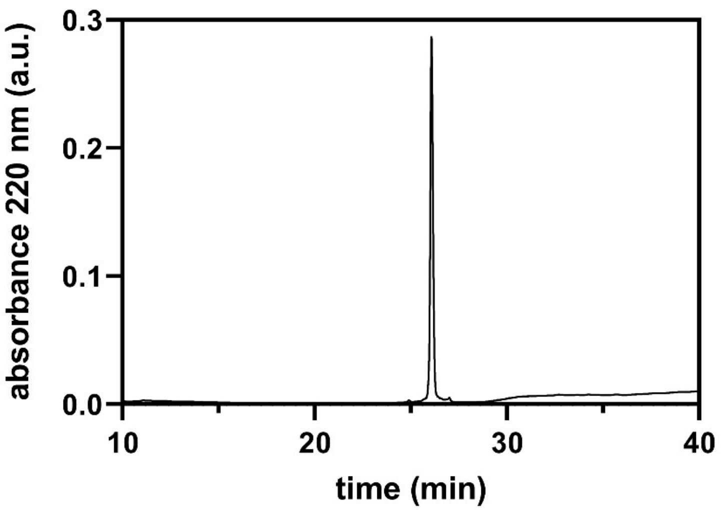


QTOF High Resolution Mass Spectrum for **Lar1** (m/z 1089.6342 for $[M+H]^+$ found 1103.6047)

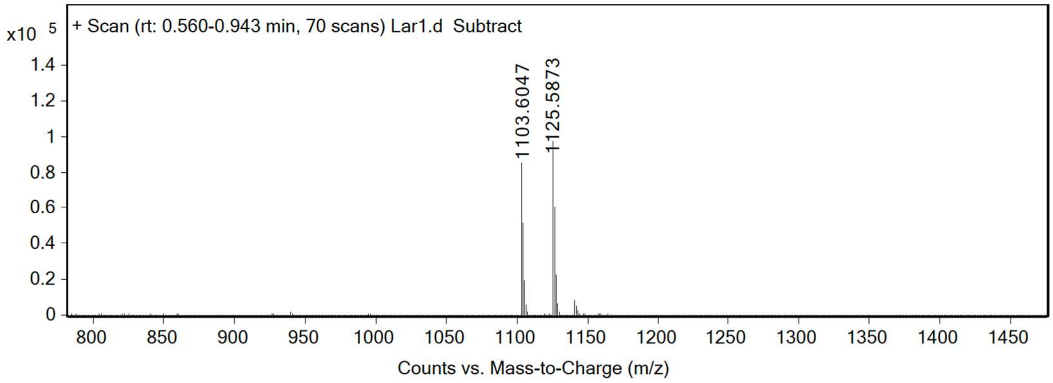

**Lar2**

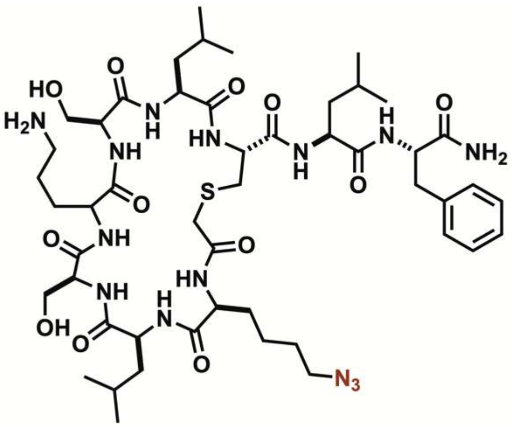


Analytical HPLC Chromatogram of **Lar2**.

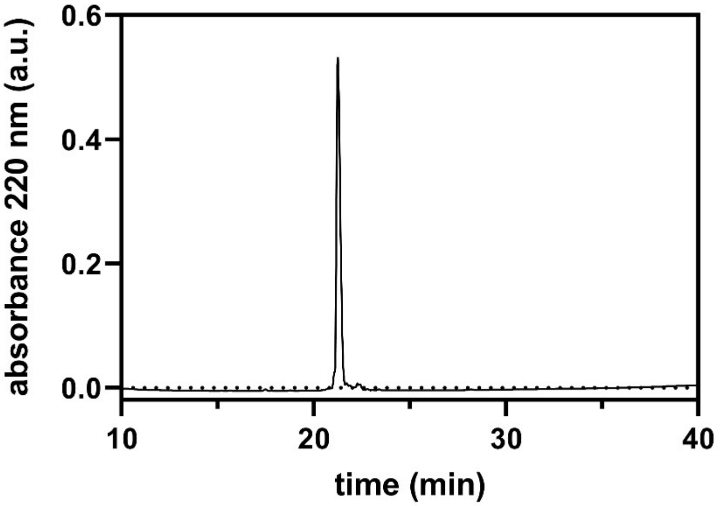


QTOF High Resolution Mass Spectrum for **Lar2** (m/z 1103.5957 for $[M+H]^+$ found 1103.6056)

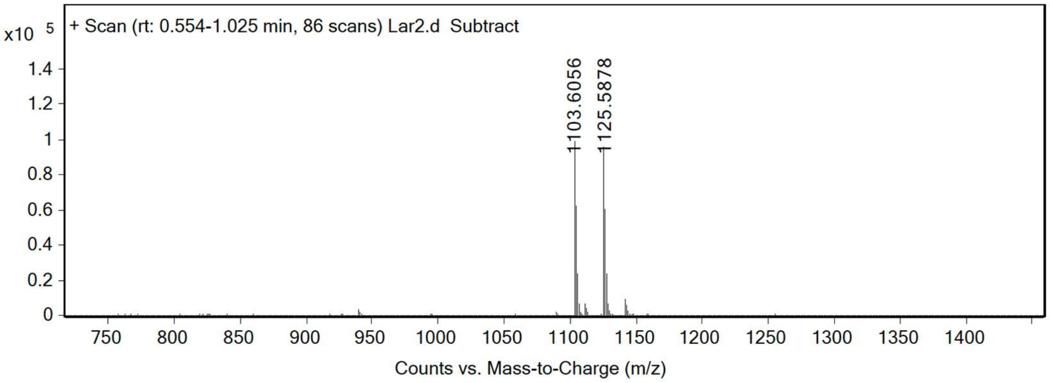

**Lar3**

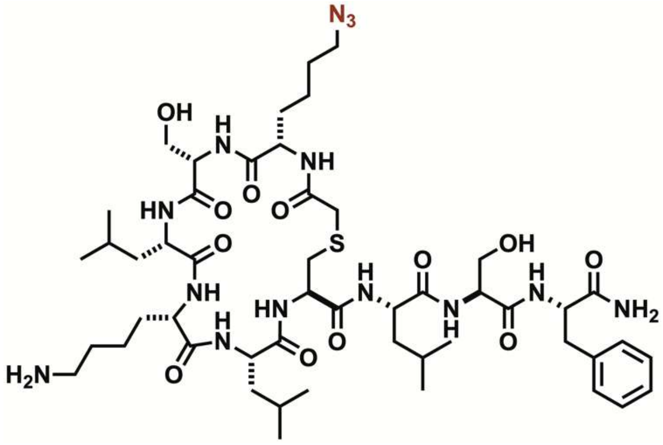


Analytical HPLC Chromatogram of **Lar3**.

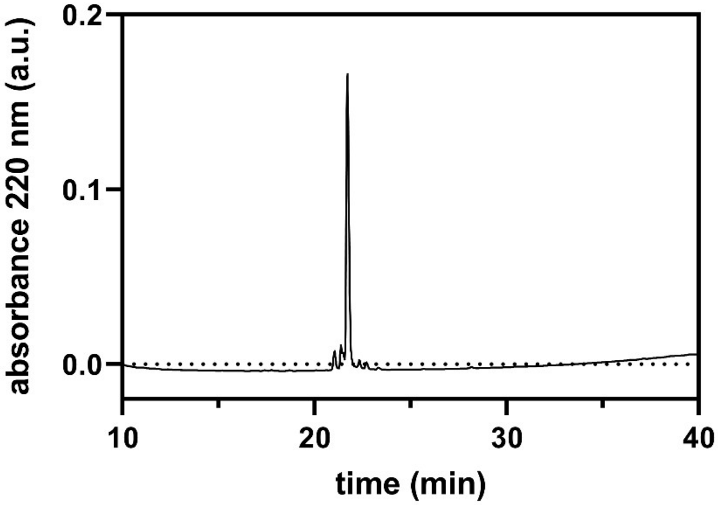


QTOF High Resolution Mass Spectrum for **Lar3** (m/z 1103.5957 for $[M+H]^+$ found 1103.6056)

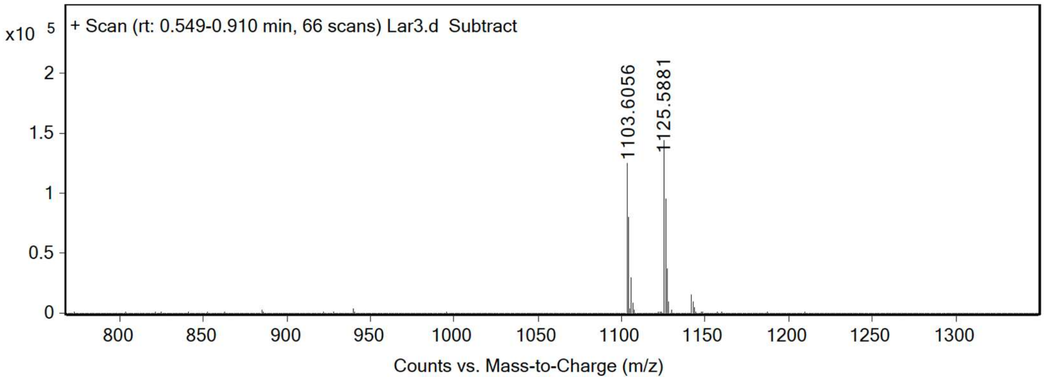

**Lar4**

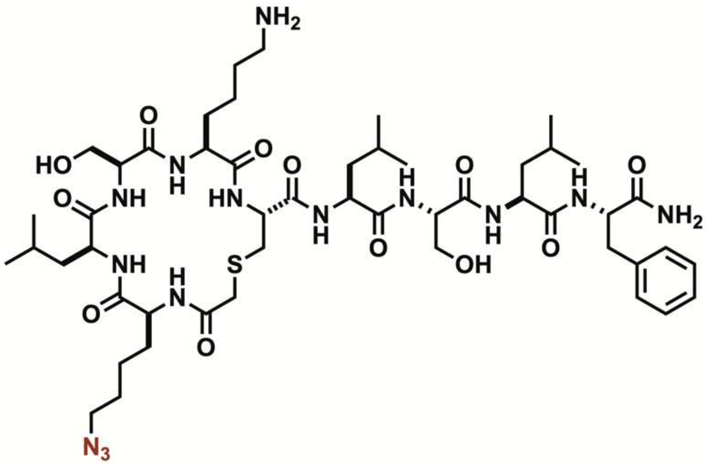


Analytical HPLC Chromatogram of **Lar4**.

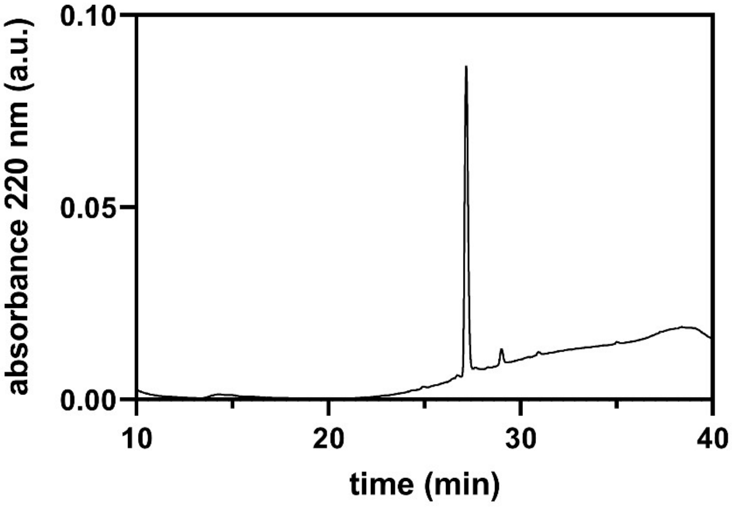


QTOF High Resolution Mass Spectrum for **Lar4** (m/z 1103.5957 for $[M+H]^+$ found 1103.6057)

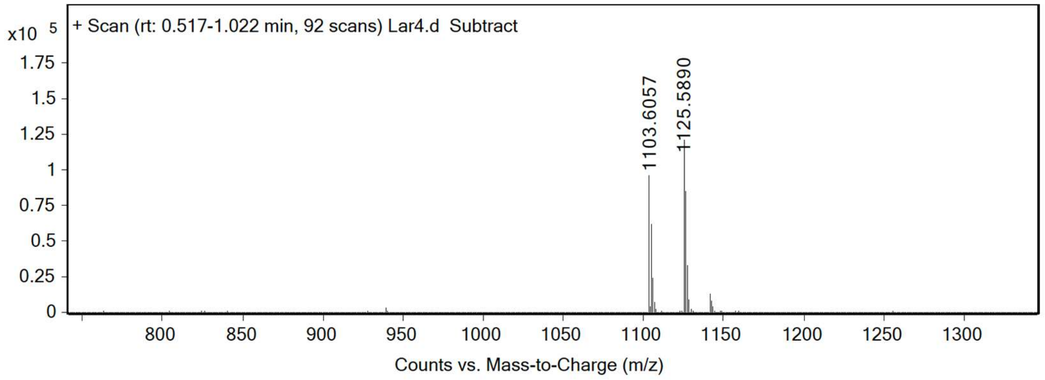

**Lar5**

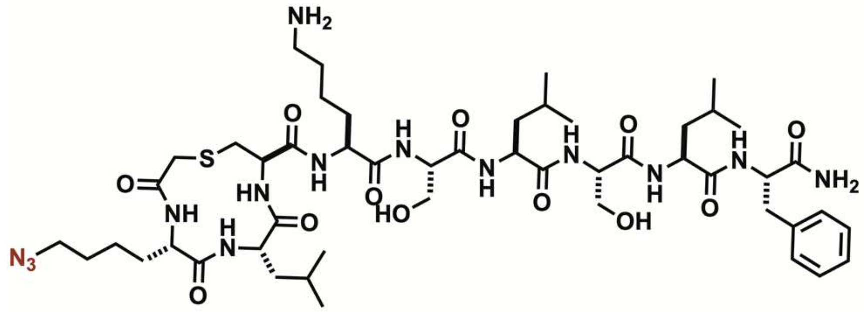


Analytical HPLC Chromatogram of **Lar5**.

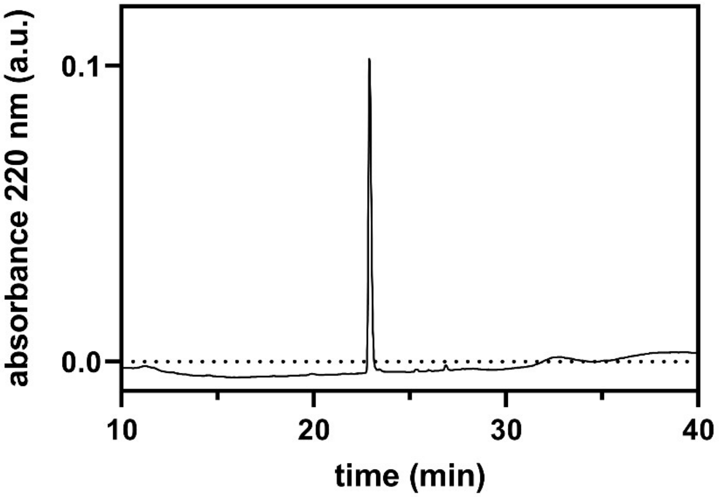


QTOF High Resolution Mass Spectrum for **Lar5** (m/z 1103.5957 for $[M+H]^+$ found 1103.6039)

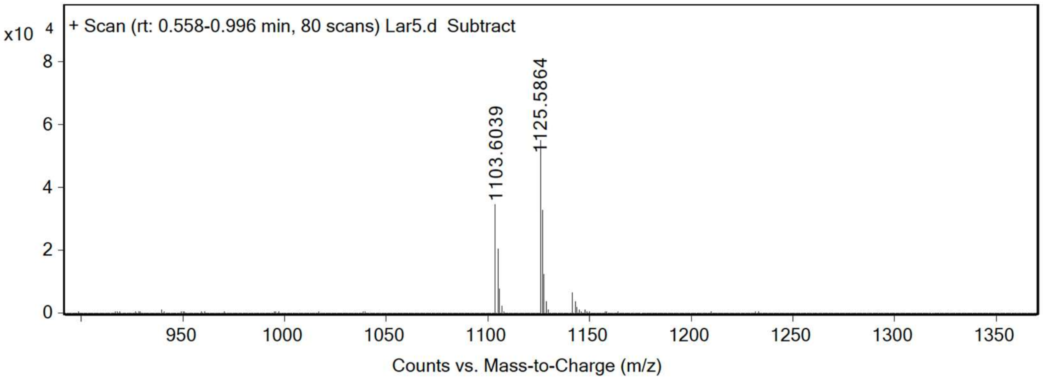

**Dit0**

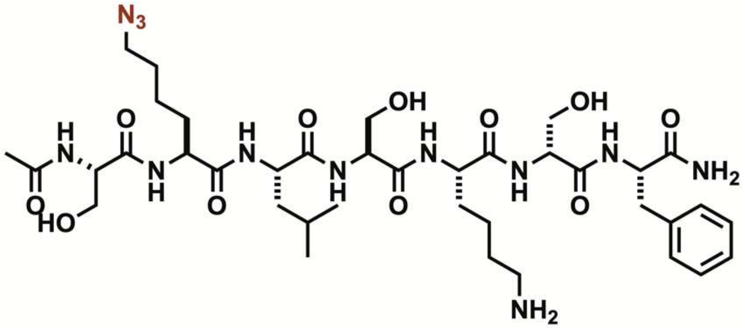


Analytical HPLC Chromatogram of **Dit0**.

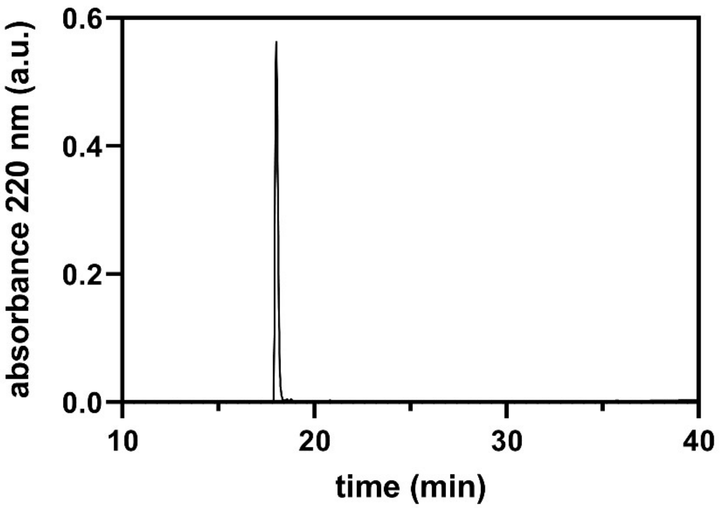


QTOF High Resolution Mass Spectrum for **Dit0** (m/z 863.4661 for $[M+H]^+$ found 863.4819)

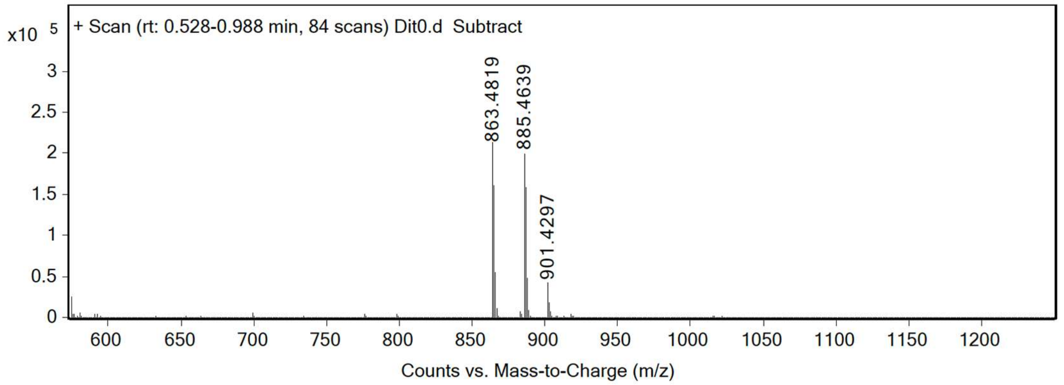

**Dit1**

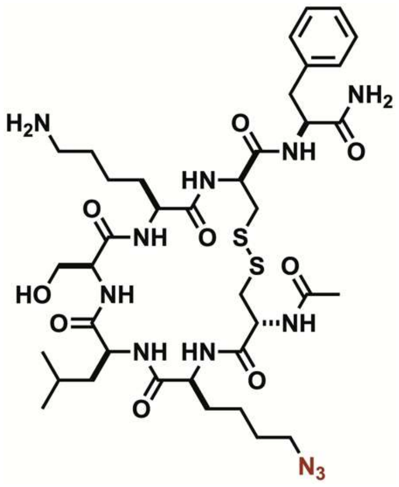


Analytical HPLC Chromatogram of **Dit1**.

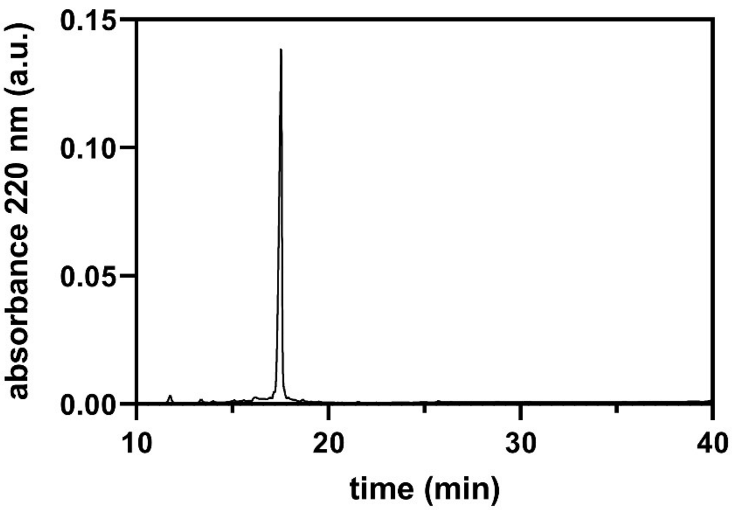


QTOF High Resolution Mass Spectrum for **Dit1** (m/z 893.4048 for $[M+H]^+$ found 893.4134)

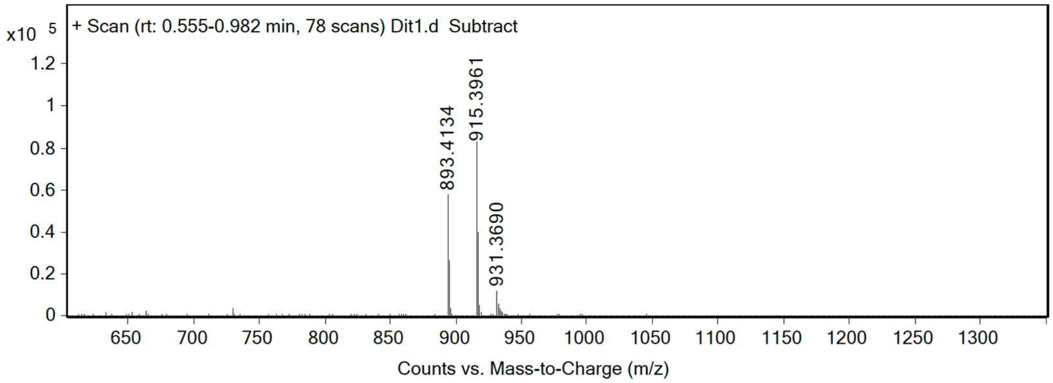

**Dit2**

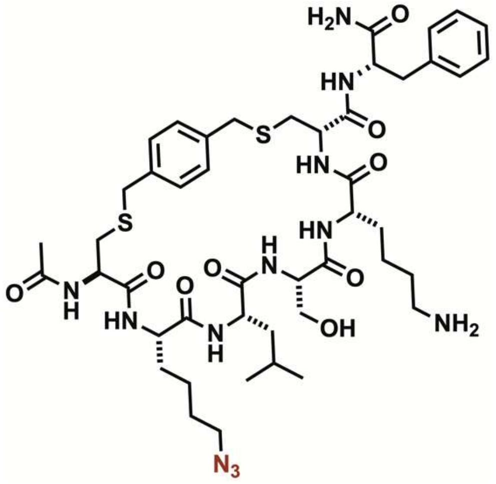


Analytical HPLC Chromatogram of **Dit2**.

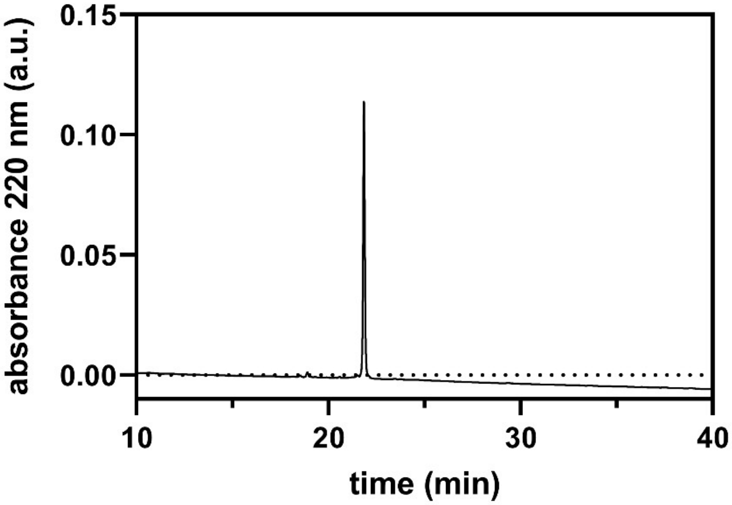


QTOF High Resolution Mass Spectrum for **Dit2** (m/z 997.4674 for $[M+H]^+$ found 997.4771)

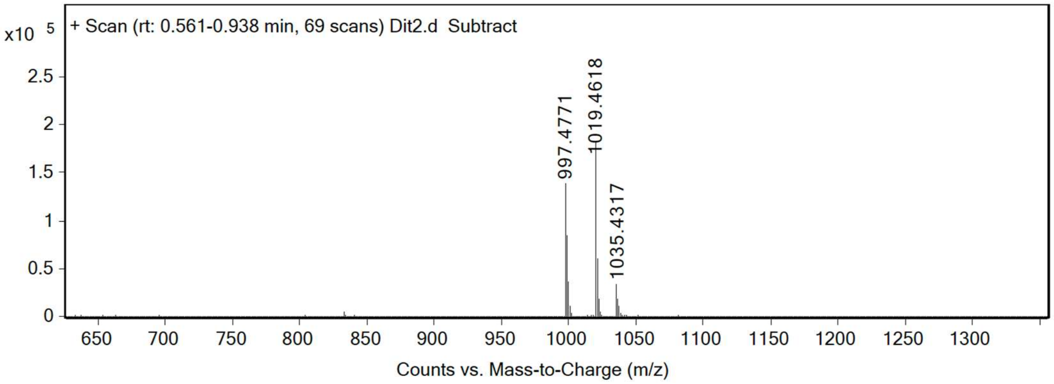

**Nmet0**

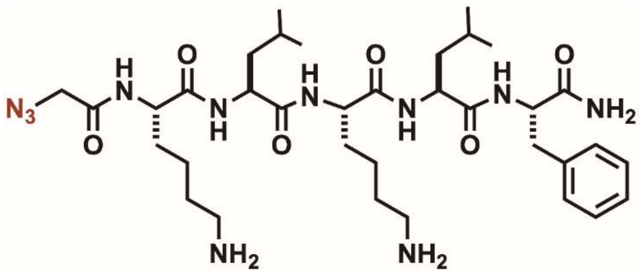


Analytical HPLC Chromatogram of **Nmet0**.

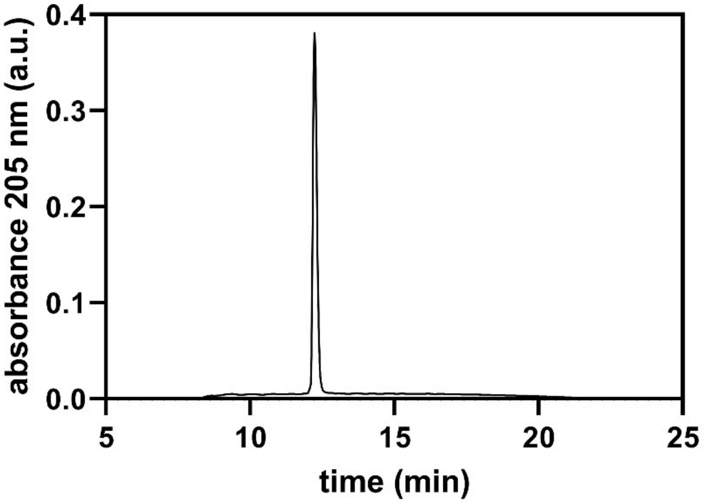


QTOF High Resolution Mass Spectrum for **Nmet0** (m/z 730.4723 for $[M+H]^+$ found 730.4719)

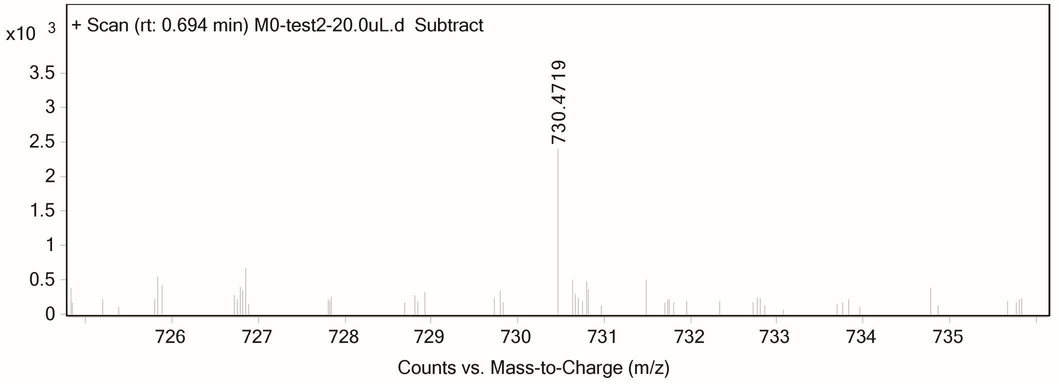

**Nmet1**

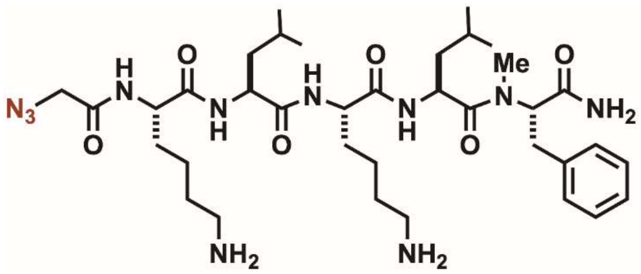


Analytical HPLC Chromatogram of **Nmet1**

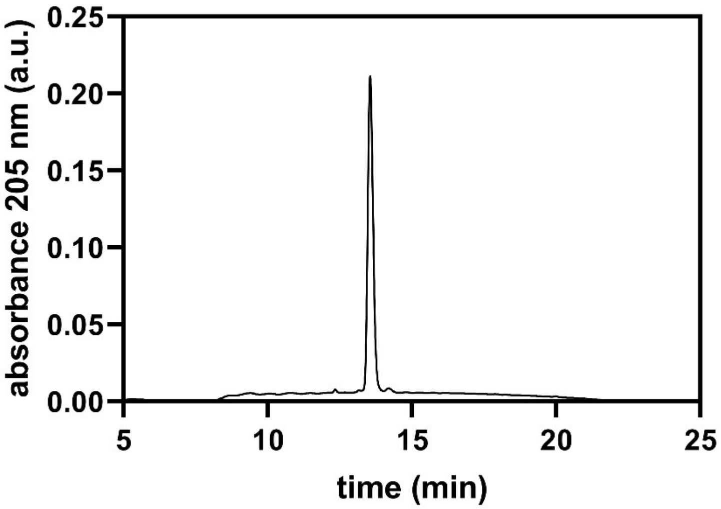


QTOF High Resolution Mass Spectrum for **Nmet1** (m/z 744.4879 for $[M+H]^+$ found 744.4878)

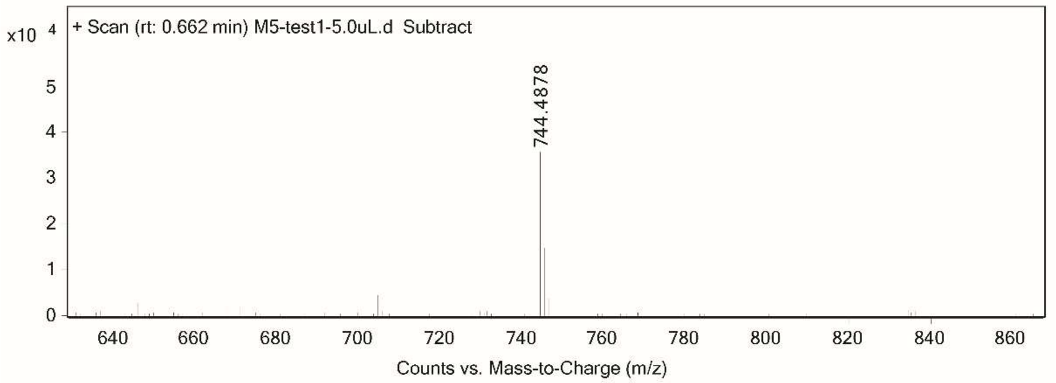

**Nmet2**

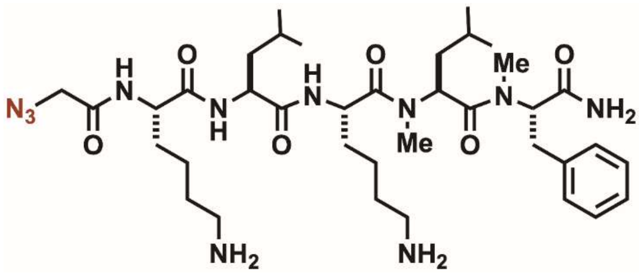


Analytical HPLC Chromatogram of **Nmet2**.

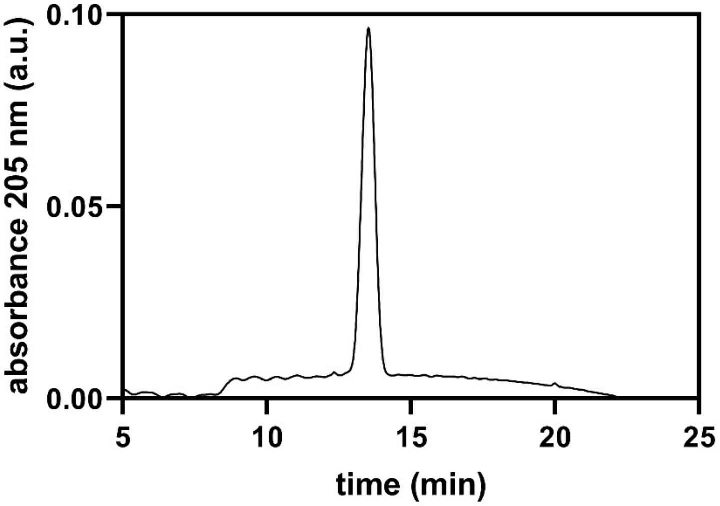


QTOF High Resolution Mass Spectrum for **Nmet2** (m/z 758.5036 for $[M+H]^+$ found 758.5039)

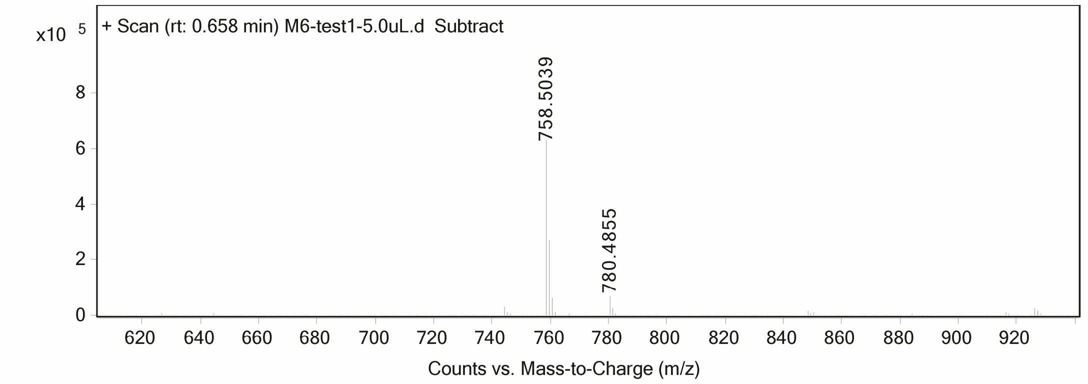

**Nmet3**

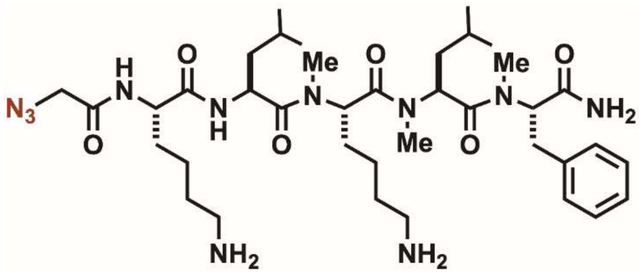


Analytical HPLC Chromatogram of **Nmet3**.

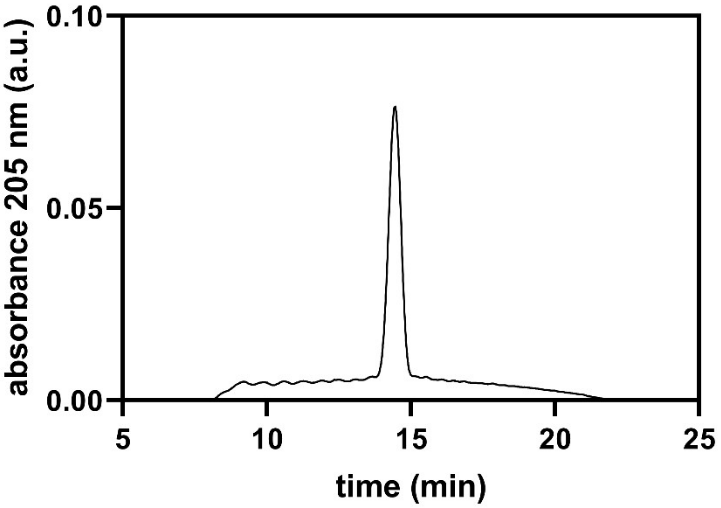


QTOF High Resolution Mass Spectrum for **Nmet3** (m/z 772.5192 for $[M+H]^+$ found 772.5191)

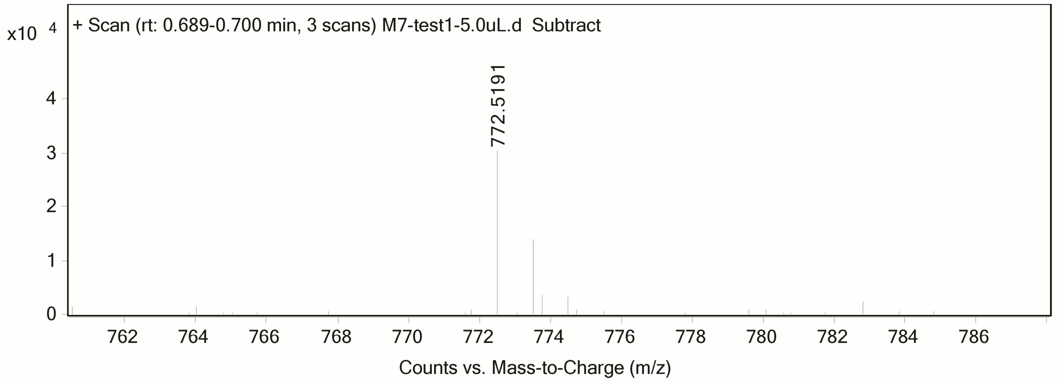

**Nmet4**

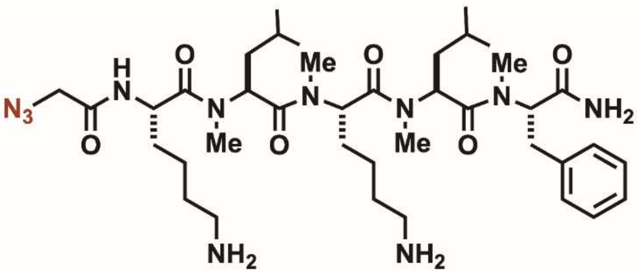


Analytical HPLC Chromatogram of **Nmet4**.

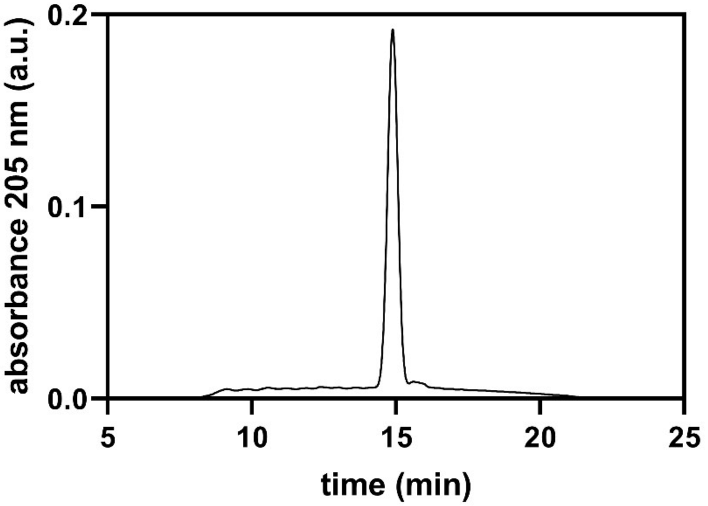


QTOF High Resolution Mass Spectrum for **Nmet4** (m/z 786.5349 for $[M+H]^+$ found 786.5351)

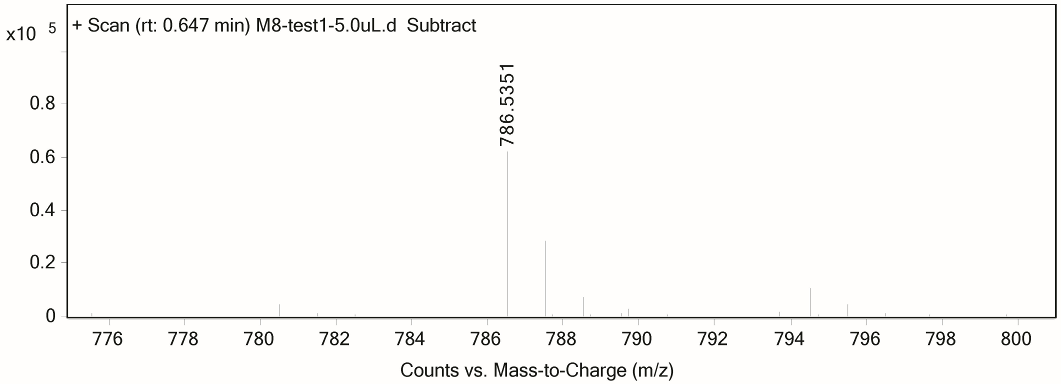

**Nmet5**

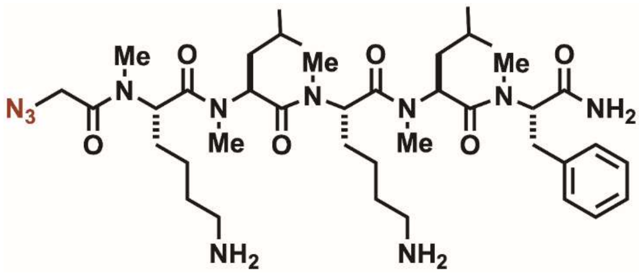

Analytical HPLC Chromatogram of **Nmet5**.

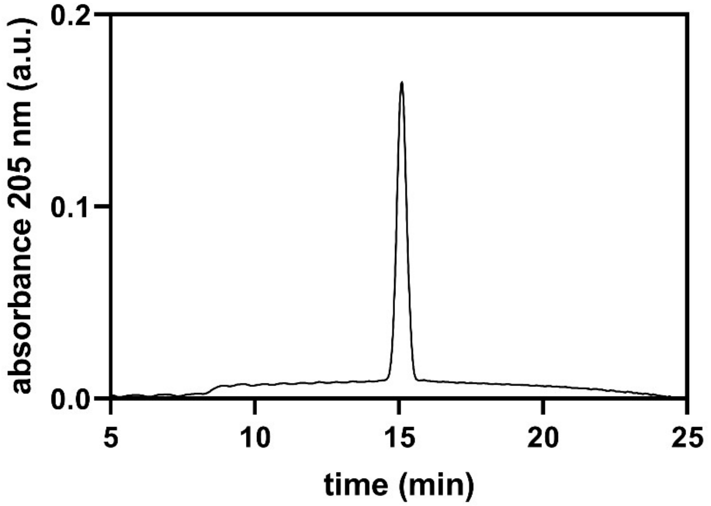


QTOF High Resolution Mass Spectrum for **Nmet5** (m/z 800.5505 for $[M+H]^+$ found 800.5506)

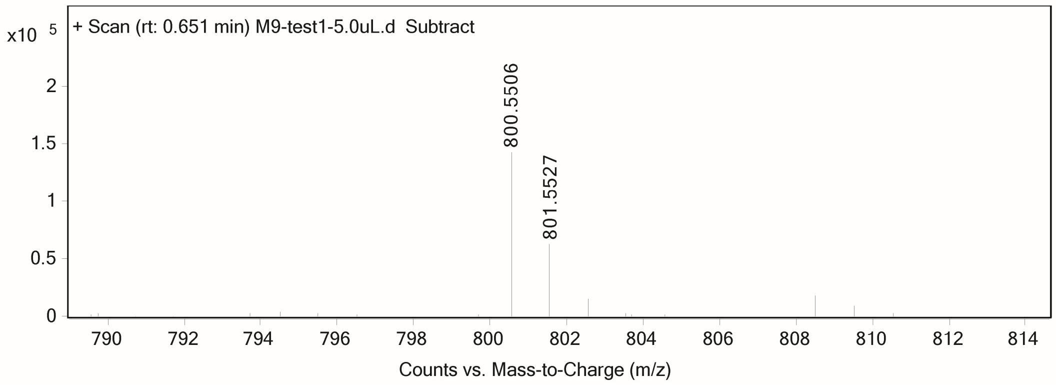

**Nmet6**

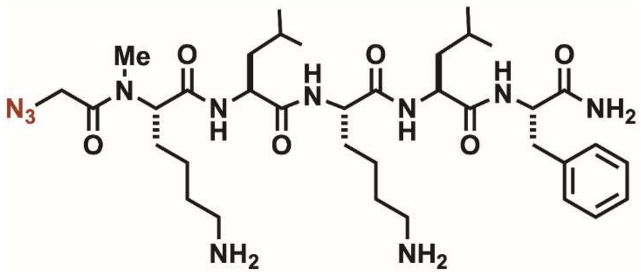


Analytical HPLC Chromatogram of **Nmet6**.

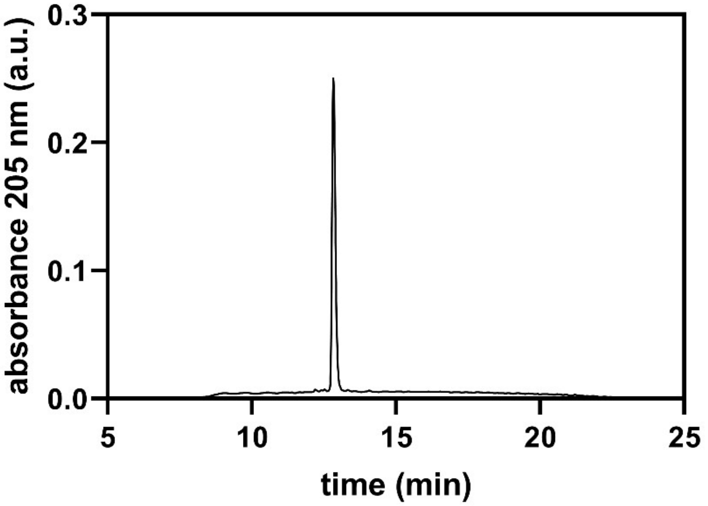


QTOF High Resolution Mass Spectrum for **Nmet6** (m/z 744.4879 for $[M+H]^+$ found 744.4916)

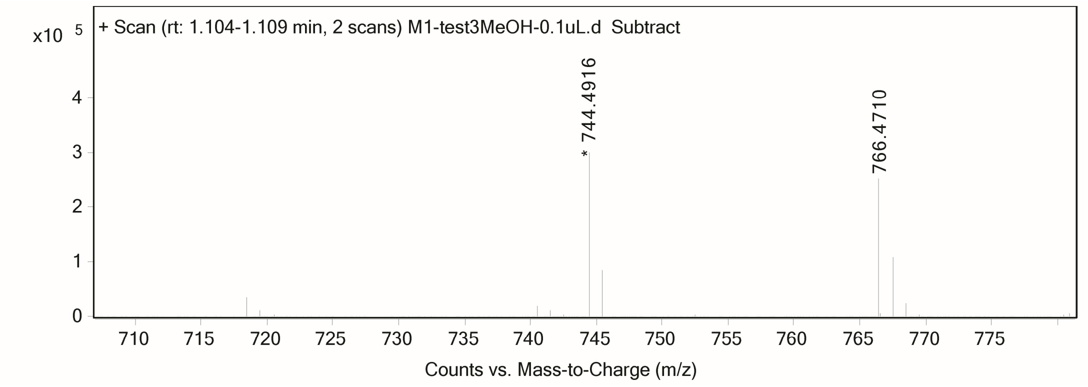

**Nmet7**

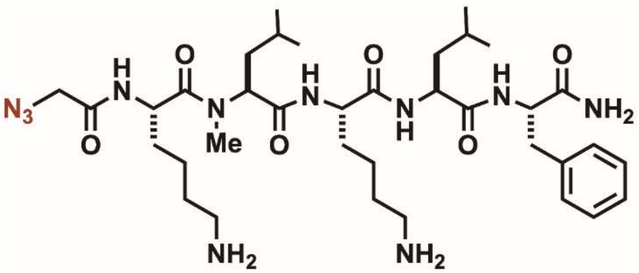


Analytical HPLC Chromatogram of **Nmet7**.

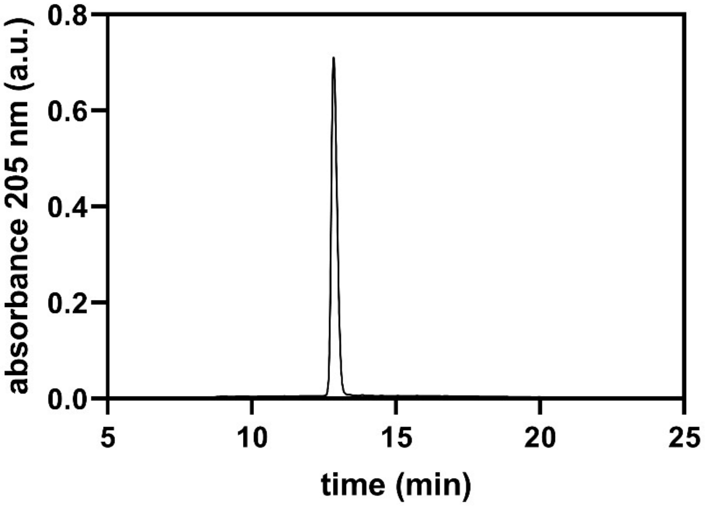


QTOF High Resolution Mass Spectrum for **Nmet7** (m/z 744.4879 for $[M+H]^+$ found 744.4873)

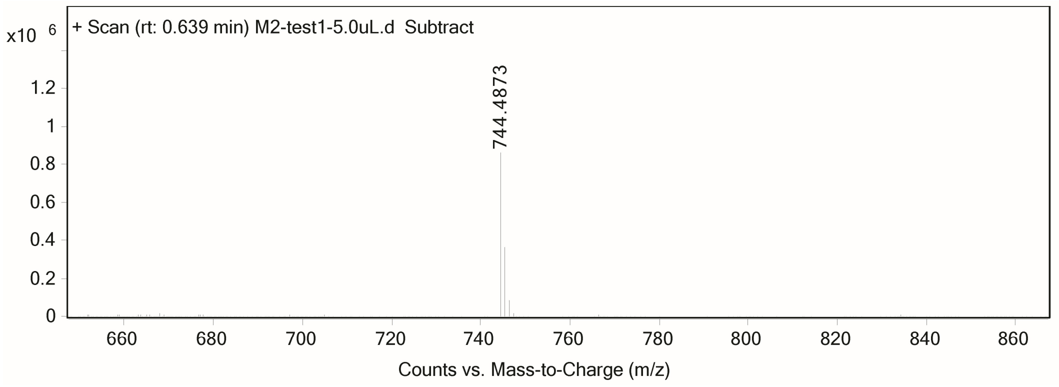

**Nmet8**

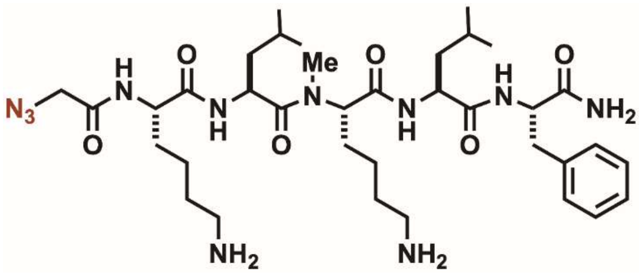


Analytical HPLC Chromatogram of **Nmet8**.

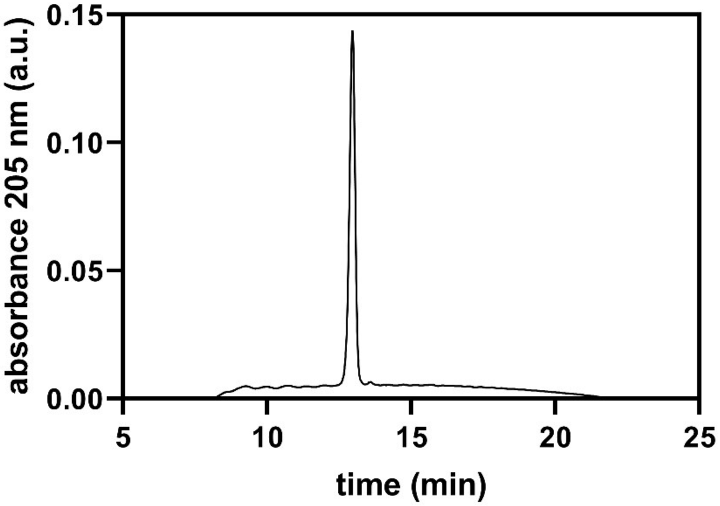


QTOF High Resolution Mass Spectrum for **Nmet8** (m/z 744.4879 for $[M+H]^+$ found 744.4882)

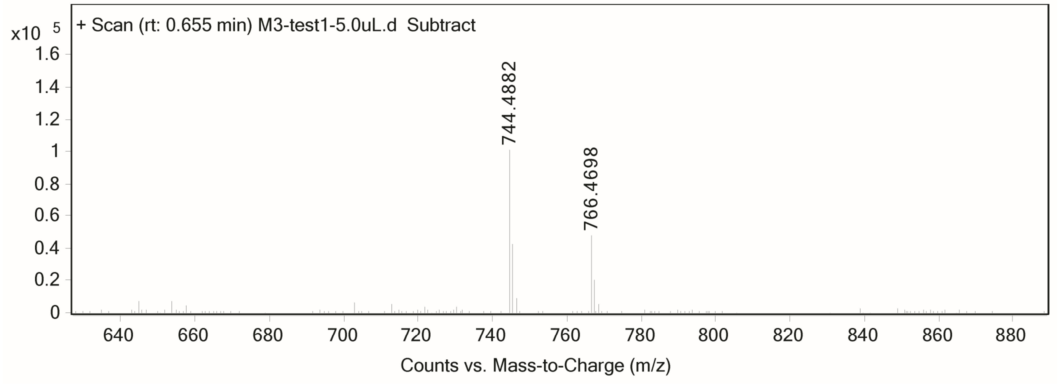

**Nmet9**

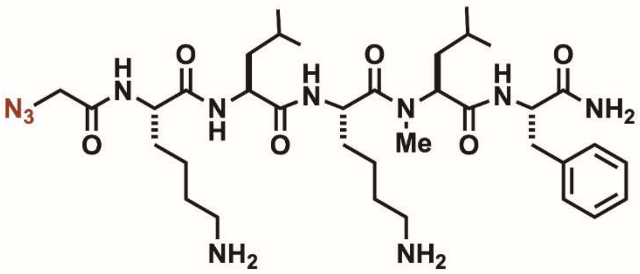


Analytical HPLC Chromatogram of **Nmet9**.

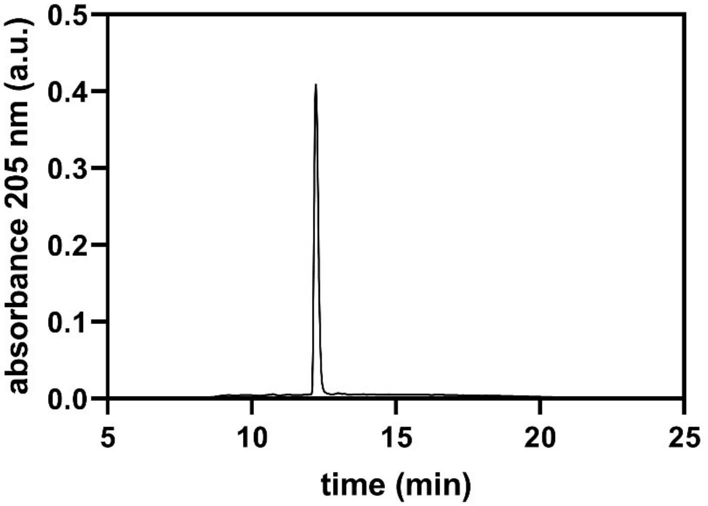


QTOF High Resolution Mass Spectrum for **Nmet9** (m/z 744.4879 for $[M+H]^+$ found 744.4875)

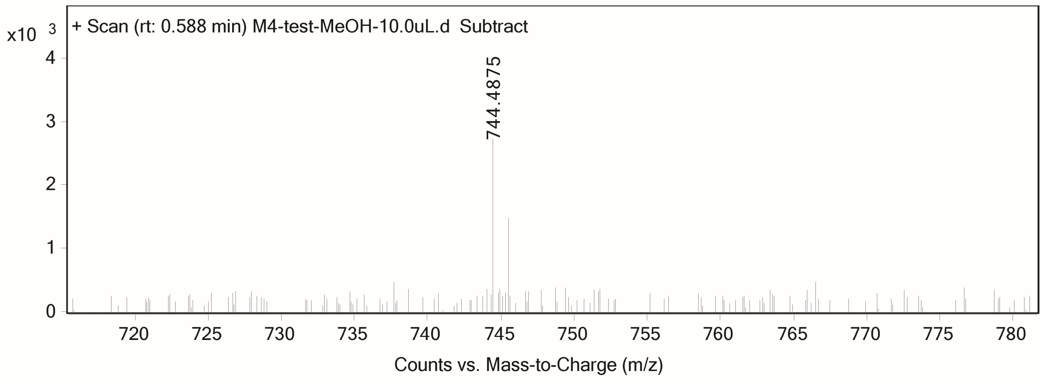

**Nalk1**

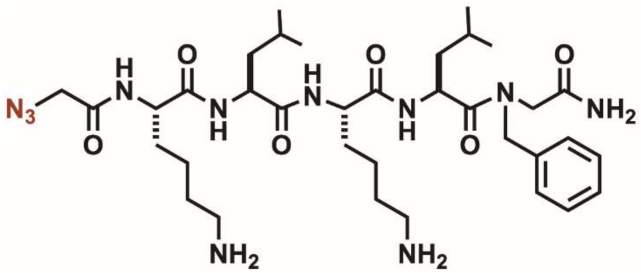

Analytical HPLC Chromatogram of **Nalk1**.

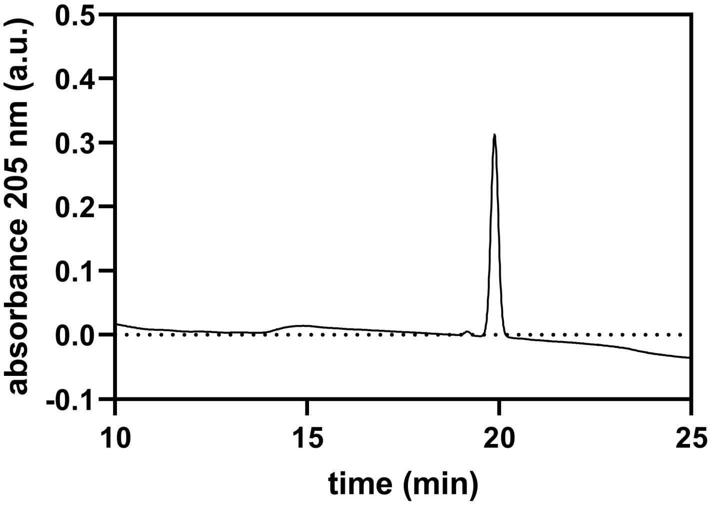


QTOF High Resolution Mass Spectrum for **Nalk1** (m/z 730.4723 for $[M+H]^+$ found 730.4732)

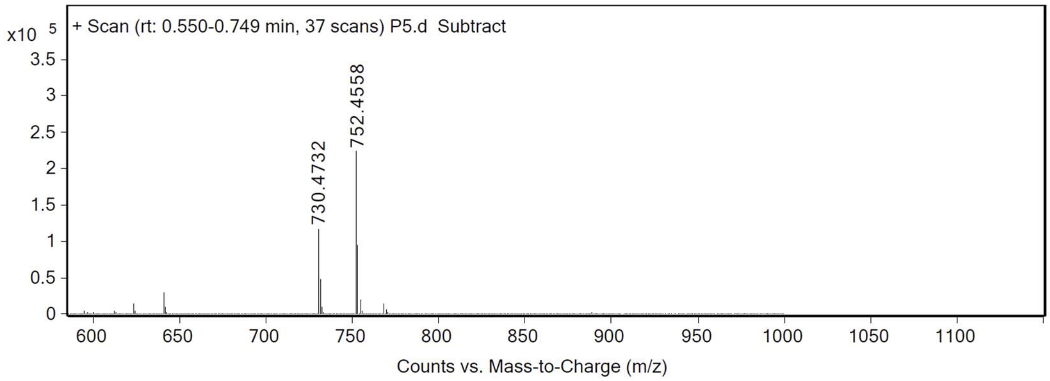

**Nalk2**

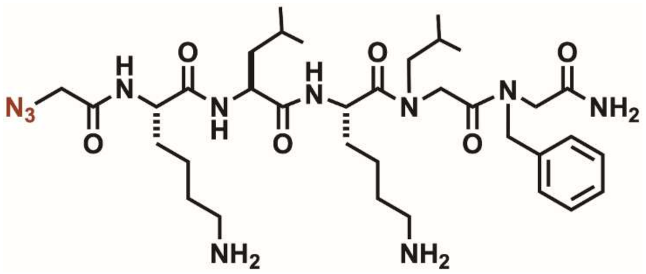


Analytical HPLC Chromatogram of **Nalk2**.

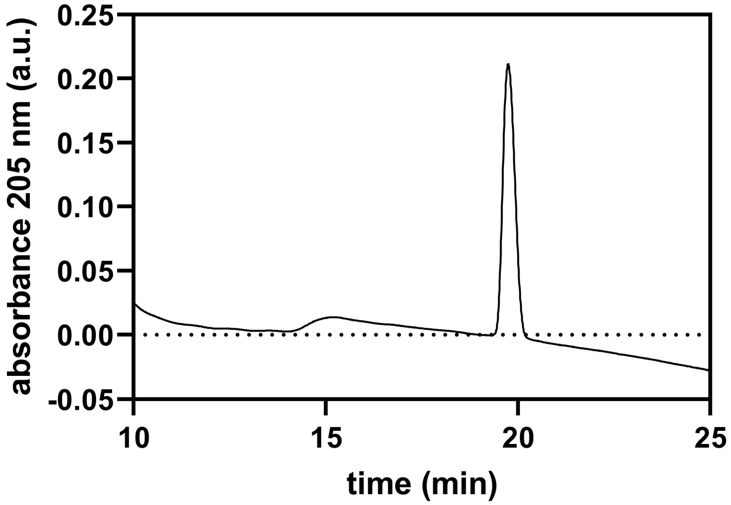


QTOF High Resolution Mass Spectrum for **Nalk2** (m/z 730.4723 for $[M+H]^+$ found 730.4732)

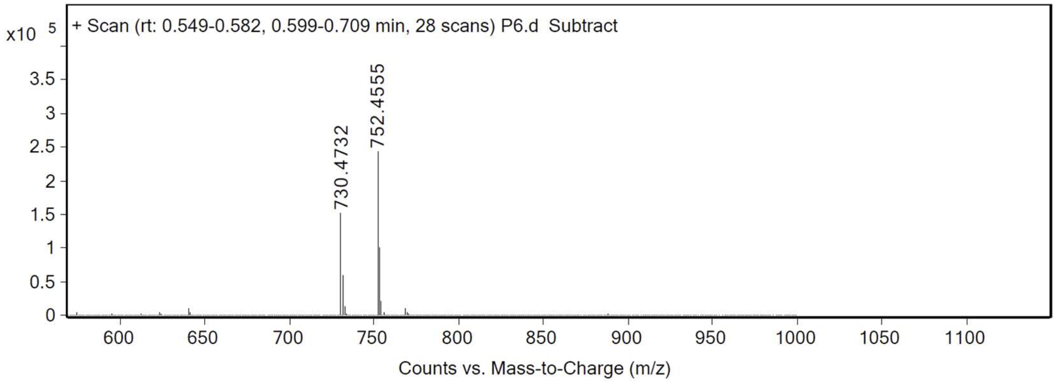

**Nalk3**

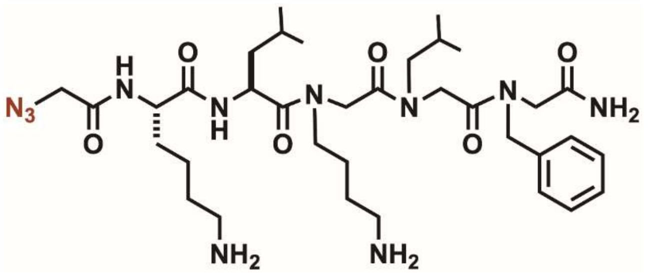


Analytical HPLC Chromatogram of **Nalk3**.

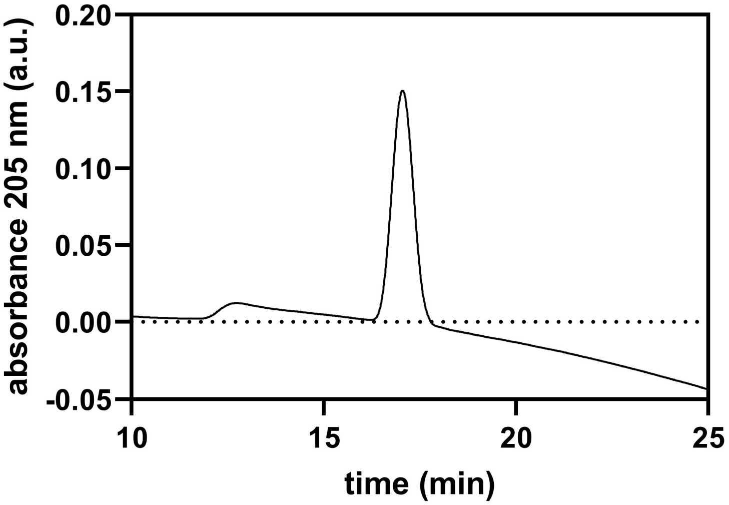


QTOF High Resolution Mass Spectrum for **Nalk3** (m/z 730.4723 for $[M+H]^+$ found 730.4732)

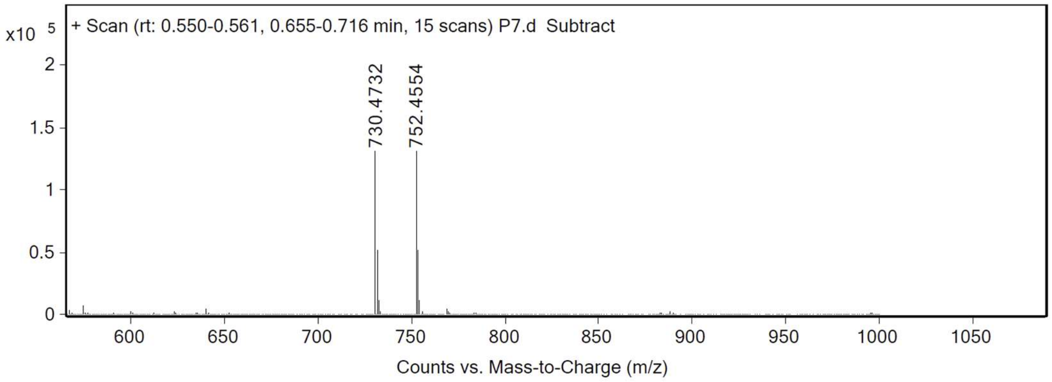

**Nalk4**

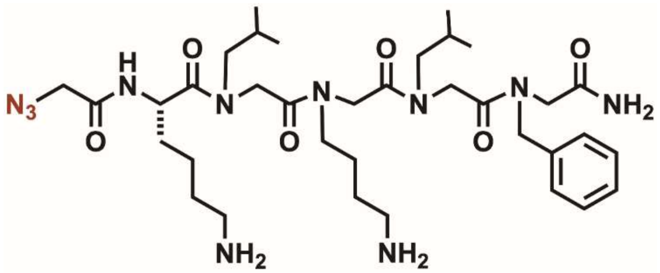

Analytical HPLC Chromatogram of **Nalk4**

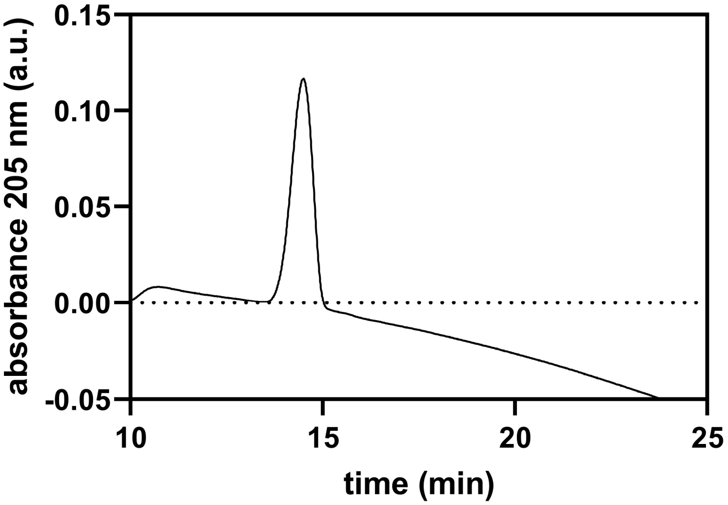


QTOF High Resolution Mass Spectrum for **Nalk4** (m/z 730.4723 for $[M+H]^+$ found 730.4730)

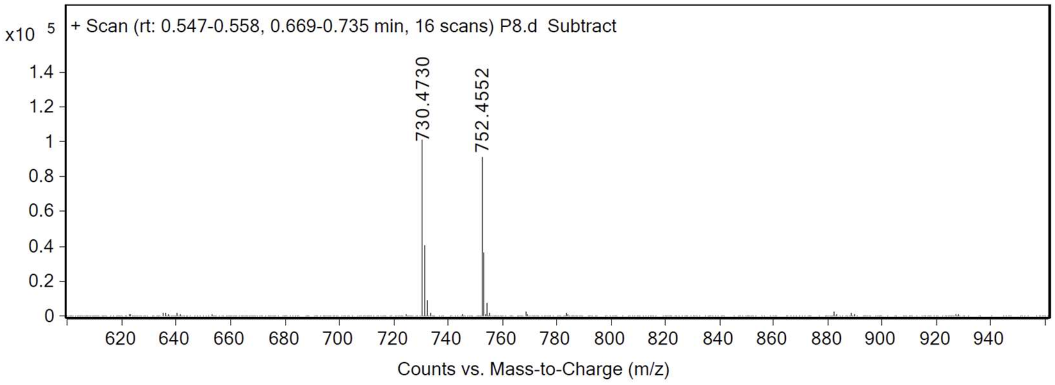

**Nalk5**

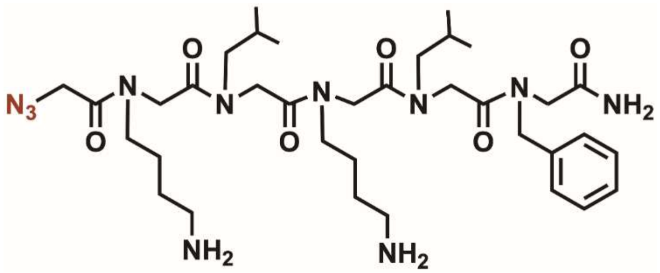


Analytical HPLC Chromatogram of **Nalk5**.

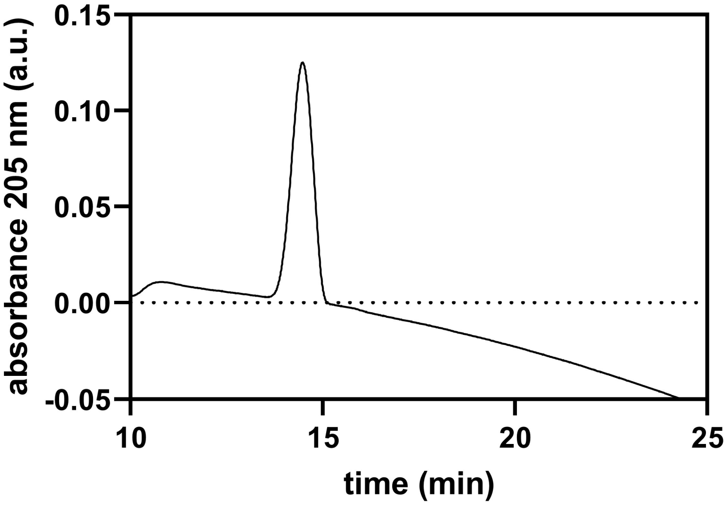


QTOF High Resolution Mass Spectrum for **Nalk5** (m/z 730.4723 for $[M+H]^+$ found 730.4730)

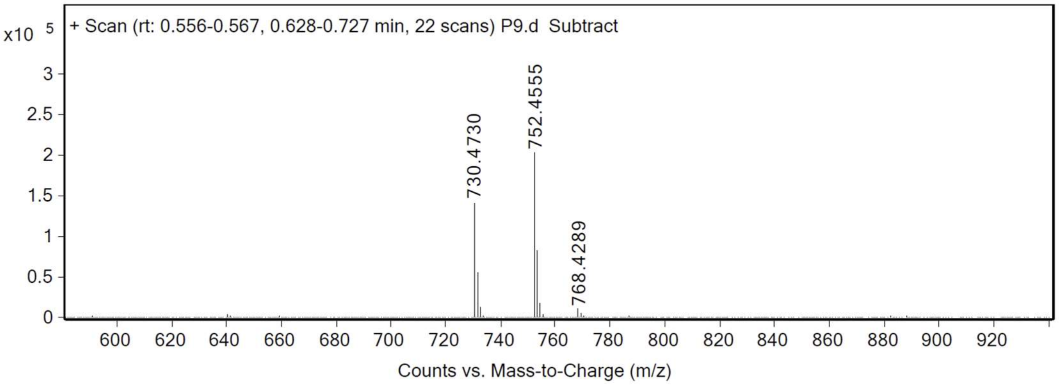

**Nalk6**

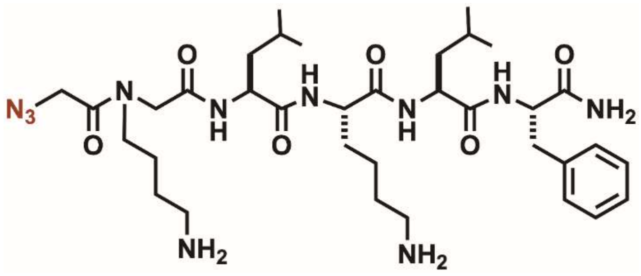


Analytical HPLC Chromatogram of **Nalk6**.

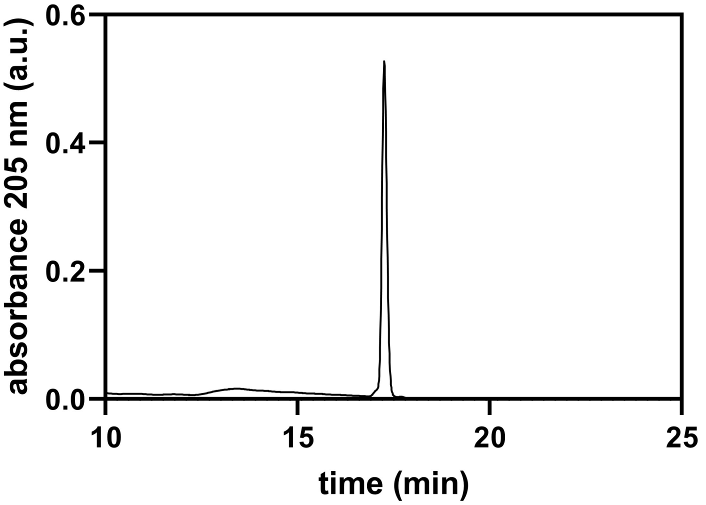


QTOF High Resolution Mass Spectrum for **Nalk6** (m/z 730.4723 for $[M+H]^+$ found 730.4729)

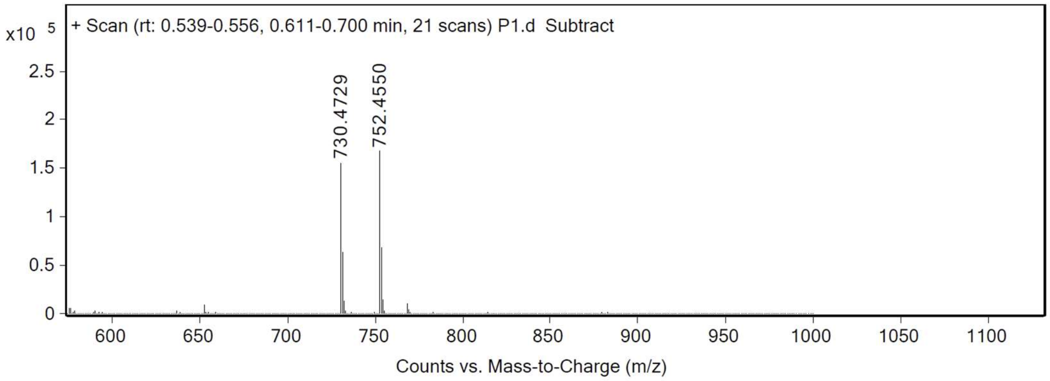

**NaIk7**

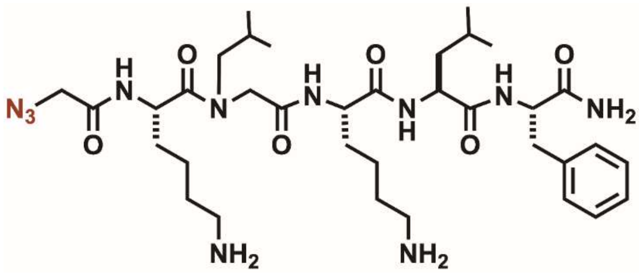


Analytical HPLC Chromatogram of **NaIk7**.

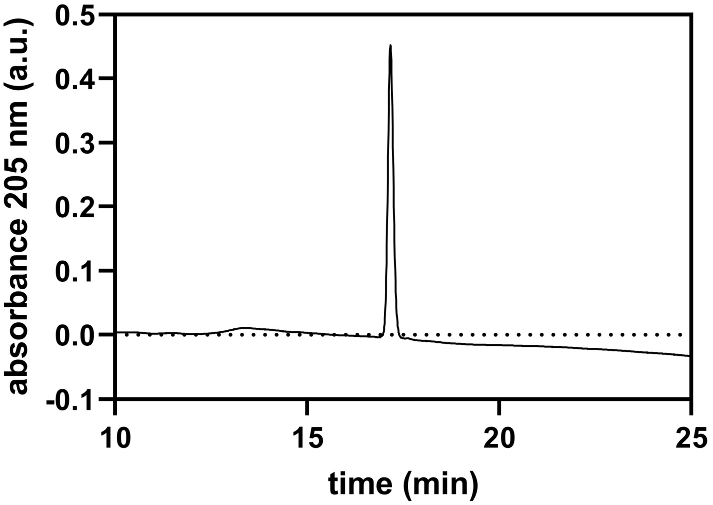


QTOF High Resolution Mass Spectrum for **NaIk7** (m/z 730.4723 for $[M+H]^+$ found 730.4729)

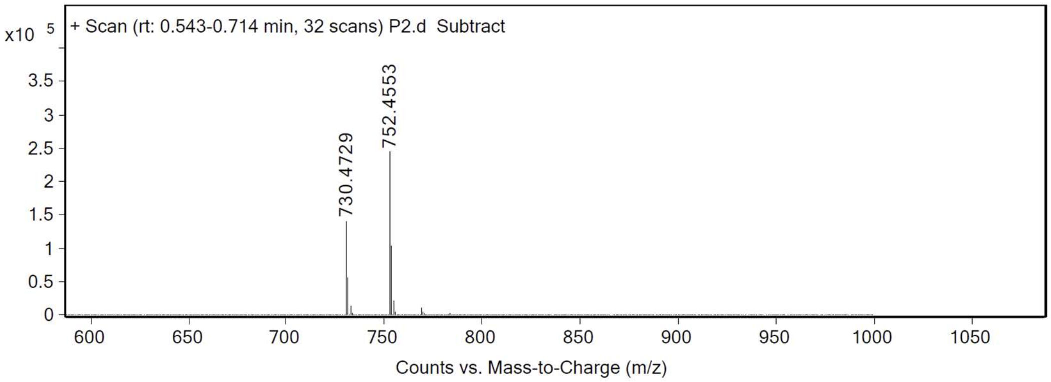

**Nalk8**

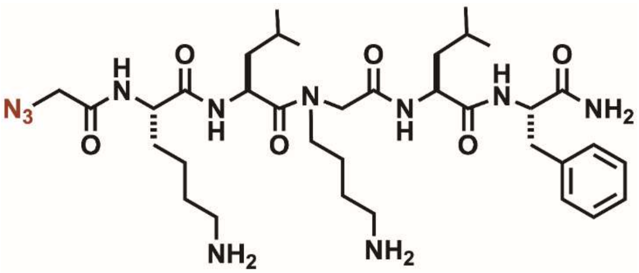


Analytical HPLC Chromatogram of **Nalk8.**

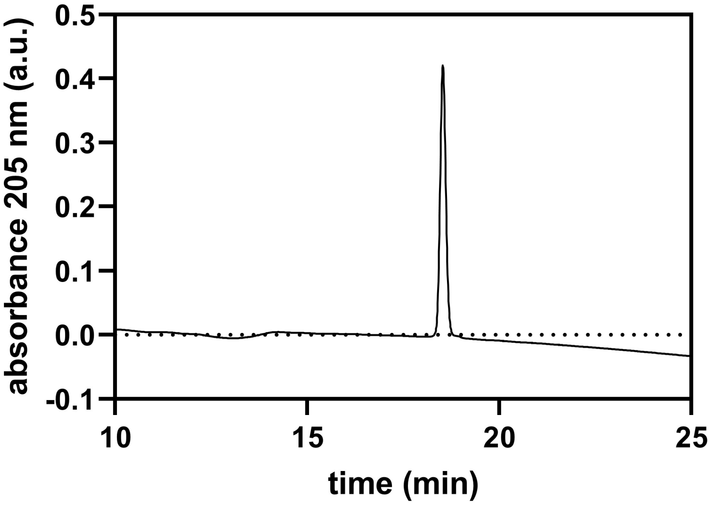


QTOF High Resolution Mass Spectrum for **Nalk8** (m/z 730.4723 for $[M+H]^+$ found 730.4727)

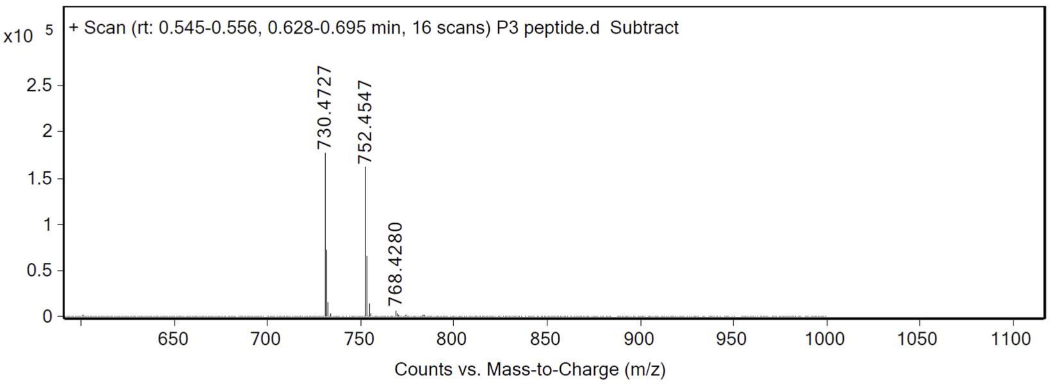

**Nalk9**

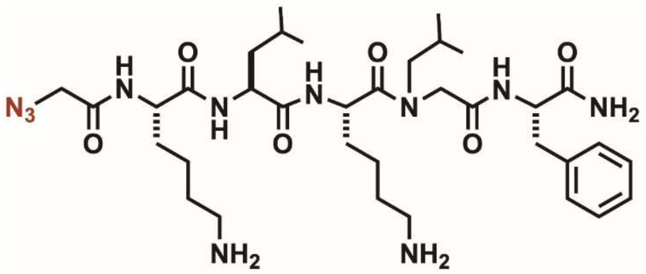


Analytical HPLC Chromatogram of **Nalk9**.

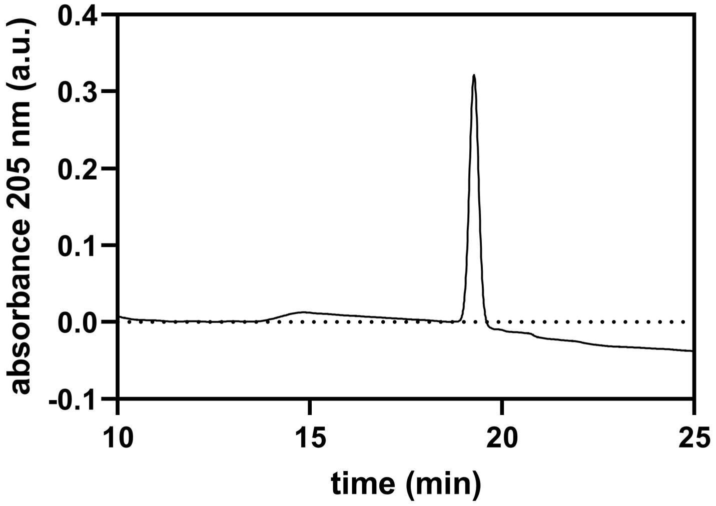


QTOF High Resolution Mass Spectrum for **Nalk9** (m/z 730.4723 for $[M+H]^+$ found 730.4729)

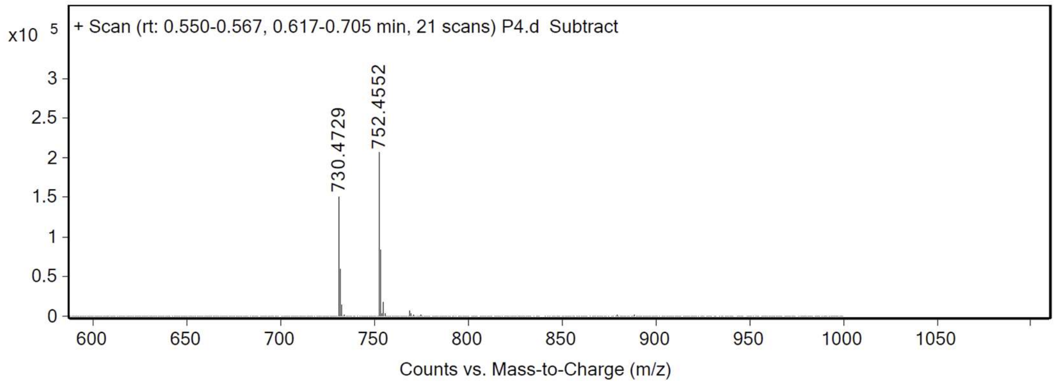

**tri**

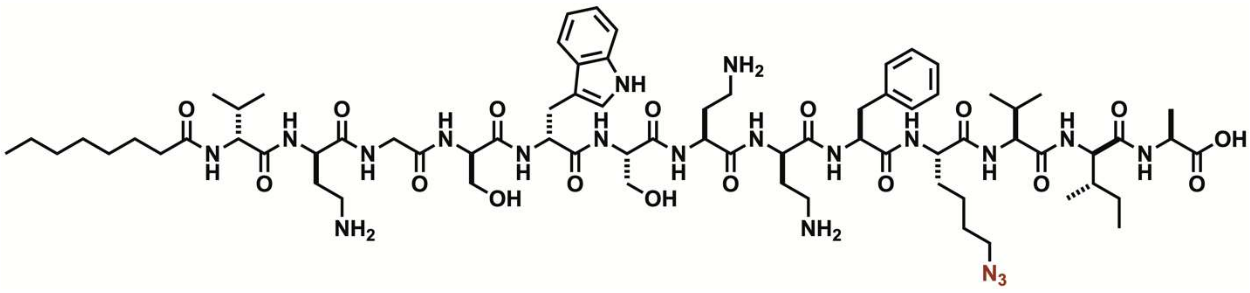


Analytical HPLC Chromatogram of **tri**.

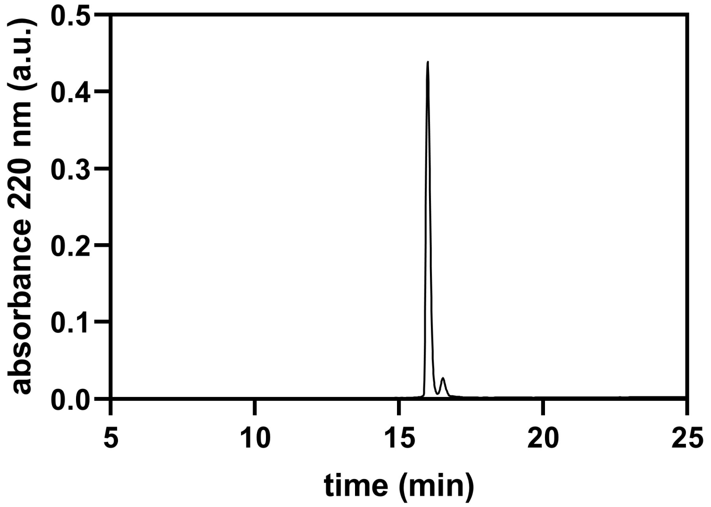


QTOF High Resolution Mass Spectrum for **tri** (m/z 1545.8900 for $[M+H]^+$ found 1545.8900)

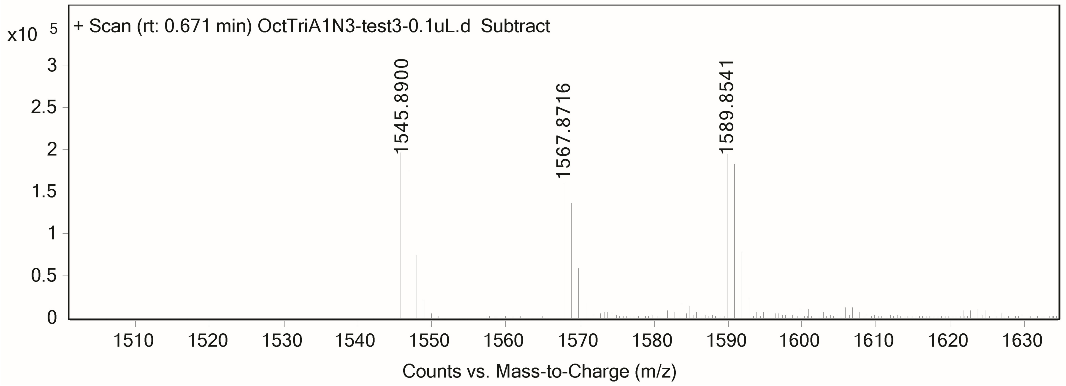

**triMe1**

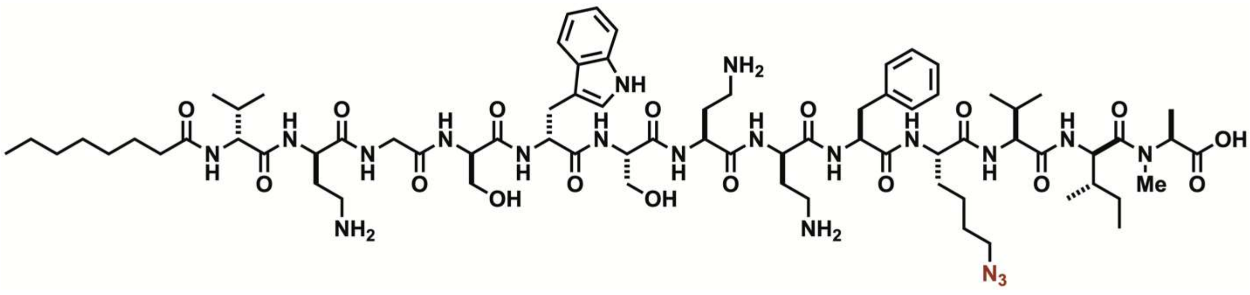


Analytical HPLC Chromatogram of **triMe1**

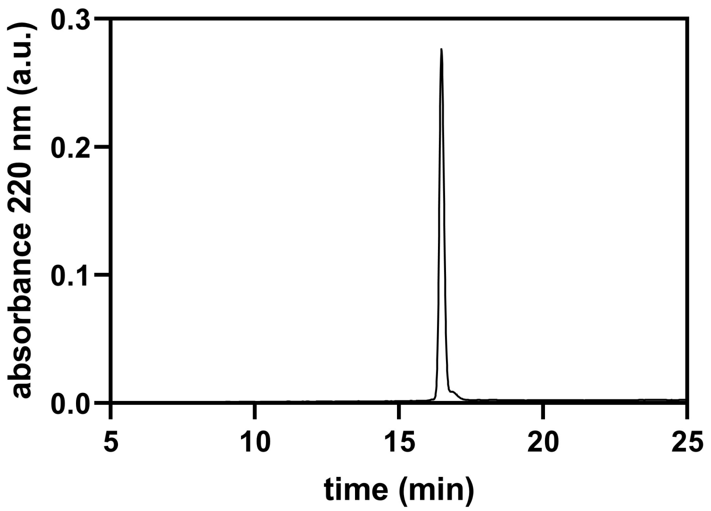


QTOF High Resolution Mass Spectrum for **triMe1** (m/z 1559.9057 for $[M+H]^+$ found 1559.9060)

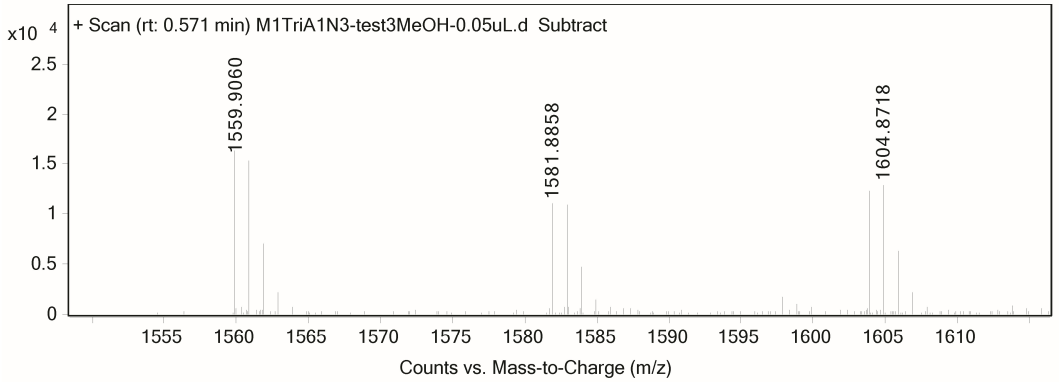

**triMe2**

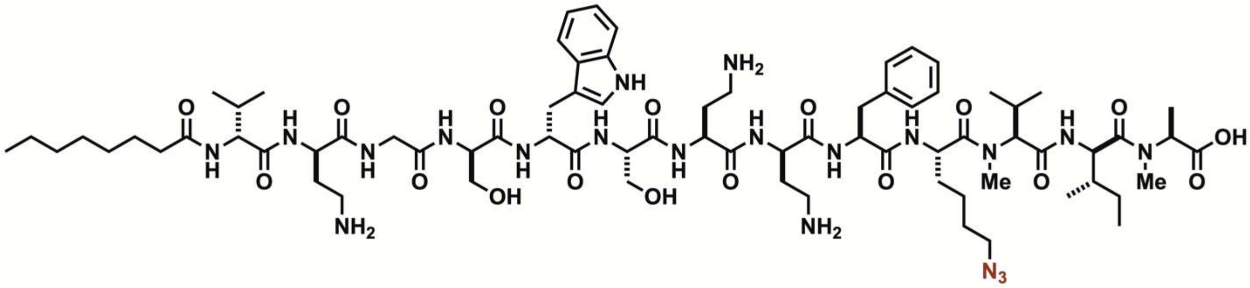


Analytical HPLC Chromatogram of **triMe2**

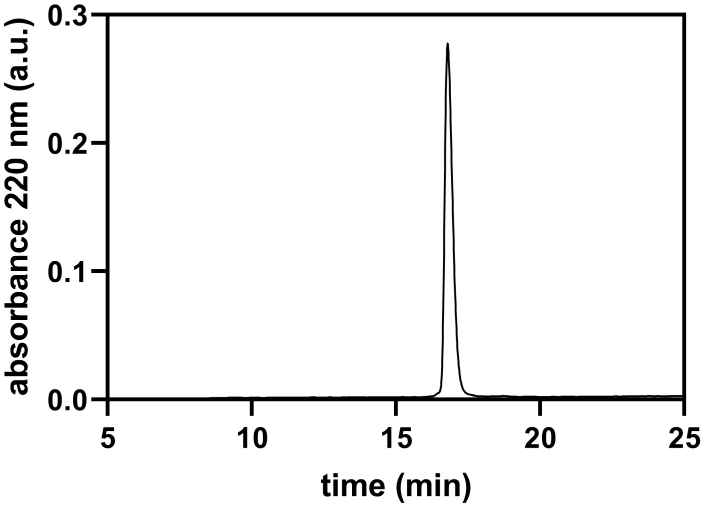


QTOF High Resolution Mass Spectrum for **triMe2** (m/z 1573.9213 for $[M+H]^+$ found 1573.9219)

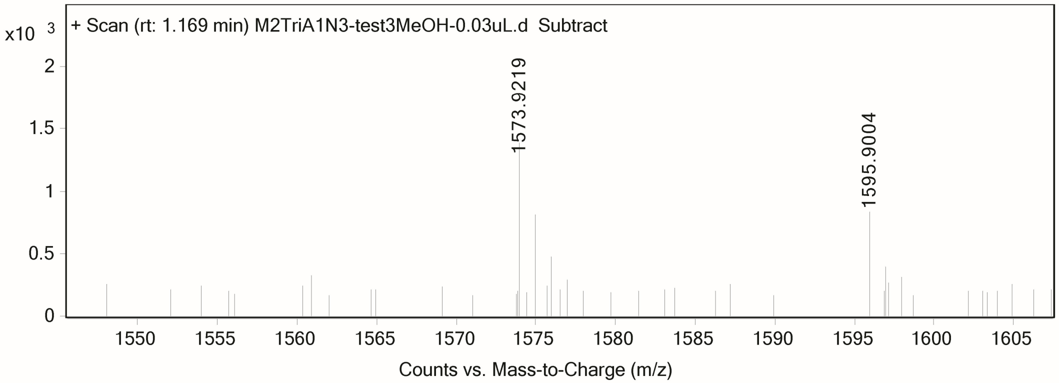

**triMe3**

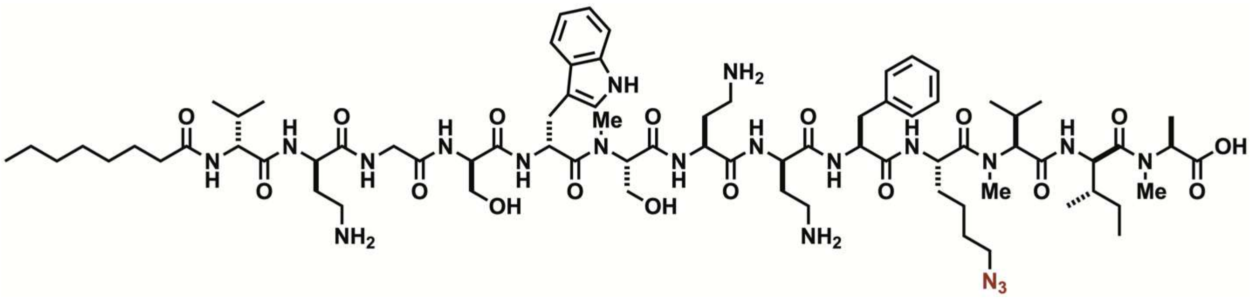


Analytical HPLC Chromatogram of **triMe3**

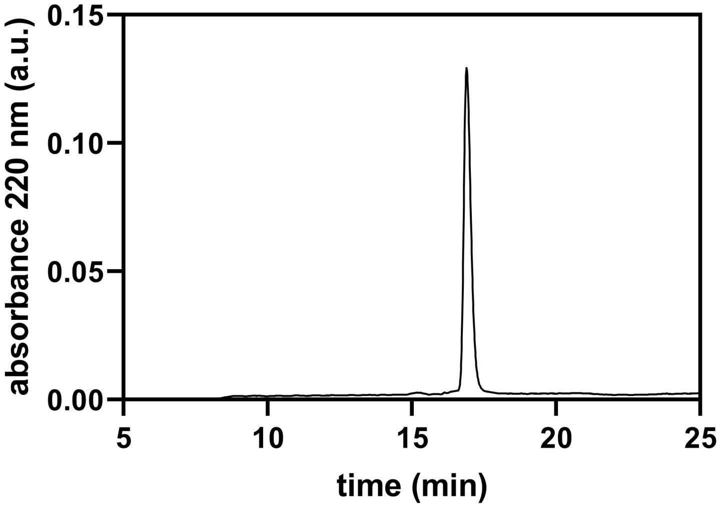


QTOF High Resolution Mass Spectrum for **triMe3** (m/z 1587.9370 for $[M+H]^+$ found 1587.9371)

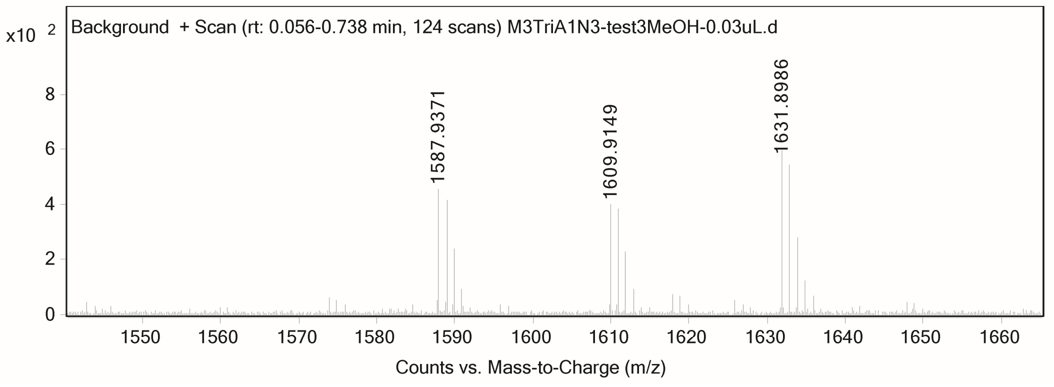

**triCyc**

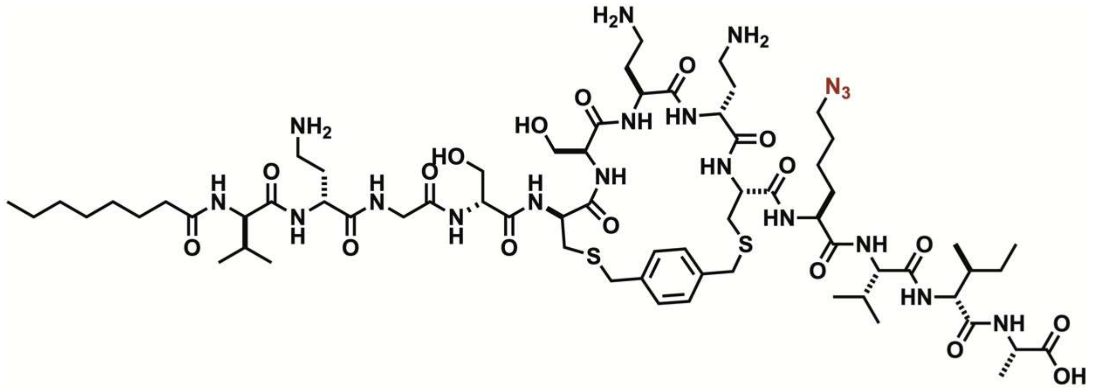

Analytical HPLC Chromatogram of **triCyc**.

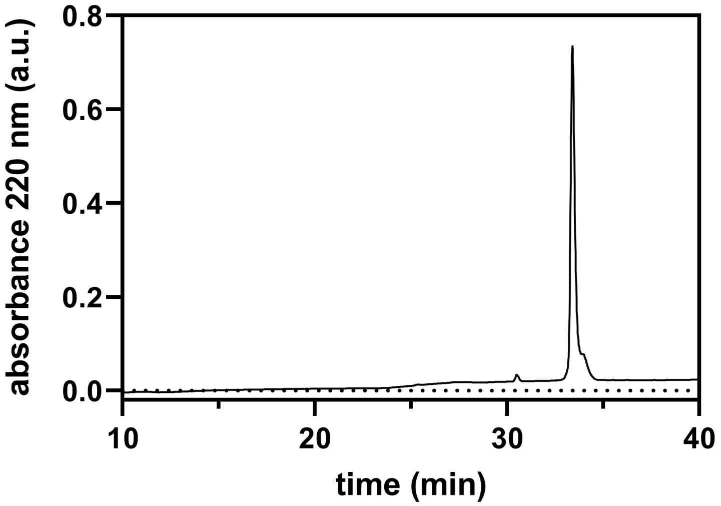


QTOF High Resolution Mass Spectrum for **triCyc** (m/z 1520.8003 for $[M+H]^+$ found 1520.8104)

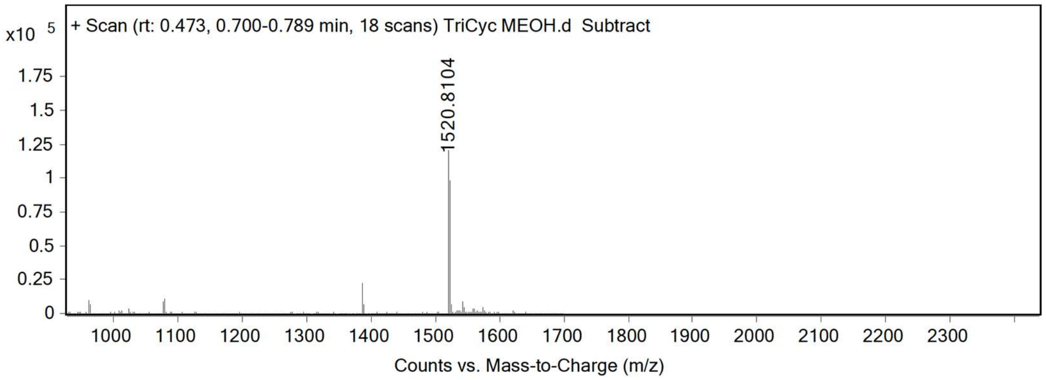

**triLin**

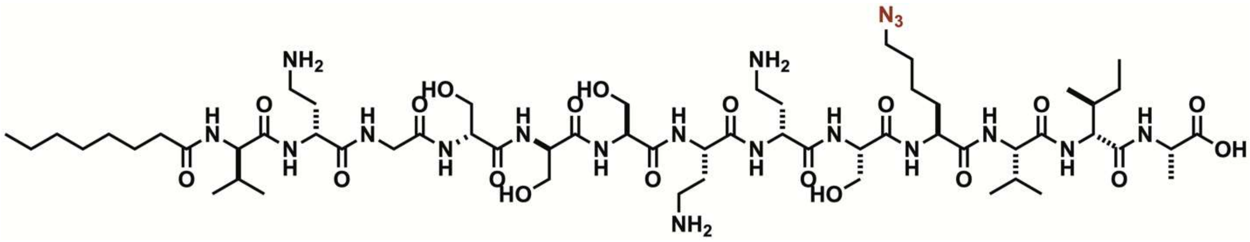

Analytical HPLC Chromatogram of **triLin**.

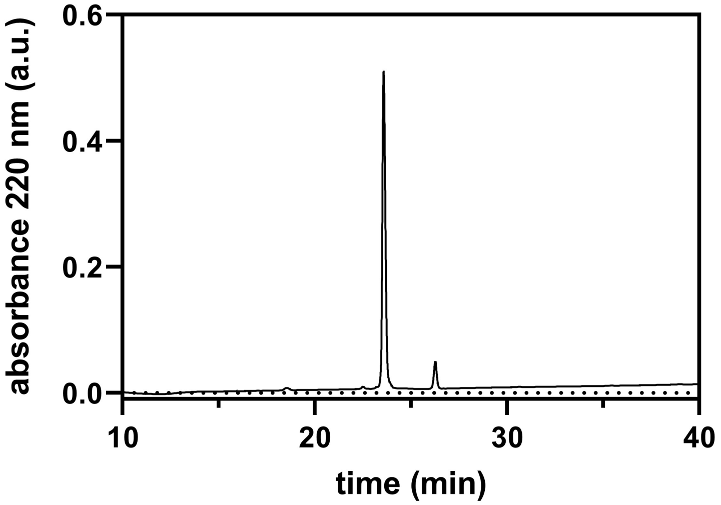


QTOF High Resolution Mass Spectrum for **triLin** (m/z 1386.7991 for $[M+H]^+$ found 1386.8347)

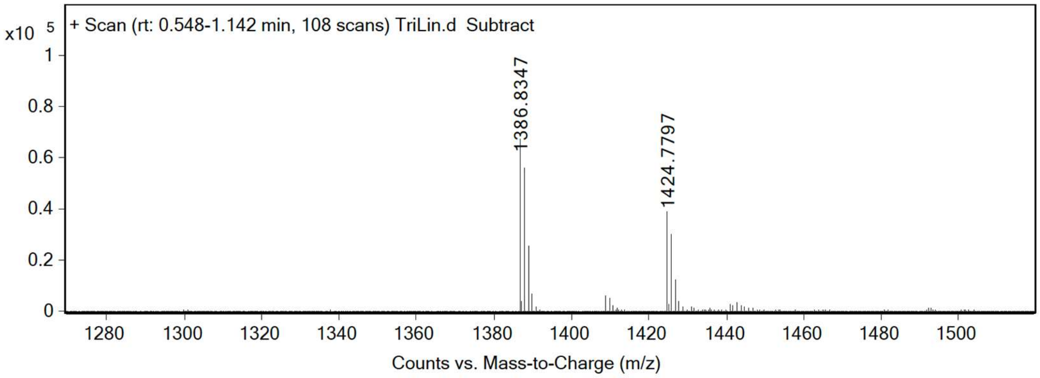

**grisMeCyc**

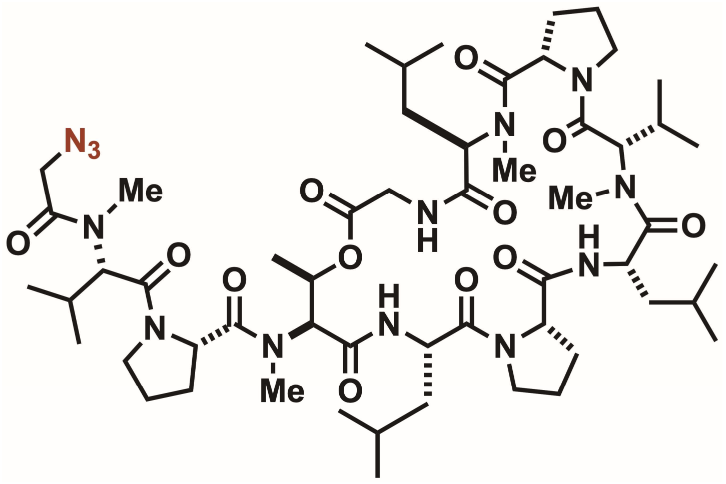


Analytical HPLC Chromatogram of **grisMeCyc**.

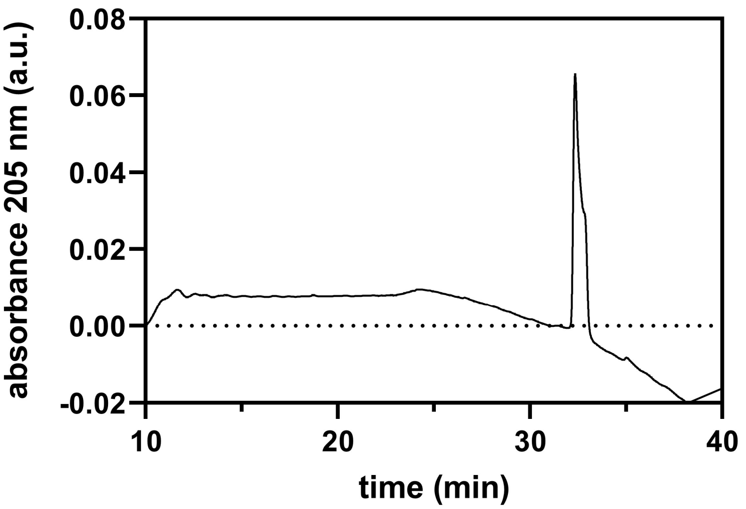


QTOF High Resolution Mass Spectrum for **grisMeCyc** (m/z 1126.6983 for $[M+H]^+$ found 1126.6987)

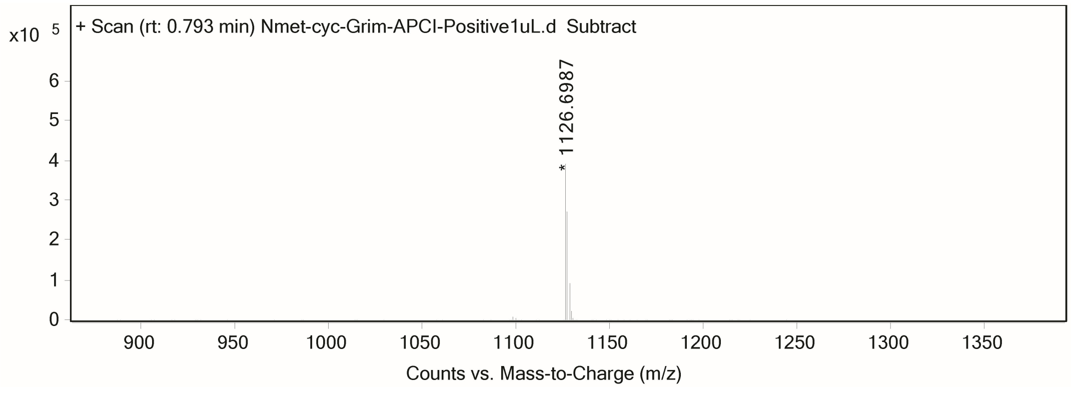

**grisCyc**

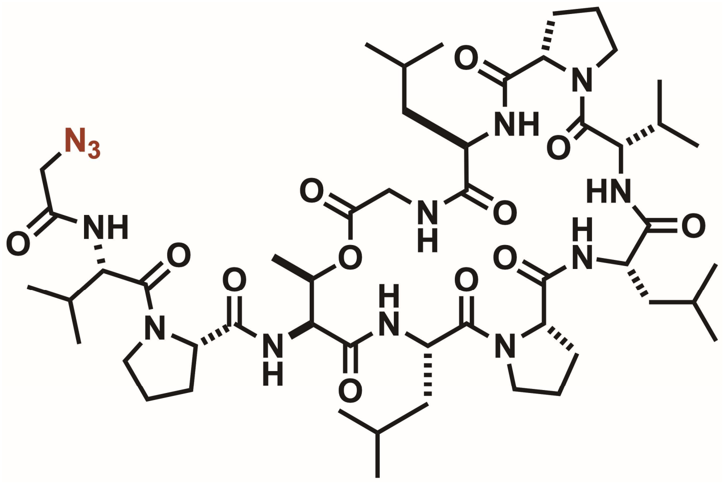

Analytical HPLC Chromatogram of **grisCyc**

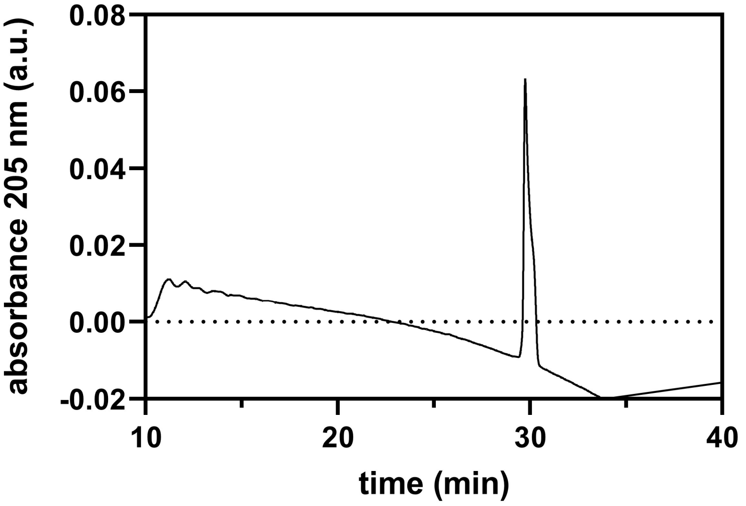


QTOF High Resolution Mass Spectrum for **grisCyc** (m/z 1070.6357 for $[M+H]^+$ found 1070.6355)

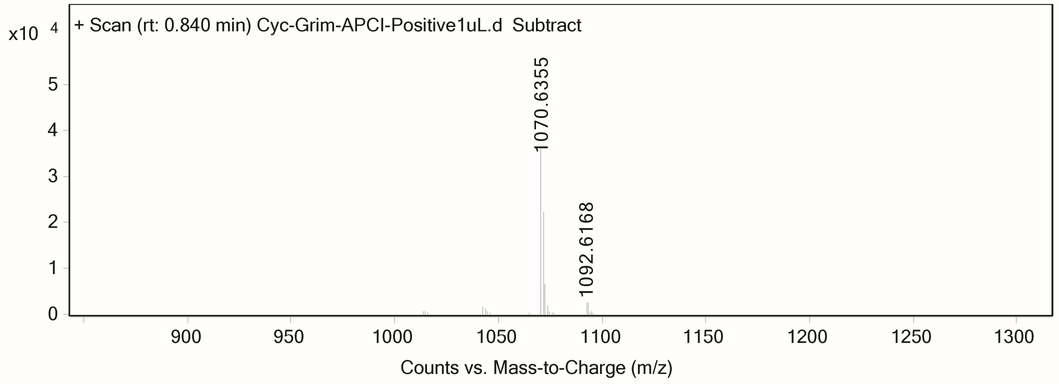

**grisMeLin**

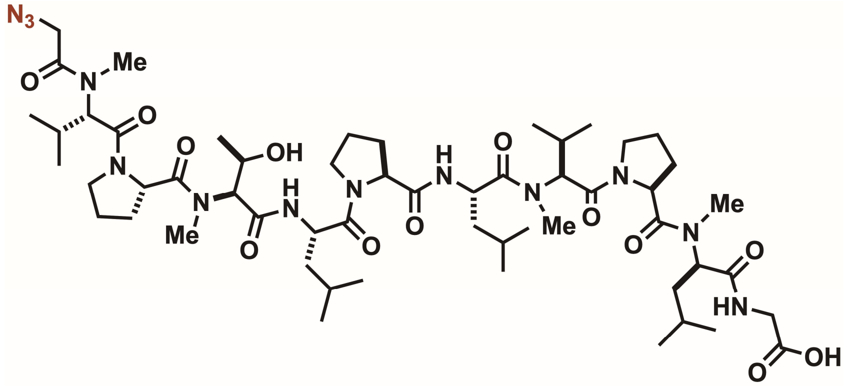


Analytical HPLC Chromatogram of **grisMeLin**

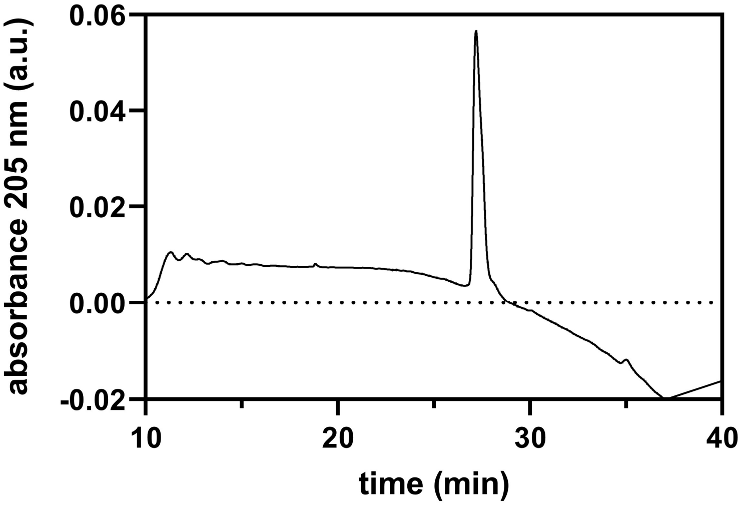


QTOF High Resolution Mass Spectrum for **grisMeLin** (m/z 1143.7248 for $[M+H]^+$ found 1143.7244)

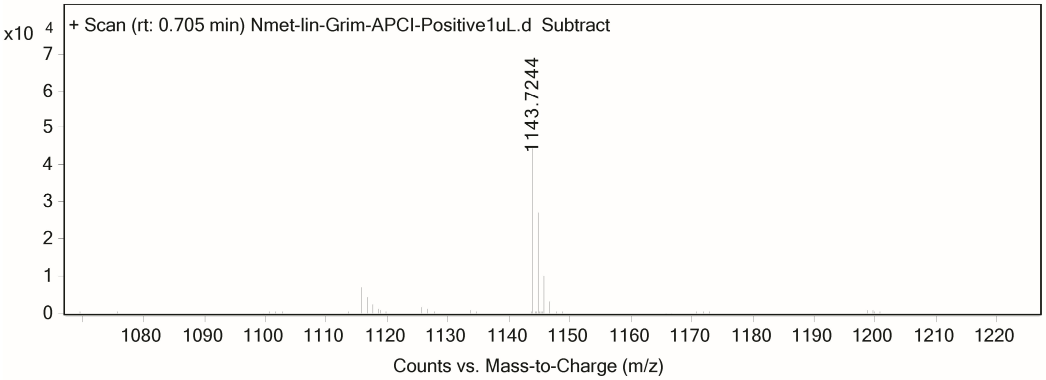

**grisLin**

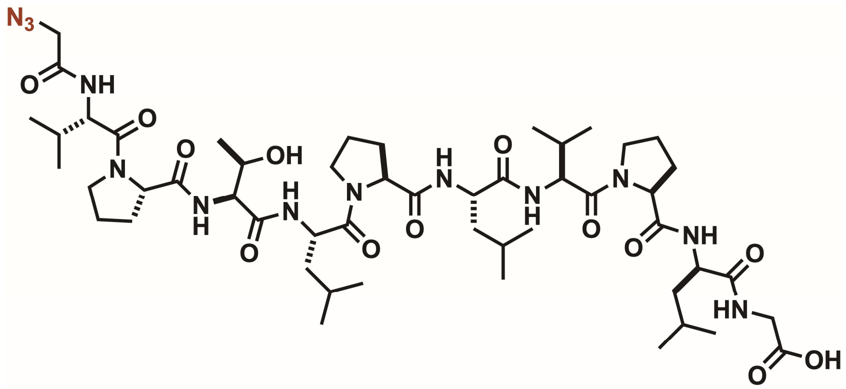

Analytical HPLC Chromatogram of **grisLin**.

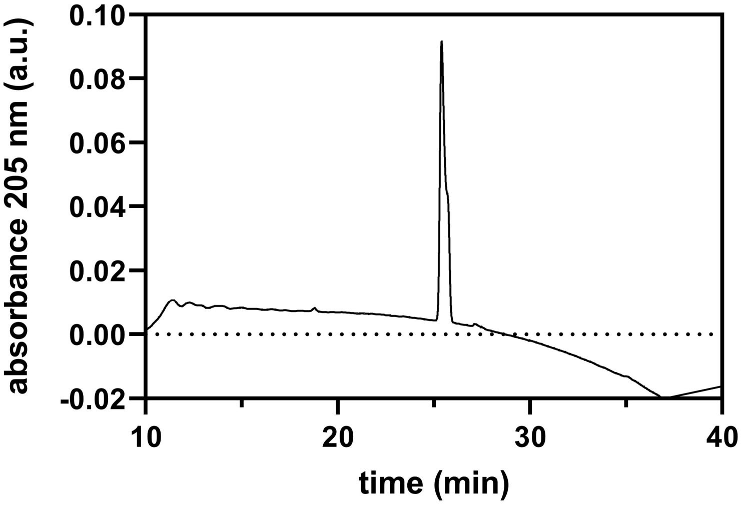


QTOF High Resolution Mass Spectrum for **grisLin** (m/z 1086.6622 for $[M+H]^+$ found 1087.6622)

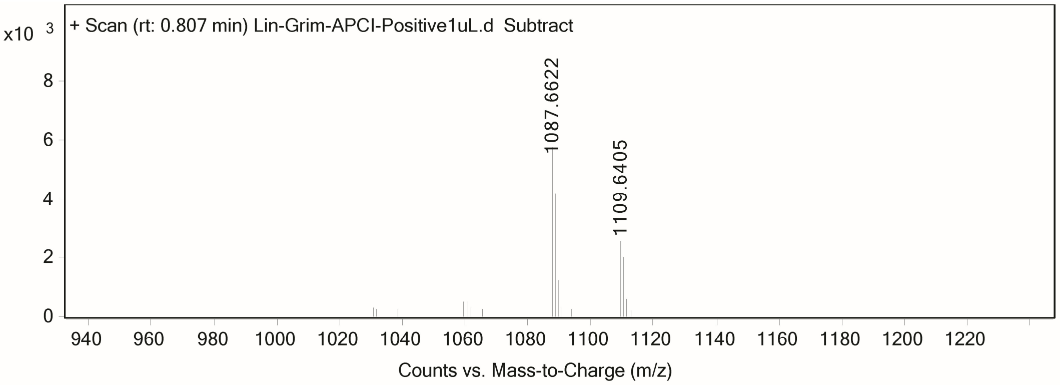
